# Genome-Wide Analysis of *TCP* Family Genes and Their Constitutive Expression Pattern Analysis in the Melon (*Cucumis melo*)

**DOI:** 10.1101/2024.07.30.605410

**Authors:** Md Jahid Hasan Jone, Md Nure Adil Siddique, Manosh Kumar Biswas, Mohammad Rashed Hossain

## Abstract

*TCP* proteins are plant-specific transcription factors that play essential roles in various developmental processes, including leaf morphogenesis and senescence, flowering, lateral branching, hormone crosstalk, and stress responses. However, the specific functions of *TCP* genes in melon remain largely unknown. This study identified and characterized 29 putative *TCP* genes in melon. These genes were classified into two classes: Class-I (13 genes) and Class-II (16 genes). The chromosomal location, gene structure, conserved motifs, structural homology, cis-regulating elements, transcript expression patterns, and potential protein-protein interactions were further analyzed. The results revealed that the putative *CmTCP* genes are distributed across nine of the twelve melon chromosomes and exhibit diverse expression patterns in different tissues and during floral organ development. Phylogenetic analysis suggests that some *CmTCP* genes may have similar functions to their homologs in other plant species, while others may have undergone functional diversification. This study provides a valuable resource for future investigations into the specific roles of individual *CmTCP* genes in melon development and paves the way for elucidating the mechanisms by which *TCP* proteins regulate leaf elongation, floral development, and lateral branching.

## INTRODUCTION

Melon (*Cucumis melo* L., 2n = 24) is an important economic crop grown worldwide for its sweet, edible and fleshy fruit. It is herbaceous, diploid and monoecious or andro-monoecious in nature. It produces male flowers in the main stem while lateral branches produce female or hermaphroditic flowers (Fukino et al. 2012). Well-maintained lateral branches result in good-quality fruits, higher yields and reduced cost of production as it is vital for light penetration, photosynthesis and energy partitioning. Lateral branching plays a significant role in the architecture of the melon plant (Fukino et al. 2012; Shahwar et al. 2023). Breaking the dominance of apical meristem through main stem pruning increases the lateral branching and leaf expansion, positively affecting fruit quality (Pereira et al. 2003; Widaryanto et al. 2020). However, the transcription factors responsible for lateral branching in melons were rarely studied before.

The *TCP* family genes are a group of plant-specific (Michael and Pilar 2016; Navaud et al. 2007) transcription factors encoding genes named after *TEOSINTE BRANCHED 1* (*TB1)*, *CYCLOIDEA* (*CYC*) and *PCF* proteins initially found in Maize, Garden snapdragon and Rice, respectively (Cubas et al. 1999; Finlayson 2007; Kosugi and Ohashi 2002). The conserved region of *TCP* is predicted to form a basic helix-loop-helix (bHLH) structure of 59 amino acid residues (Cubas et al. 1999). Based on sequence homology, *TCP* proteins are divided into two subfamilies namely, Class-I (conserved four-amino-acid deletion in basic region) and Class-II (Manassero et al. 2013). Class-I subfamily contains *PCF* (*TCP*-*P*) proteins whereas Class-II subfamily is again divided into two clades: *CIN* and *CYC/TB1* (Manassero et al. 2013).

The *TCP* proteins that have been characterized thus far function in meristem growth: TB1 influences the growth of axillary meristems, while *CYC* regulates the growth of floral meristems and primordia. *PCF* bind to the promoter of a gene involved in meristematic cell division (Cubas et al. 1999). A number of *CYC/TB1* genes are found to be involved in the growth of auxiliary meristems and buds, which produce lateral shoots (Michael and Pilar 2016).

The functions of *TCP* proteins in different plants were extensively studied. Members of the *TCP* family were found to be involved in the morphogenesis of shoot lateral organs (Aguilar-Martinez et al. 2007; Finlayson 2007), shoot branching (Martin-Trillo et al. 2011), cell growth and proliferation in leaf (Aguilar-Martinez and Sinha 2013; Kosugi and Ohashi 1997), hormone signalling (Daviere et al. 2014; Li and Zachgo 2013), controlling apical dominance (Doebley et al. 1997), leaf and floral development (Aguilar-Martinez and Sinha 2013). Overall, *TCP* proteins affect leaf shape and induce branching activity in plants (Michael and Pilar 2016).

Despite the importance of lateral branching in melon and its predictable relation with *TCP* family genes, a comprehensive genome-wide analysis of these transcription factors is yet to be conducted. The present study aimed to identify genome-wide *TCP* family genes and to analyze their structure, motif distribution, evolution and syntenic relation with other crops, to predict protein structure (homology modelling), sequence similarity, cis-regulatory elements, and to analyze protein-protein interaction and expression profiling. The findings may provide a deeper insight into the abundance, and evolutionary history of this gene family and may reveal a set of candidate genes responsible for lateral branching in melon.

## MATERIALS AND METHODS

### Sequence Retrieval and Confirmation

A comprehensive search of five databases, namely PlantTFDB (https://planttfdb.gao-lab.org/), iTAK (http://itak.feilab.net/), Melonomics v4.0 (https://www.melonomics.net/), CuGenDB (http://cucurbitgenomics.org/) and CuGenDBv2 (http://cucurbitgenomics.org/v2/), was conducted to identify melon *TCP* family genes and their corresponding proteins (Jin et al. 2017; Yu et al. 2022; Zheng et al. 2016). Consolidating the results from all five databases unique gene IDs for all *TCP* genes were recorded. To confirm the presence of *TCP* and other potential domains, these sequences were analyzed using NCBI Conserved Domain (www.ncbi.nlm.nih.gov/Structure/cdd/wrpsb.cgi) Search and GenomeNet Motif Search (www.genome.jp/tools/motif/). Following this analysis, sequences containing the *TCP* domain were selected for further investigation Genomic information and protein sequences for *TCP* genes in Cucumber and Tomato were retrieved from published articles (Parapunova et al. 2014; Wen et al. 2020). Corresponding genome databases (www.arabidopsis.org/, https://rapdb.dna.affrc.go.jp/) and published articles were used to retrieve data for Arabidopsis and Rice (Shang et al. 2022; Su et al. 2021; Yao et al. 2007).

### Characteristics of *CmTCP* Proteins

Protein length (aa), Molecular weight (KDa), Isoelectric points (pI), and Grand Average of Hydropathy (GRAVY) values were determined from Expasy-Protparam (Gasteiger et al. 2003). Subcellular localizations were predicted by CELLO: Subcellular Localization Predictive System (Yu et al. 2006).

### Chromosomal Localization and Gene Duplication Analysis

Chromosome lengths and chromosomal positions of *CmTCP* genes were retrieved from the CuGenDBv2 database (Yu et al. 2022). Genes were mapped on chromosomes using TBtools software (Chen et al. 2020a). Nine chromosomes that contain *CmTCP* gene(s) were visualized excluding the rest three. Duplicate gene pairs and the type of duplications were determined using MCScanX (Wang et al. 2012) and then visualized in TBtools software.

### Genomic Structures, Motif Distribution, and Cis-Acting Elements Analysis

GSDS_2.0_ web tools (Hu et al. 2015) were used to analyze the exon-intron structures of the *TCP* family genes. TBtools software (Chen et al. 2020a) and MEME web tools (Bailey et al. 2015) were used to analyze the motif distribution.

Upstream sequences of each *CmTCP* gene (2000 bp) were retrieved using PlantCARE (Lescot et al. 2002) and a heatmap was generated based on the numbers using MS Excel 2019. The Basic Biosequence view in TBtools was used to localize the cis-regulatory elements in the upstream region. The grouping of cis-regulating elements according to their functions was carried out as the procedures described in Wu et al (Wu et al. 2023).

### Sequence Alignment and Phylogenetic Analysis

BioEdit Sequence Alignment Editor (Alzohairy 2011) was used to align *CmTCP* protein sequences (Supplementary File 1) via ClustalW and visualized to find the *TCP* domain region. Molecular Evolutionary Genetics Analysis (MEGA11 and MEGA7) software (Kumar et al. 2016; Tamura et al. 2021) was also used to align the *TCP* sequences of melon again via ClustalW with default settings. A rectangular phylogenetic tree was constructed with the aligned *CmTCP* sequences, via MEGA11, using the Maximum Likelihood method and 1000 replicates for bootstrap. A circular phylogenetic tree was constructed with the aligned sequences of Melon, Cucumber, Rice, Tomato and Arabidopsis, via MEGA7, using the Maximum Likelihood method and 1000 replicates for bootstrap. Finally, the tree was visualized using Interactive Tree Of Life (iTOL) v5 (Letunic and Bork 2021).

### Synteny Relationships

Genomic annotation (.gff) and genome sequence (.fa) files of Cucumber and Arabidopsis were retrieved from their genome database (CuGenDBv2 and TAIR Database) and Phytozome (Goodstein et al. 2012) was used to retrieve the files of Rice, and Tomato. The syntenic relationship of *CmTCP* genes with these genomes was analyzed using the MCScanX tool (Wang et al. 2012). TBtools software was used to construct the collinearity map (Chen et al. 2020a).

### Expression Profiling Using RNA-Seq Data

Melonet DB (melonet-db.dna.affrc.go.jp/ap/mvw), a web tool for functional genomics research of melons (Yano et al. 2020) was used to analyze the expression of *CmTCP* genes based on the available RNA-seq data. FPKM values of 25 genes for 15 tissues and 45 different samples were available in the database. The tissues include-callus, seed (dry and imbibed), cotyledon, hypocotyl, root, flower (petal, anther, stigma, ovary), stem, shoot apex, leaves, tendril, fruit (flesh and epicarp). Heatmap of expression data was generated by R programming via the “pheatmap” package (Kolde 2019) using Euclidean distance and ward.D2 as a clustering method.

### Protein Structure Prediction and Sequence Similarity

Protein homology modeling was conducted using the Swiss Model (Waterhouse et al. 2018) to generate predicted three-dimensional structures based on template (known homologous protein structure) identification. Additionally, sequence similarity analysis was performed using TBtools, facilitating the comparison of amino acid sequences and the construction of a similarity matrix.

### Interaction Network of *CmTCP* Proteins

Protein-protein association networks of *CmTCP* proteins were predicted and visualized using the String Database tool (Szklarczyk et al. 2023). Three clusters of proteins were predicted using the K-means clustering method as well (Kalaivani et al. 2018).

## RESULT

### Identification of *TCP* genes in the Melon genome

A total of 29 *TCP* genes were identified in PlantTFDB and iTAK. Besides, 23 *TCP* genes were identified in Melonomics v4.0, while 50 and 47 genes were identified in CuGenDB and CuGenDBv2, respectively. These altogether yielded 53 unique gene IDs which were further analyzed based on Conserved Domain and Motif analyses. Finally, 29 sequences containing the *TCP* domain were selected for subsequent analysis (Supplementary Table 1). These 29 sequences were named *CmTCP-01 to CmTCP-29* according to their location in various chromosomes. The size of *CmTCP* proteins ranges from 61 to 600 amino acids with a molecular weight of 7.11516 to 65.4692 kDa (Table 1). The highest size and molecular weight were found in the case of *CmTCP-10* and the lowest was for *CmTCP-04*.

**Table 1.**
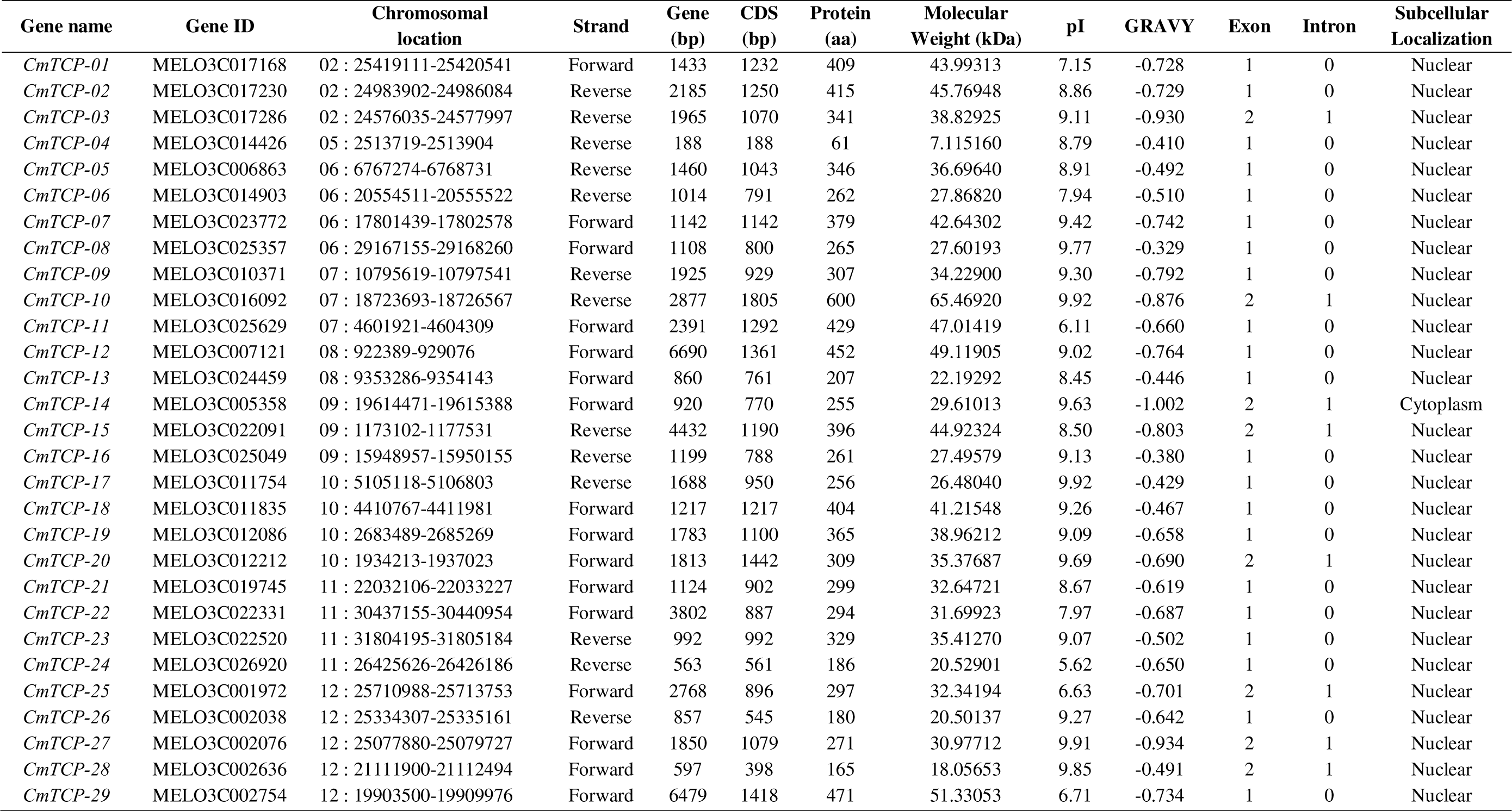
Characteristics of the 29 *CmTCP* family gene in melon.

The isoelectric point ranged from 5.62 to 9.92, where the pI value for 25 proteins was more than 7 indicating that *CmTCP* proteins are mostly basic in nature. Contrastingly, genes *CmTCP-11*, *CmTCP-24*, *CmTCP-25*, and *CmTCP-29* were found to be acidic in nature. Only one protein (*CmTCP-14*) is predicted to be localized at the cytoplasm whereas all other genes are predicted to be localized at the nucleus (Table 1). The GRAVY value for *CmTCP* protein ranges from -0.329 (*CmTCP*-08) to -1.002 (*CmTCP*-14). The negative value for every protein indicates that *CmTCP* proteins are non-polar and hydrophobic in nature (Table 1).

### Chromosomal localization

The chromosomal distribution of *CmTCP* family genes illustrates that they are distributed throughout the 9 (out of 12) chromosomes in the melon genome while chromosomes 1, 3 and 4 contain no *TCP* gene (Figure 1). The highest number (five) of *CmTCP* genes were found in chromosome 12. Chromosomes 6, 10, and 11 contain 4 *CmTCP* genes whereas chromosomes 2, 7, and 9 contain 3 *CmTCP* genes in each. Chromosomes 8 and 5 contain 2 and 1 genes respectively. Most of the genes are located far from the centromere with a tendency to be located toward the telomere regions. Only a few genes such as *CmTCP-06, CmTCP-07, and CmTCP-09* are located in the centromeric regions (Figure 1).

**Fig. 1.**
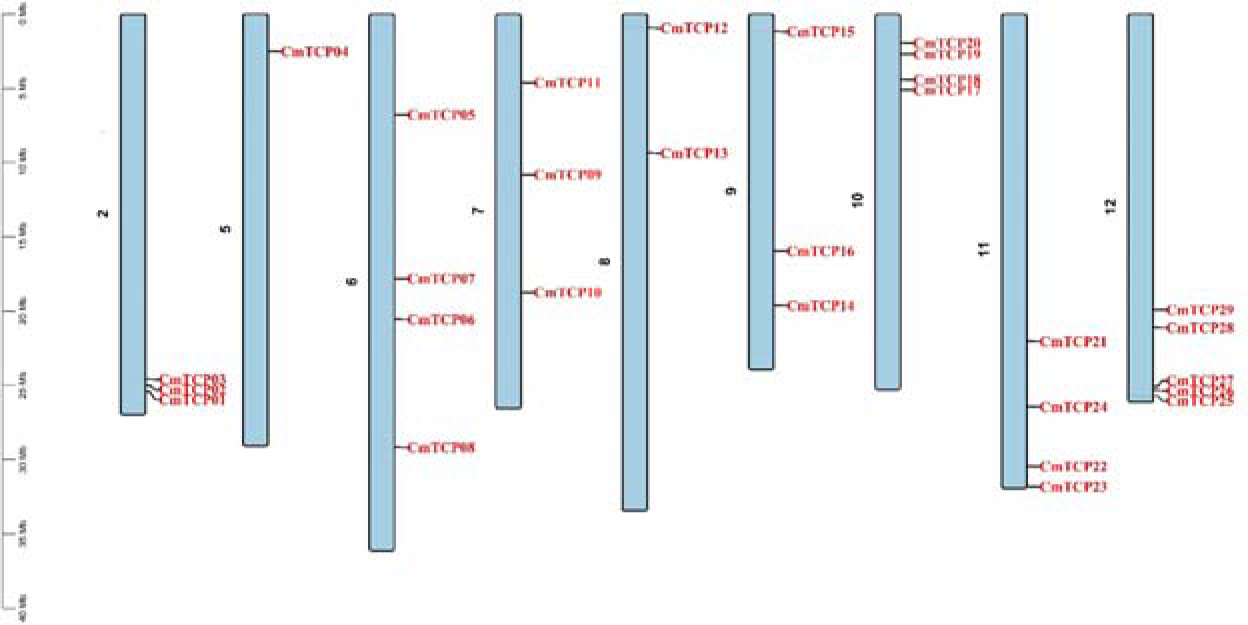
Chromosomal distribution of 29 *CmTCP* genes in melon. The length of the chromosomes is indicated by the scale on the left.

### Sequence Alignment and *TCP* Domain

The alignment of *CmTCP* proteins (Supplementary File 1) revealed the *TCP* domain with 59 amino acid residues consisting of a ‘basic’ region and a helix-loop-helix (HLH) structure (Figure 2). There are 19 residues that have at least 80% similarity (Figure 2). Based on the similarity of residues, the ‘basic’ region is the most conservative followed by the ‘helix’ region. The loop region shows a great variation compared to the other two regions. Based on the sequence characteristics of the ‘basic’ region, *CmTCP* proteins are divided into two classes: class I and class II. In addition, the *TCP* genes were again divided into three groups namely, PCF, CYC/TB1 and CIN based on the last three residues of the first ‘helix’ region. Proteins with TRE amino acids were put in the *PCF* group while proteins containing QDM and QDK/QDR were assigned to *CYC/TB1* and *CIN* groups, respectively. All Class I genes fell within the *PCF* group as well. Class I (*PCF*) contains 13 *CmTCP* proteins which have 4 amino acid residues less than that of class II proteins. The *CIN* group (class II) has 10 *CmTCP* proteins and the *CYC/TB1* group (class II) has 6 *CmTCP* proteins, respectively.

**Fig. 2.**
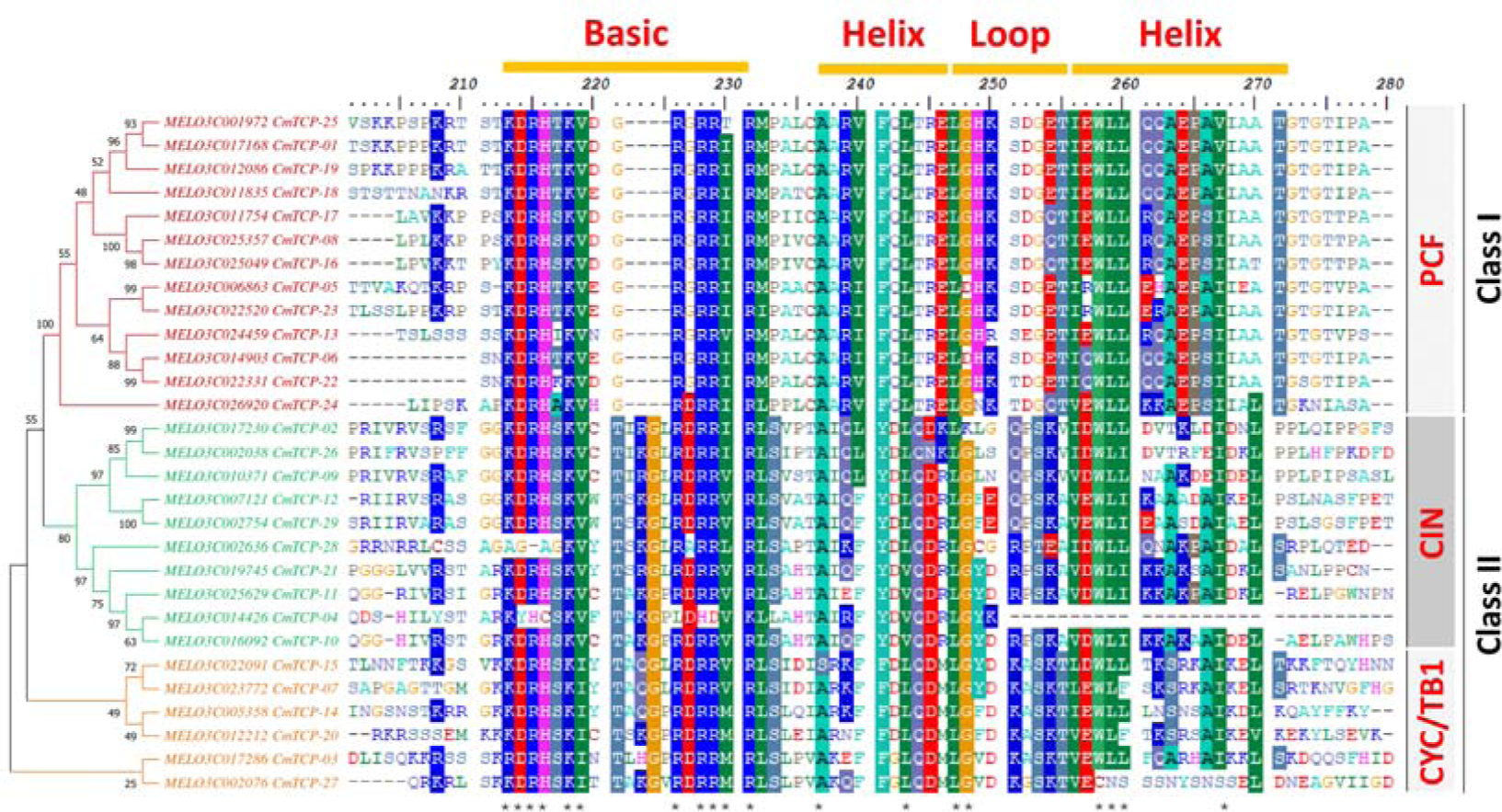
Alignment of 29 *CmTCP* protein sequences. Similar amino acids are shaded using 50% as the threshold value. Conserved residues (at least 80%) are indicated with an Asterisk (*) at the bottom. The alignment of domain-related regions of *CmTCP* protein sequences is shown here. The complete alignment is shown in Supplementary File 2.

### Phylogeny, Gene structure and Motif distribution

A phylogeny was constructed using 29 *CmTCP* proteins (Figure 3A). A total of 10 homologue gene pairs were found among the three groups. The *PCF* group contains four pairs (*CmTCP-25/CmTCP-01, CmTCP-08/CmTCP-16, CmTCP-05/CmTCP-23* and *CmTCP-06/CmTCP-22*) while *CIN* group consists three pairs (*CmTCP-02/CmTCP-26, CmTCP-12/CmTCP-29,* and *CmTCP-04/CmTCP-10*) and *CYC/TB1* group contains another three pairs of proteins (*CmTCP-15/CmTCP-07, CmTCP-14/CmTCP-20* and *CmTCP-03/CmTCP-27*). Closely related proteins generally contain similar structure and motif composition with some exceptions.

**Fig. 3.**
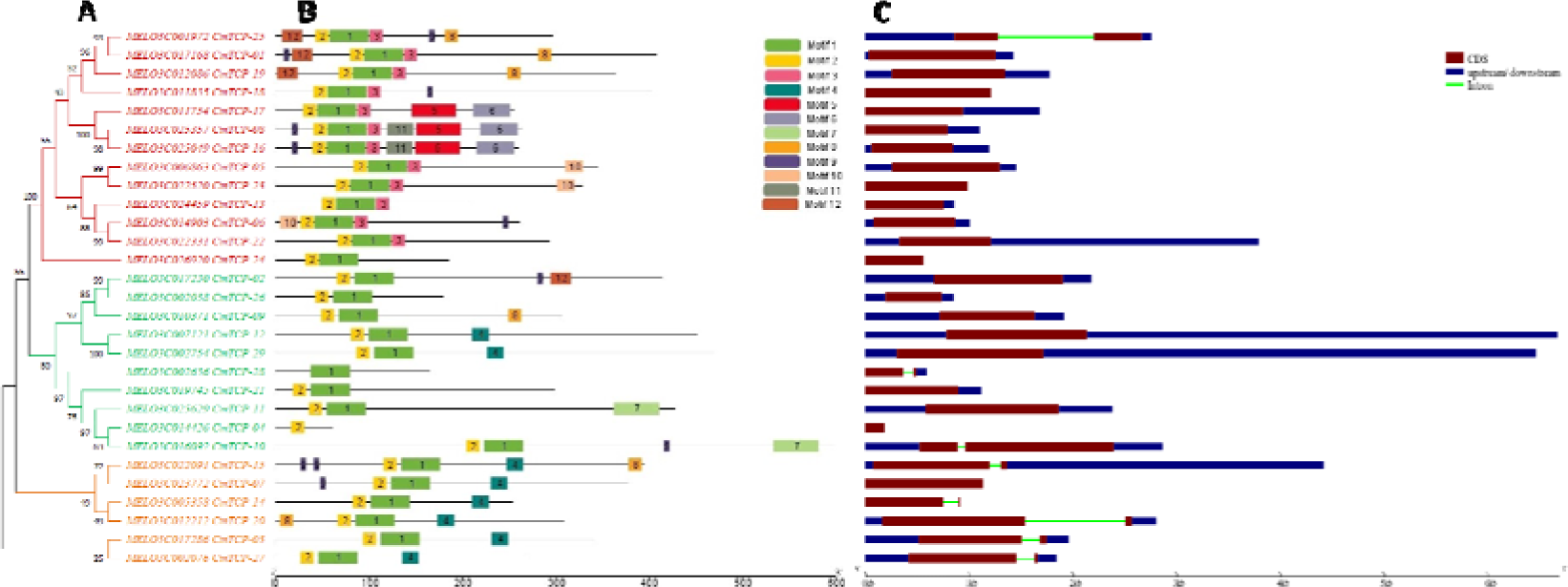
*CmTCP* sequence analysis- **(A)** Phylogenetic tree. *PCF*, *CIN* and *CYC/TB1* groups are indicated in red, green and gold colour, respectively; **(B)** Motif distribution. Motif numbers are indicated with coloured boxes as per the legends shown on the right; **(C)** Gene structure analysis. dark red, dark blue and bright green colours illustrate exons, UTRs and introns, respectively. Scales below the graphs are proportional to the length of the chromosomes, motifs and exons/introns.

A total of 12 motifs were identified in these 29 *TCP* genes (Figure 3B) of which Motif 1 and Motif 2 are widely distributed in every protein except *CmTCP-28* (lacks Motif 2) and *CmTCP-04* (lacks Motif 1). Alignment of *CmTCP* protein sequences (Figure 2) shows that the sequences of *CmTCP-28* and *CmTCP-04* are either mutated or incomplete. Hence, it can be assumed that Motif 1 and Motif 2 are probably located within the *TCP* domain region. There is a visual difference in motif distribution between the two classes. Motif 3 is a part of every Class-I (*PCF*) protein (except *CmTCP-24*). Motif 4 and Motif 7 are found in Class-II while Motif 5, Motif 6, Motif 10 and Motif 11 are found only in Class-I proteins. Motif 8, Motif 9 and Motif 12 are found in both classes. Interestingly, *CmTCP-15* contains two segments of Motif 9 side-by-side (Figure 3B).

The exon-intron structures of 29 *TCP* genes were analized to understand the characteristics of *TCP* gDNA which revealed that *TCP* genes have a maximum of two exons (Figure 3C). Twenty-one *CmTCP* genes do not have any introns while 8 genes (*CmTCP-03, CmTCP-10, CmTCP-14, CmTCP-15, CmTCP-20, CmTCP-25, CmTCP-27, and CmTCP-28*) contain a single intron (Figure 3C). The size of *CmTCP* genes ranges from 188 bp to 6690 bp (Table 1). *CmTCP-12* is found to be the largest gene followed by *CmTCP-29* while the smallest gene was *CmTCP-04.* Among six genes, 5 *CYC/TB1* group genes contain an intron except *CmTCP-07* (Figure 3C).

### Promoter Analysis

Gene expression patterns are largely determined by the cis-regulating elements present in the promoter region (Yamaguchi-Shinozaki and Shinozaki 2005). A total of 32 cis-acting elements were found involved in response to light responses (6), phytohormone responses (11), stress responses (11), and growth and development (4) (Wu et al. 2023).

We found Box 4, G-Box, GTI-motif, I-box, TCCC-motif and TCT-motif for light responsiveness. The elements for phytohormone responsiveness were divided into six groups, such as Methyl Jasmonate (CGTCA-motif and TGACG-motif), ABA (ABRE and DRE1), Auxin (AuxRR-core and TGA-element), Ethylene (ERE), Salicylic Acid (TCA-element) and Gibberellin (GARE-motif, P-box and TATC-box). Defence and stress response elements included as-1, MBS, MYC, WRE3, WUN-motif, ARE, DRE core, GC-motif, LTR, STRE and TC-rich repeats. Moreover, several other elements related to growth and development, such as meristem-specific gene activation, zein metabolism, circadian control element, and seed-specific expressions were found (Figure 4).

**Fig. 4.**
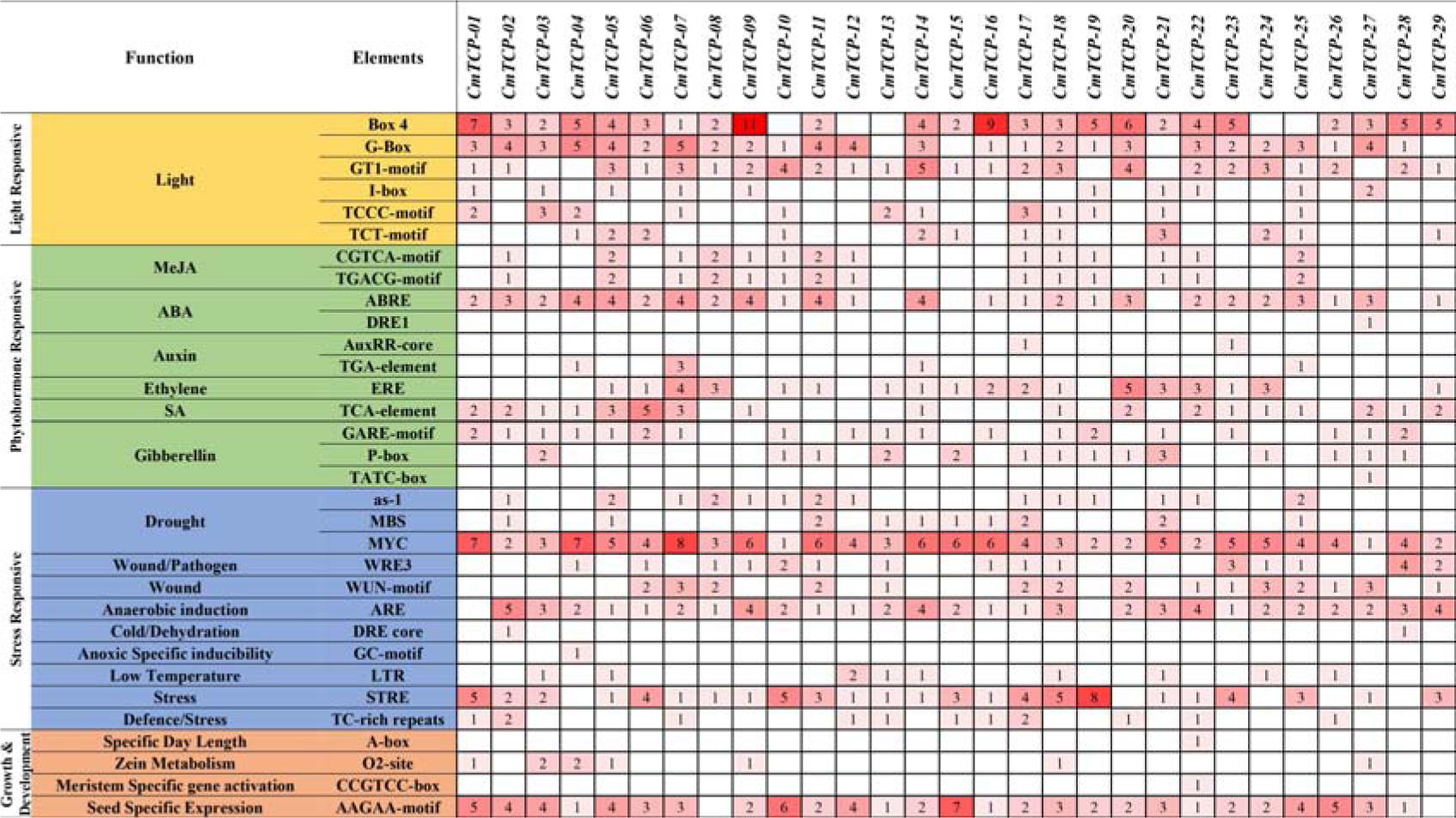
Cis-acting elements identified in the promoter region of *CmTCP* genes.

Each gene consists of multiple cis-regulating elements from each group. Drought-responsive element, MYC were found in the promoter region of every *CmTCP* gene whereas DRE1 and TATC-box were only found in *CmTCP-27* and A-box and CCGTCC-box in *CmTCP-22* (Figure 4). These results indicated that *CmTCP* genes might be involved in the response to various stresses, light, and phytohormones along with growth and development.

### Evolutionary Analysis of the *CmTCP* Gene Family

A total of 131 *TCP* proteins from 5 different plants namely, Melon (29 proteins), Cucumber (27 proteins), Rice (21 proteins), Arabidopsis (24 proteins), and Tomato (30 proteins) were used to study the evolution of *CmTCP* proteins in the melon genome (Figure 5) (Supplementary File 2).

**Fig. 5.**
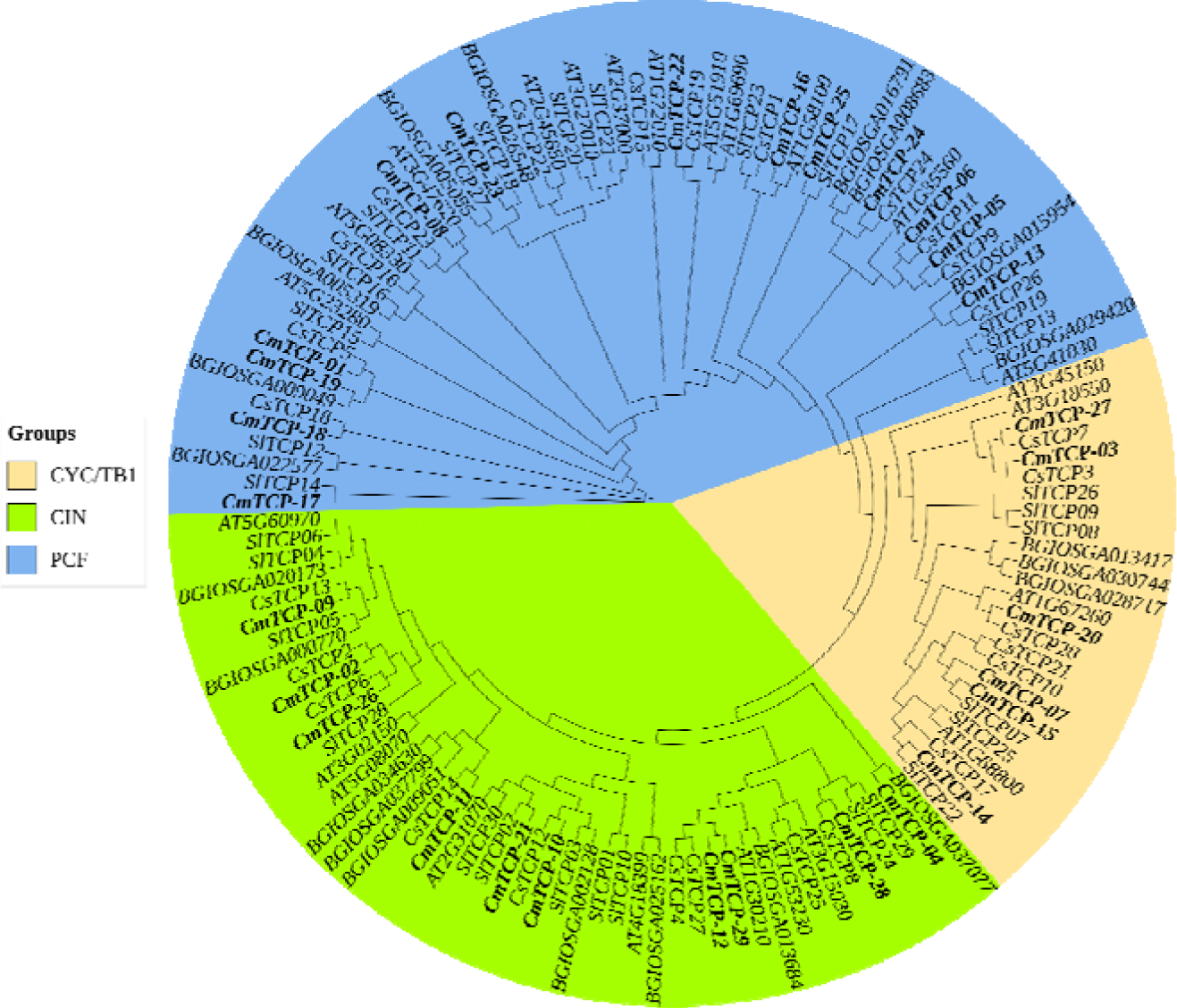
Phylogenetic analysis of *TCP* proteins in Melon (*CmTCP*), Cucumber (*CsTCP*), Rice (*BGIOSGA*), Arabidopsis (*At*), and Tomato (*SlTCP*). The phylogenetic tree was constructed by the Maximum likelihood method using 1000 bootstrap replicates. *CmTCP* proteins are highlighted in black bold letters.

Despite a huge difference in the size of the genome and the number of genes in these plants, the *TCP* encoding genes in each genome are more or less similar in number (Supplementary Table 2) in these genomes. The phylogenetic analysis also demonstrated that several of the *CmTCP* proteins were discovered to be more closely related to those of Cucumber followed by Tomato (Figure 5). For instance, *CmTCP09/CsTCP13, CmTCP26/CsTCP6, CmTCP28/CsTCP8, CmTCP21/SlTCP02, CmTCP14/SlTCP22,* and *CmTCP25/SlTCP17.* All the Arabidopsis *TCP*s in this phylogenetic tree belong to the same group as reported previously (Martin-Trillo and Cubas 2010).

### Gene Duplication and Syntenic Relationship

There are 15 whole genome duplications or segmental duplications (syntenic pairs) found among *CmTCP* genes with no evidence of tandem duplication (Figure 6, Supplementary Table 3). Out of 10 homologous gene pairs, seven showed gene duplication events except *CmTCP-08/CmTCP-16, CmTCP-04/CmTCP-10* and *CmTCP-15/CmTCP-07*. Moreover, eight syntenic pairs, *CmTCP-01/CmTCP-19, CmTCP-03/CmTCP-20, CmTCP-04/CmTCP-21, CmTCP-11/CmTCP-10, CmTCP-13/CmTCP-24, CmTCP-15/CmTCP-20, CmTCP-19/CmTCP-25* and *CmTCP-20/CmTCP-27* are non-homologous gene pairs. These results depict that whole genome duplication or segmental duplication probably plays an essential 13 role in the evolution of *CmTCP* genes together with other factors including random insertion events.

**Fig. 6.**
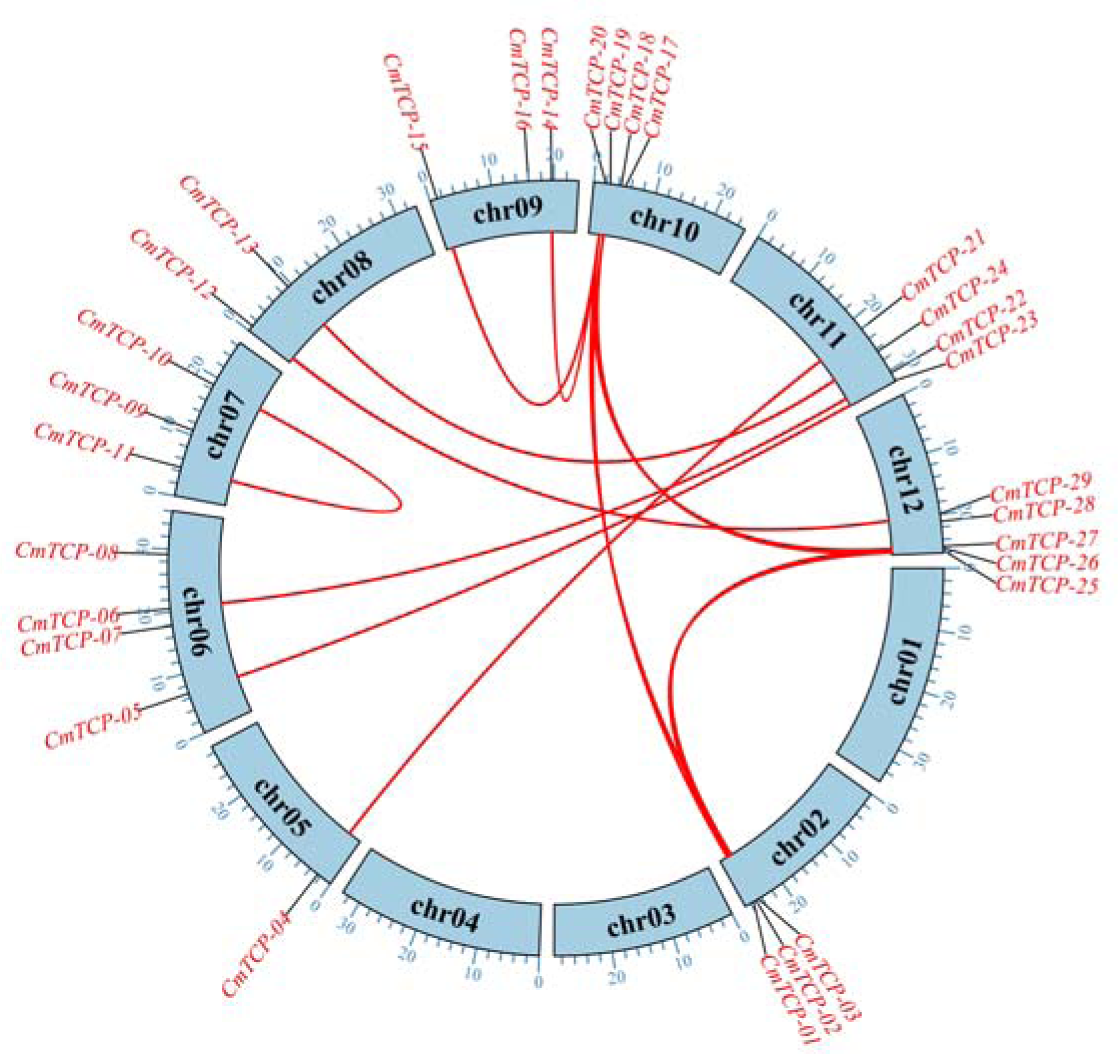
Gene duplication among 29 *CmTCP* genes. Duplicate genes are indicated using red lines.

To understand the evolution and abundance of *TCP* genes, syntenic relations of *CmTCP* genes with other crops were studied. A total of 151 syntenic pairs of *CmTCP* genes were calculated for these four species (Figure 7, Supplementary Table 3). Among them, the highest number of pairwise syntenic blocks were found in Cucumber (49) followed by Arabidopsis (48) and Tomato (43) whereas only 11 pairwise syntenic blocks were found between melon and rice (Figure 6, Supplementary Table 3). *CmTCP-20* showed the maximum number of syntenic blocks (14) followed by *CmTCP-19* (11) while no syntenic blocks were found in the case of *CmTCP-08, CmTCP-09* and *CmTCP-16* (Figure 7, Supplementary Table 3).

**Fig 7.**
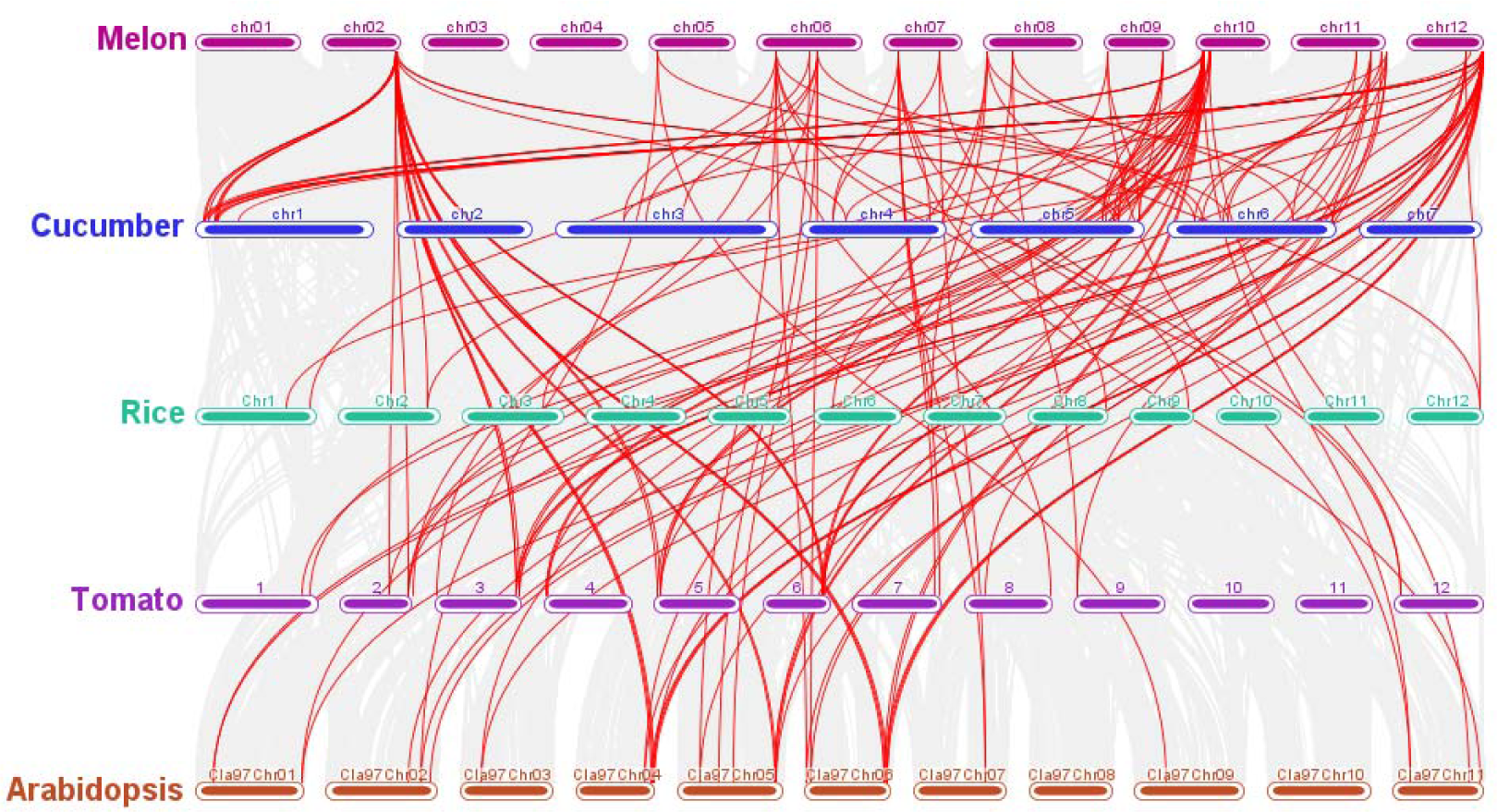
Syntenic relationships of 29 Melon *CmTCP* genes with that of Cucumber, Rice, Tomato and Arabidopsis. The grey lines in the background represent the collinear blocks within Melon and other plant genomes, while the red lines highlight the syntenic *TCP* gene pairs.

### Expression pattern analysis of *CmTCP* genes

RNA expression data of 29 *CmTCP* genes from 45 samples of 25 vegetative and reproductive tissues were extracted from Melonet DB (in FPKM unit) (Figure 8). Among them, the highest number of genes (twenty-four) were expressed in shoot apex followed by tendril and young leaves where 21 and 20 genes were expressed, respectively. The lowest number of genes (nine) were expressed in the fruit flesh after a week of harvesting (Supplementary Table 4). This result indicates that *CmTCP* genes are more expressed in the vegetative stage compared to the reproductive stage. No expression records were found for five genes (*CmTCP-03, CmTCP-07, CmTCP-10, CmTCP-11,* and *CmTCP-23*) whereas eight genes (*CmTCP-05, CmTCP-06, CmTCP-12, CmTCP-13, CmTCP-17, CmTCP-18, CmTCP-22,* and *CmTCP-29*) were expressed in every samples (Supplementary Table 4). The expression patterns indicate that *CmTCP-17* and *CmTCP-22* were highly expressed in every tissue compared to others, and may have an influence on overall growth and development. According to the clusters, 11 genes were moderately expressed and 16 genes were either expressed at a very low level or did not show any expression. Several genes are expressed in a specific tissue much higher than in other tissues indicating their regulation of the tissue- specific phenotypes. For example, *CmTCP-09 was* expressed higher in anther and petals whereas *CmTCP-13 was* expressed in dry seed and fruit epicarp. Such expression data illustrates the possible function and involvement of *CmTCP* genes in various stages of plant growth and development.

**Fig. 8.**
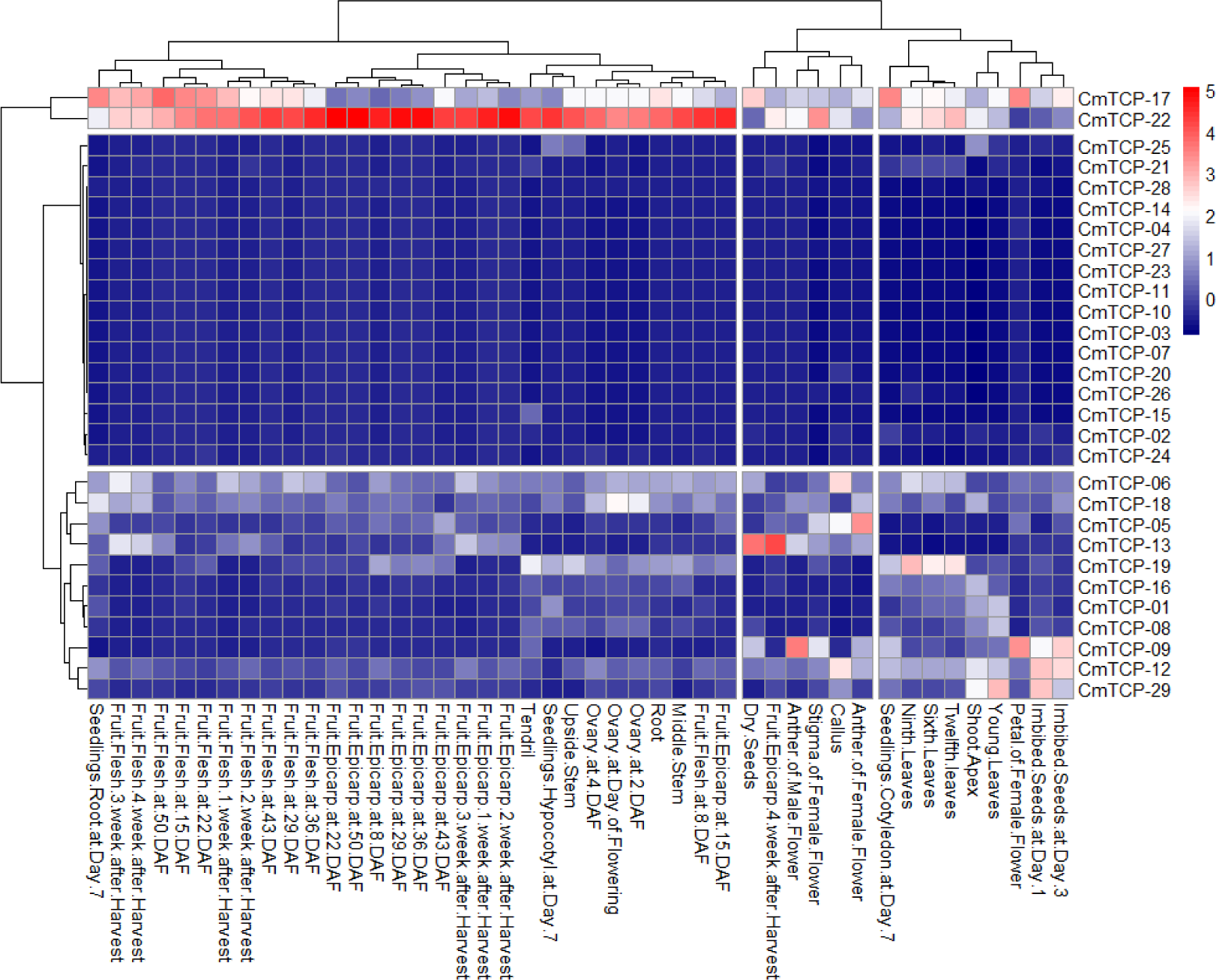
Expression pattern of *CmTCP* genes in different melon tissues and developmental stages. The heatmap was generated by the RNA expression data (FPKM) from Melonet-DB.

**Fig. 9.**
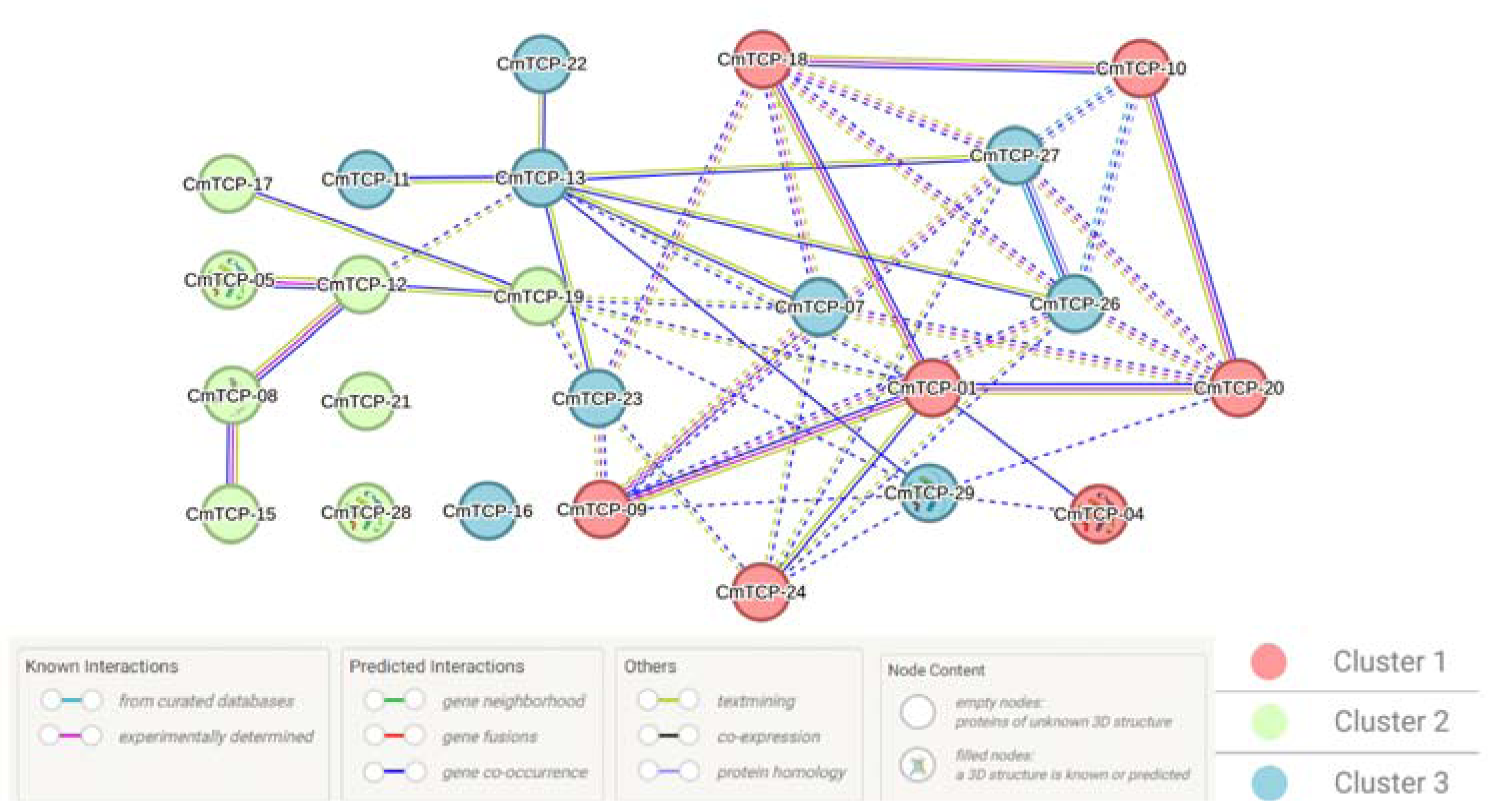
Predicted network of protein-protein interaction of *CmTCP* proteins. Each node indicates a *CmTCP* protein and lines indicate association among them. Different colours indicate clusters predicted using the K-means method. Each node represents a protein and the lines are for interactions among them.

### Protein Structure Prediction and Sequence Similarity

Protein homology modeling is a powerful technique used to predict the three-dimensional structure of a protein based on its amino acid sequence similarity to known protein structures. In this study, the Swiss Model, a widely used homology modeling tool was used to generate structural models for the target proteins. Predicted structures of *CmTCP* proteins along with template information are presented in Supplementary File 3. GMQE (Global Model Quality Estimation) is a measure used in protein homology modeling to estimate the reliability of a generated model. The highest GMQE value was found in the case of *CmTCP-26* (0.65) and the lowest was for *CmTCP-10* (0.36). Proteins of each group showed a similar pattern in the number of alpha helix and beta sheets (Supplementary File 3). The sequence similarity among the *CmTCP* proteins ranged from 4.69 (*CmTCP-0* vs *CmTCP-18)* to 93.58 (*CmTCP- 08* vs *CmTCP-16)*. For the PCF group of proteins, the range is 21.14 (*CmTCP-24* vs *CmTCP- 18)* to 93.58 (*CmTCP-08* vs *CmTCP-16)* which is 7.05 (*CmTCP-04* vs *CmTCP-29)* to 65.05 (*CmTCP-12* vs *CmTCP-29)* and 27.21 (*CmTCP-15* vs *CmTCP-27)* to 45 (*CmTCP-07* vs *CmTCP-15)* for CIN and CYC/TB1 groups respectively. Comparatively, proteins from similar groups showed a higher similarity percentage (Supplementary Table 5).

### Predicted network of *CmTCP* proteins

The protein-protein interaction network includes three clusters of 24 genes predicted with String-db. Cluster 1 includes seven proteins, whereas Cluster 2 includes eight proteins and the rest belong to Cluster 2 (Figure 8). Several proteins like *CmTCP-01, CmTCP-07, CmTCP-09, CmTCP-10, CmTCP-13, CmTCP-18, CmTCP-26, CmTCP-27,* and *CmTCP-29* are highly connected to other proteins and bear main nodes. The result suggested that these proteins are possibly responsible for diverse functions in the melon genome.

## DISCUSSION

The *TCP* gene family has been found to be involved in a range of biological processes (Martin-Trillo and Cubas 2010) including light response (Chen et al. 2024), reproductive development (Viola and Gonzalez 2023), growth and development in plants (Jiang et al. 2023) and environmental adaptation (Wangdan Xiong and Juan 2022). We employed *in silico* bioinformatical analyses to identify and analyze the genome-wide *TCP* family genes, in a way to understand the genomic features, evolutionary processes and functional characteristics of those in melon.

### *TCP* Family Genes: Numbers & Chromosomal Distribution

The numbers of *TCP* genes in different plant species generally ranges from 17 to 75 genes (Li et al. 2022) and varies widely among species such as Arabidopsis (24), Cucumber (27), Potato (23), Rice (22), and Cotton (74) etc. (Bao et al. 2019; Li 2015; Wen et al. 2020; Yao et al. 2007; Zheng et al. 2018). We have identified 29 *TCP* genes in the melon genome (Table 1). The probable reason for the variation in the number of *TCP* genes might be the differences in genome size, chromosome numbers, gene duplication and evolutionary pathway etc. (Huang et al. 2023). Most of the genes seemed to be distributed throughout the chromosomes and mostly toward the telomeric region (Figure 1). Similar patterns of chromosomal distributions of *TCP* family genes were observed in the case of *Melastoma candidum* (Li et al. 2022), Ginger (Jiang et al. 2023), Tea (Shang et al. 2022), and *Dendrobium chrysotoxum* (Huang et al. 2023). All of the *CmTCP* proteins are predicted to be localized in the nucleus except *CmTCP-14* which is localized in cytoplasm (Table 1). As *CmTCP* proteins are located at the nucleus, they can regulate gene expression as transcription factors binding with DNA target sequences (Xiao et al. 2022).

### Gene Structure, Motif and Protein Domains

The structure of most of the *CmTCP* genes is very simple with either only one or no introns at all (Figure 3C). Of the 29 *CmTCP* genes, only 8 genes have introns. This may be because introns were lost during the evolution of gene structures (Ma et al. 2014). Intron-free genes often respond fast to stress, even though they are not suitable for recombination or species evolution (Sang et al. 2016). Furthermore, as shown in Figure 3, *CmTCP* genes within the same group displayed similarity in exon-intron structures and conserved motifs, indicating a strong evolutionary relationship among these genes. Every protein group (Figure 3A), with a few notable exceptions, has a distinct variation in the distribution of their motifs, such as the unique distribution of motifs 3 and 4 (Figure 3B). The complex structure of *TCP* proteins revealed that they contain a distinct DNA-binding motif (Zhang et al. 2023b).

Despite some variations, proteins of each group also exhibited a similar pattern in their structures, especially in the number of alpha helix and beta sheets (Supplementary File 3). The functional differentiation of the *CmTCP* protein is supported by the unique motif distribution and structural homology among different groups. A basic-helix-loop-helix (bHLH) structure with 59 amino acids residue was found to form the *TCP* domain in melon (Figure 2) which is similar to the findings of previous studies (Aggarwal et al. 2010; Shang et al. 2022). Sequence alignment of 29 *CmTCP* genes exhibits a highly conserved *TCP* domain which can be divided into two classes (Li 2015). The classification is based on 4 residual differences within the basic region (Figure 2). Li (Li 2015) explained the event as a residual insertion in Class-II genes. Through the *bHLH* structure and sequence similarity with other species (Jiang et al. 2023; Li 2015; Ma et al. 2014; Shang et al. 2022), it can be assumed that the *TCP* domain region is relatively conserved in the melon evolutionary process.

### Evolutionary Relationship

The sequence differences of *TCP* genes throughout the species could be the result of diverse plant selection processes such as natural selection, mutation or evolution. Using whole- genome data from several species, synteny analysis can be used to hypothesize the locations of both orthologous and paralogous genes (Cao et al. 2016; Lin et al. 2014). Moreover, Gene duplication can produce new genetic material that can be used by selection, drift, and mutation to produce specialized or novel gene functions (Magadum et al. 2013). The variation within the *TCP* genes indicated that *CmTCP* genes might be involved during the process of evolution from its common and close ancestors (Figure 6) and gene duplication results in abundance and functional differences of *CmTCP* genes. Phylogenetic analyses of homologous genes across multiple genomes can provide comprehensive insights into the extent of evolutionary and functional diversification within a specific gene family (Stavrinides and Ochman 2009).The phylogenetic analysis showed that *CmTCP* genes are closely related with *CsTCP* genes indicating the close evolutionary relationship between melon and cucumber. Some of the *CmTCP*s were found to be closely clustered with homologs of cucumber, this suggested that these proteins might serve similar roles in melon and cucumber (Pearson 2013).

### Cis-regulating Elements

Several important cis-regulating elements were found to be present in the promoter analysis. These elements included those related to the expression of meristems and endosperms, responses to light, hormones (such as gibberellin, auxin, salicylic acid, MeJA, and auxin), stress (anaerobic, low temperature), and drought response. These results suggest that the *CmTCP* gene family may have roles in regulating hormone levels to govern melon growth and development in addition to participating in responses to abiotic stresses.

### Putative Functions of *TCP* gene family

Class II CYC/TB1 clade *TCP* genes are primarily involved in controlling the development of lateral branches or flowers by regulating axillary meristems (Zhang et al. 2023a). The proteins are known to control the transition from cell division to leaf expansion (Kazama et al. 2010; Qi et al. 2017). The size, shape, flatness, and complexity of the leaf lamina are significantly determined by the time of the transition. The *CYC/TB1* group of proteins essentially assist in the regulation of axillary meristem development, which leads to the formation of either lateral branches or flowers (Tähtiharju et al. 2011; Yu et al. 2021). Insights into gene function are frequently obtained by analyzing the temporal and spatial expression patterns of a gene within the species . In order to find out if *CmTCP* plays any crucial role in growth and development, especially in lateral branching, we consequently looked at the expression patterns to understand gene expression and phylogeny of the proteins to predict functional homology by comparing with the genes of other species with known roles.

### Expressions of *TCP* family genes in melon

The expression pattern reveals that almost every *CmTCP* gene, except *CmTCP-23* from the *PCF* group (Class-I) is moderately to highly expressed in every tissue (Figure 7, Supplementary Table 4). It indicates the possible involvement of class-I genes in the overall growth and development of various organs including leaves, flowers, fruits and others (Viola et al. 2023). In previous studies, it was found that class-I genes are involved in the elongation of leaves and internode (Kieffer et al. 2011; Koyama et al. 2010), pollen and flower bud development (Balsemao-Pires et al. 2013; Takeda et al. 2006), trichome branching (Camoirano et al. 2020), and fruit ripening (Parapunova et al. 2014). *SlTCP18* a homolog of *CmTCP-18* was predicted to control the fruit ripening in tomatoes (Parapunova et al. 2014). Though the genes were closely related to each other, no expression-data in fruit tissue was found for *CmTCP-18* gene (Figure 7, Supplementary Table 4).

The *CIN* group of *TCP* genes are found to be expressed highly in flower buds and control plant flowering (Lan and Qin 2020; Wen et al. 2020). *Petunia* PhLA, a *CIN*-*TCP* gene, has been shown to be involved in flower morphogenesis (Chen et al. 2020b). Most of the *CsTCP* genes from the *CIN* group in cucumber were found to be involved in the regulation of female flower bud development. Homologs of these genes like *CmTCP-09, CmTCP-12,* and *CmTCP-29* showed higher expression in floral tissue indicating their potential involvement in floral development. As *CmTCP-12,* and *CmTCP-29* are expressed in every type of tissues, these genes might have roles in the overall growth and development (Figure 7, Supplementary Table 4). In addition, *CmTCP-09* might be responsible for floral growth and development as it was found to be expressed in mostly floral parts.

Previous studies suggest that *CYC/TB1* genes are responsible for the regulation of branching in different plants (Bai et al. 2012; Poza-Carrion et al. 2007; Shang et al. 2022; Wen et al. 2020; Zhao et al. 2018). *CsTCP21,* a homolog of *CmTCP-15* was found to be involved in the regulation of tendril growth and development. Similar expression data indicates the involvement of *CmTCP-15* in tendril growth and development as the gene was found to be expressed only in tendril (Figure 7, Supplementary Table 4)*. CsTCP3* and *CsTCP7, an* ortholog of *AtTCP18* (AT3G18550), were found to be expressed in auxiliary buds of cucumber and *CsTCP3* was predicted to be responsible for lateral branching in cucumber (Aguilar-Martinez et al. 2007; van Es et al. 2019; Wen et al. 2020). Interestingly, *CmTCP-03* and *CmTCP-27* were closely related with *CsTCP3, CsTCP7,* and *AtTCP18* (Figure 4). The expression data indicated that *CmTCP-03* and *CmTCP-27* were not expressed in any of the tissues (Figure 7, Supplementary Table 4), probably due to the lack of expression data from the auxiliary bud. Though the Melonet DB does not contain any expression-related data from the auxiliary buds, based on the homologs, we can predict that these genes might be organ- specific (expressed in auxiliary bud) and probably regulate lateral branching in melon as well. The primary endogenous regulators of plant growth and development are GA and ethylene (Katyayini et al. 2020; Wang et al. 2010). The presence of TATC-motif also justifies the involvement of *CmTCP-27* in lateral branching as TATC-motif is a GA-related cis-element and was found to affect the growth and development of lateral organs (Wen et al. 2020).

## CONCLUSION

The present study identifies 29 *TCP* genes in the melon genome, distributed mainly in the telomeric regions of 9 chromosomes. Structure, motif, domain distribution and sequence alignment indicated that the genes are highly conserved in this species. Gene duplication and synteny analysis depict that the genes are developed during evolution as well as from duplication within the genome. Phylogeny and expression data indicated that *CmTCP-09* might be responsible for floral development, and *CmTCP-03* and *CmTCP-27* probably regulate lateral branching in melon. The findings paves the way for any future functional genomic study aiming at elucidating the exact functions of the *TCP* genes in melon which then can be used for improvement of melon via breeding and biotechnological means.

## Acknowledgements

Not applicable

## Author Contributions

MJHJ, MNAS and MRH were responsible for conceptualization. MJHJ and MNAS was responsible for data collection and curation. MJHJ was responsible for analysis and writing the original draft. MKB was responsible for synteny analysis. MRH were responsible for fund acquisition, supervising the work, reviewing and finalizing the draft. All authors have read and agreed to the final version of the manuscript.

## Ethical approval

Not applicable.

## Informed consent

Not applicable.

