## Supplementary File 1 for "Genome-Wide Analysis of *TCP* Family Genes and Their Constitutive Expression Pattern Analysis in the Melon (*Cucumis melo*)"

### *Supplementary File 1: Sequence Alignment of 29 TCP proteins in Melon*

```
                   10         20         30         40         50         60         70         80         90        100
           ....|....| ....|....| ....|....| ....|....| ....|....| ....|....| ....|....| ....|....| ....|....| ....|....|
MELO3C001972 ---------- ---------- ---------- ---------- ---------- ---------- ---------M EGNDQLHYHQ NHRSN----- ----------
MELO3C017168 ---------- ---------- ---------- ---------- ---------- ---------- ---------M EGGDELHHHH HHHHQNHHHQ Q---------
MELO3C012086 ---------- ---------- ---------- ---------- ---------- ---------- -------MIM DGENGIRR-- ---------- ----------
MELO3C011835 ---------- ---------- ---------- ---------- ---------- ---------- ---------- ---------- ---------- ----------
MELO3C011754 ---------- ---------- ---------- ---------- ---------- ---------- ---------M SNSDHTNG-- ---------- ----------
MELO3C025357 ---------- ---------- ---------- ---------- ---------- ---------- ---------M SASDAVN--- ---------- ----------
MELO3C025049 ---------- ---------- ---------- ---------- ---------- ---------- ---------M SASDGVN--- ---------- ----------
MELO3C006863 ---------- ---------- ---------- ---------- ---------- ---------- ---------M EEVNQNHRLY EKEGSSNG-- ----------
MELO3C022520 ---------- ---------- ---------- ---------- ---------- ---------- ---------M AD-NQKQELD DDDDSPP--- ----------
MELO3C024459 ---------- ---------- ---------- ---------- ---------- ---------- ---------M GSDIVLHKFP S--------- ----------
MELO3C014903 ---------- ---------- ---------- ---------- ---------- ---------- ---------M GENK------ ---------- ----------
MELO3C022331 ---------- ---------- ---------- ---------- ---------- ---------- ---------M ADQNNNNNNN NKHHPLLPSP ATSSSPLTTP
MELO3C026920 ---------- ---------- ---------- ---------- ---------- ---------- ---------- ---------- ---------- ----------
MELO3C017230 ---------- ---------- ---------- ---------- ---------- --------MI YENYKDMLRN SNSKDQKGSE AKEEGIS--- ----------
MELO3C002038 ---------- ---------- ---------- ---------- ---------- --------MI SK-----LKE DGAHD----E AKQ------- ----------
MELO3C010371 ---------- ---------- ---------- ---------- ---------- --------MP AD------KH DERHNMIKNS KEE------- ----------
MELO3C007121 ---------- ---------- ---------- ---------- ---------- ------MEVV DNIQEQPRKF PRLANGRSSD SNEIGLK--- ----------
MELO3C002754 ---------- ---------- ---------- ---------- ---------- ------MEVV D-IQEQACRF PRIGNGRN-D SNKLGQK--- ----------
MELO3C002636 ---------- ---------- ---------- ---------- ---------- ---------- ---------- ---------- ---------- ----------
MELO3C019745 ---------- ---------- ---------- ---------- ---------- ---------- ---------- ---------- ---------- ----------
MELO3C025629 ---------- ---------- ---------- ---------- ---------- ---------- ---------M GEFSNQRPAK SAKPG----- ----------
MELO3C014426 ---------- ---------- ---------- ---------- ---------- ---------- ---------- ---------- ---------- ----------
MELO3C016092 MAELFGRGRR GGSSFISKSR SNRLQSGGSS SNCRKKKERE RREKKIEEAI RIRKKHCPFK AGRRKAKQSE SGSSTKSPSS SSKPDSNFEA CRRKSRGRRP
MELO3C022091 ---------- ---------- ---------- ---------- ---------- ---MFSSTNN LHLFPLQQYF PSSSSSSPSY HHHHLVAPLP SPPPHHHHPE
MELO3C023772 ---------- ---------- ---------- ---------- ---------- ------MGSS SYNPNPFPCF PSSSSSTTNS YNLPLSIPSL LN--------
MELO3C005358 ---------- ---------- ---------- ---------- ---------- ----MDDHQQ QQHRENQILI NSYDDHDLHY FPHPFLDD-- ----------
MELO3C012212 ---------- ---------- ---------- ---------- ---------- ------MIIK SSFWATFCIL KNTNYLQTIL IKLELLK--- ----------
MELO3C017286 ---------- ---------- ---------- ---------- ---------- ---MFVFNSC SNNNHGNDYS ISFPDHSFLH FPSPFDDD-- ----------
MELO3C002076 ---------- ---------- ---------- ---------- ---------- ---------- ---------- ---------- ---------- ----------

                  110        120        130        140        150        160        170        180        190        200
           ....|....| ....|....| ....|....| ....|....| ....|....| ....|....| ....|....| ....|....| ....|....| ....|....|
MELO3C001972 ---------- ---------- ---------- ---------- -----ASFPL QLLEKK-EYD -------PQP PLEI------ ---------- ----------
MELO3C017168 ---------- ---------- ---------- ---------- HHQHRPTFPF QLLEKK-DED AASCSTSPYP SLAISA---- ---DTATTTP TTNSSVVAAE
MELO3C012086 ---------- ---------- ---------- ---------- -----PNFPL QLLEKKKEEE EEQQQQHPCS SSANNGGTK- ---TTTTITT TPNSNLQSSS
MELO3C011835 ---------- ---------- ---------- ---------- ----MASTSS SASTHQLDAS LVIATRSDTT AAAADP---- ---------- -------SPR
MELO3C011754 ---------- ---------- ---------- ---------- -----VSNAA AID-SHKN-- -------QSS SNGA------ ---------- ----------
MELO3C025357 ---------- ---------- ---------- ---------- -----NNNAA IIEHSHQQHT HRIHGNNQSS NNAATP---- ---------- ----------
MELO3C025049 ---------- ---------- ---------- ---------- -----NNNAA IIDPSHQQHI HQIQAN-QSS NNATTP---- ---------- ----------
MELO3C006863 ---------- ---------- ---------- ---------- GYENLTNSTV DSSNPAKAED VLSDRHVGPP STAVPKKEKE GIMEHGLDIM PPLSLSQVPS
MELO3C022520 ---------- ---------- ---------- ---------- -----THLRI NDSDSSRMDL LPLTPKQEHP DSNLP----- ----HDLLTF NP----DIPK
MELO3C024459 ---------- ---------- ---------- ---------- -SSDPHSQPP TASHSVVGSN SSPATISRQS LLPRKSN--- ---------- ----------
MELO3C014903 ---------- ---------- ---------- ---------- -PAEIKDFQI AIAEKEEGK- --------KQ LAPKRS---- ---------- ----------
MELO3C022331 PLPT------ ---------- ---------- ---------- PSSDIKDFTL IISNKDQNNS NSNSNNNNKQ LAPKRT---- ---------- ----------
MELO3C026920 ---------- ---------- ---------- ---------- -----MGSKN MRVKKPTSNS PRNAPKLNSP IASESK---- ---------- ----------
MELO3C017230 ---------- ---------- ---------- ---------- ---------- -IRSSSSNNN DGKYSFPKAP TSSSRQW--- ---------- ----SGFR-N
MELO3C002038 ---------- ---------- ---------- ---------- ---------- ---------M EGKR---KKA RTCSRQG--- ---------- ----QFLRGN
MELO3C010371 ---------- ---------- ---------- ---------- ---------- --------NS PPPKLAAGSS SSSATPW--- ---------- ----LRLK-D
MELO3C007121 ---------- ---------- ---------- ---------- ------GAEN VRRGDEKDKE -SKRDGGITG VGG------- -GGVGGADDE TNRLRGWHHS
MELO3C002754 ---------- ---------- ---------- ---------- ------GIVQ YSPDDEEDKE FSKRVAVVTS ATVNPTGRCE PGGIGGVG-- ANRLHGWHHS
MELO3C002636 ---------- ---------- ---------- ---------- ---------- ---------- ---------M GSLDDHD--- ---------- ----KKKAKI
MELO3C019745 ---------- ---------- ---------- ---------- ---------- ---------- -------MGI NTIAG----- ---------- -----QILHV
MELO3C025629 ---------- ---------- ---------- ---------- ---------- ---------- ---------V RTCGD----- ---------- -GGGSEIVEV
MELO3C014426 ---------- ---------- ---------- ---------- ---------- ---------- ---------M KTNTY----- ---------- ----DEIVLV
MELO3C016092 RFSTPPPPRP PPPPSSAPTA SFQFLDSQSQ QPHQQQAPKK RSYFQLPPPH PVAAPAASAS TFADQSIEYA RKMGESHRQT STSSRLGMRT GGGGGEIVEV
MELO3C022091 PVYIGPPQDP VLSG------ ---------- ---------- ---VVAPIFL QVQEEDRDQA RMKNEQQEVA VLMSQNG--- -----NNNNN SNGQCFSSST
MELO3C023772 ---------- ---------- ---------- -------FDP SSNNTSTVNF HHQPHQDPLS FLAPYFPIAH FSTLTPPETA VINFAVAGNN IQALGDMVVP
MELO3C005358 ---------- ---------- ---------- ---------- ---DDLFLSH FLSQQQQFFE QRDQQLDETT KKMKKKK--- ---------- -KLCGSSSSR
MELO3C012212 ---------- ---------- ---------- ---------- -----VAITQ TMKQQQQQQP IIPPNTKPKP IIVHQNP--- ---------- ----------
MELO3C017286 ---------- ---------- ---------- ---------- ---GNPTNSL LLQQQSQHDI FLHHHIPLNE PSPPPPSTFV N----ALRSS ETINNDVFHQ
MELO3C002076 ---------- ---------- ---------- ---------- ---------- -SPPIDNMSM MMMNSDHKMK KMMKKVP--- ---------- ----------

                  210        220        230        240        250        260        270        280        290        300
           ....|....| ....|....| ....|....| ....|....| ....|....| ....|....| ....|....| ....|....| ....|....| ....|....|
MELO3C001972 VSKKPSPKRT STKDRHTKVD G----RGRRT RMPALCAARV FQLTRELGHK SDGETIEWLL QQAEPAVIAA TGTGTIPA-- ---------- ----------
MELO3C017168 TSKKPPPKRT STKDRHTKVD G----RGRRI RMPALCAARV FQLTRELGHK SDGETIEWLL QQAEPAVIAA TGTGTIPA-- ---------- ----------
MELO3C012086 SPKKPPPKRA TTKDRHTKVD G----RGRRI RMPALCAARV FQLTRELGHK SDGETIEWLL QQAEPAVIAA TGTGTIPA-- ---------- ----------
MELO3C011835 STSTTNANKR STKDRHTKVE G----RGRRI RMPATCAARV FQLTRELGHK SDGETIEWLL QQAEPAIIAA TGTGTIPA-- ---------- ----------
MELO3C011754 ----LAVKKP PSKDRHSKVD G----RGRRI RMPIICAARV FQLTRELGHK SDGQTIEWLL RQAEPSIIAA TGTGTTPA-- ---------- ----------
MELO3C025357 ----LPLKKP PSKDRHSKVD G----RGRRI RMPIVCAARV FQLTRELGHK SDGQTIEWLL RQAEPSIIAA TGTGTTPA-- ---------- ----------
MELO3C025049 ----LPVKKT PYKDRHSKVD G----RGRRI RMPIVCAARV FQLTRELGHK SDGQTIEWLL RQAEPSIIAT TGTGTTPA-- ---------- ----------
MELO3C006863 TTVAKQTKRP S-KDRHTKVE G----RGRRI RMPAACAARI FQLTRELDHK SDGETIRWLL EHAEPAIIEA TGTGTVPA-- ---------- ----------
MELO3C022520 TLSSLPPKRP STKDRHTKVE G----RGRRI RIPATCAARI FQLTRELGHK SDGETIRWLL ERAEPAIIAA TGTGTIPA-- ---------- ----------
MELO3C024459 ----TSLSSS SSKDRHIKVN G----RGRRV RMPALCAARI FQLTRELGHR SEGETIEWLL RQAEPSIIAA TGTGTVPS-- ---------- ----------
MELO3C014903 ---------- SNKDRHTKVE G----RGRRI RMPALCAARI FQLTRELDHK SDGETIQWLL QQAEPSIIAA TGTGTIPA-- ---------- ----------
MELO3C022331 ---------- SNKDRHKKVD G----RGRRI RMPALCAARV FQLTRELGHK TDGETIQWLL QQAEPSIIAA TGSGTIPA-- ---------- ----------
MELO3C026920 -----LIPSK APKDRHAKVH G----RDRRI RLPPLCAARV FQLTRELGNK TDGQTVEWLL KKAEPSIIAL TGKNIASA-- ---------- ----------
MELO3C017230 PRIVRVSRSF GGKDRHSKVC TIRGLRDRRI RLSVPTAIQL YDLQDKLKLG QPSKVIDWLL DVTKLDIDNL PPLQIPPGFS PFHQQMLFPQ YVD-------
MELO3C002038 PRIFRVSPFF GGKDRHSKVC TIKGLRDRRI RLSIPTAIQL YDLQNKLGLS QPSKVIDWLI DVTRFEIDKL PPLHFPKDFD P--------- ----------
MELO3C010371 PRIVRVSRAF GGKDRHSKVC TIRGLRDRRV RLSVSTAIQL YDLQDRLGLN QPSKVVDWLL NAAKDEIDEL PPLPIPSASL G--------- ----------
MELO3C007121 -RIIRVSRAS GGKDRHSKVW TSKGLRDRRV RLSVATAIQF YDLQDRLGFE QPSKAVEWLI KAAADAIKEL PSLNASFPET PKQLSGEKFS VTDQPGTLDS
MELO3C002754 SRIIRVARAS GGKDRHSKVW TSKGLRDRRV RLSVATAIQF YDLQDRLGFE QPSKAVEWLI EAASDAIAEL PSLSGSFPET PRQLSDEKMG SDGADQGFDS
MELO3C002636 GRRNRRLCSS AGAG-AGKVY TSKGLRARRL RLSAPTAIKF YDLQDRLGCG RPTEAIDWLL QNAKPAIDAL SRPLQTED-- ---------- ----------
MELO3C019745 PGGGLVVRST ARKDRHSKVY TSRGLRDRRV RLSAHTAIQF YDVQDRLGYD RPSKAVDWLI KKAKSAIDKL SANLPPCN-- ---------- ----------
MELO3C025629 QGG-RIVRSI GRKDRHSKVC TAKGPRDRRV RLSAHTAIEF YDVQDRLGYD RPSKAVDWLI KKAKPAIDKL -RELPGWNPN VLDMSTQKLT KQAEKKSENK
MELO3C014426 QDS-HILYST ARKYHCSKVF TAKGPLDHDV KLLAHTAIRF YDVQDRLGYK ---------- ---------- ---------- ---------- ----------
MELO3C016092 QGG-HIVRST GRKDRHSKVC TAKGPRDRRV RLSAHTAIQF YDVQDRLGYD RPSKAVDWLI KKAKAAIDEL -AELPAWHPS TGNASTQGEE QQS-LNDENE
MELO3C022091 TLNNFTKKGS VKKDRHSKIY TAQGLRDRRV RLSIDISRKF FDLQDMLGYD KASKTLDWLL TKSRKAIKEL TKKFTQYHNN N--------- ----------
MELO3C023772 SAPGAGTTGM GKKDRHSKIY TAQGLRDRRV RLSIDIARKF FDLQDMLGYD KASKTLEWLF SKSRKAIKEL SRTKNVGFHG G--------- ----------
MELO3C005358 INGSNSTKRR GKKDRHSKIY TAQGPRDRRM RLSLQIARKF FDLQDMLGFD KASKTIEWLL LNSNSAIKDL KQAYFFKY-- ---------- ----------
MELO3C012212 --RKRSSSEM KKKDRHSKIC TSKGPRDRRM RLSLEIARNF FDLQDMLGFD KASKTVEWLF TKSRSAIKEV KEKYLSEVK- ---------- ----------
MELO3C017286 DLISQKKRSS SKRDRHSKIN TLHGPRDRRM RLSLPVAKEF FGLQDMLGVD KASKTVEWLL FQARHAIKKL SKDQQSFHID G--------- ----------
MELO3C002076 -----QRKRL SKKDRHSKIT TAKGVRDRRM RLSLPVAKQF FGLQDMLGVD KGSKTVECNS SSNYSNSSEL DNEAGVIIGD Q--------- ----------
```

```
                      310        320        330        340        350        360        370        380        390        400
              ....|....| ....|....| ....|....| ....|....| ....|....| ....|....| ....|....| ....|....| ....|....| ....|....|
MELO3C001972  ---------- ---------- ---------- -----NFNSL NTSLRRSATS ISLPSQLRSS SYYHP----- ---------- ---------- ---------G
MELO3C017168  ---------- ---------- ---------- -----NFTSL NISLRSSGSS MSVPSQLRSS SYYNPNFSLH QQRRTLFPG- ------FGLS SE--TSSTAA
MELO3C012086  ---------- ---------- ---------- -----NFTSL NISLRSSGST ISAPSHLR-T SYFSPTQFAV RTRSDLWDR- ------TNNH NHNLVDDSST
MELO3C011835  ---------- ---------- ---------- -----NFSSL NISIRSSGST LSAPPSKSAP HSFHGAALAL AHHPHAYEEG FAHAALLGFH HQQQQQQQHQ
MELO3C011754  ---------- ---------- ---------- -----NFSSV SLSVRGNGGG SASLSSPSST TSTLS----- ---------- ---------- ----------
MELO3C025357  ---------- ---------- ---------- -----SFSTI SASLR----- ---------T PPAAS----- ---------- ---------- ----------
MELO3C025049  ---------- ---------- ---------- -----SFSTI SASLR----- ---------T PPAAS----- ---------- ---------- ----------
MELO3C006863  ---------- ---------- ---------- -----IAVSV GGTLKIP-TT SPARPNGEIS EAPRK----- ---------- ---------- ----------
MELO3C022520  ---------- ---------- ---------- -----IAMSV NGTLKIPTTT SSSNQDSADP SAVKK----- ---------- ---------- ----------
MELO3C024459  ---------- ---------- ---------- -----GPIST VSSAMASSG- -RSVSCRIQP VTVGG----- ---------- ---------- ----------
MELO3C014903  ---------- ---------- ---------- -----SALAT AAGSVSQQG- -ASLSAGLHQ KLEEL----- ---------- ---------- ----------
MELO3C022331  ---------- ---------- ---------- -----SALHA AGVSLSDHDS SVSVSASLTP KIEAV----- ---------- ---------- ----------
MELO3C026920  ---------- ---------- ---------- -----TLDLP GCSKPFDPSS SNINSTTTSY FEEET----- ---------- ---------- ----------
MELO3C017230  ---------- ---------- --------PH HHYYTSSSSS SSSRFCNSQP PPFMLDVKST DAYQTTMTVE KTKLWDMDLA VLPEKAGSAT TAATTTKGKW
MELO3C002038  ---------- ---------- ---------- ---------- NASILHHS-- -----DIGG- ---------- -------DAA LMKAK----- ----------
MELO3C010371  ---------- ---------- ---------- -LHYQSMLSP TTTVVHPHRS EFKIIDKAG- ---VEDETKQ KQHKSNSNPS HPNSSFSALL NNVSAPPFGF
MELO3C007121  VE-------- ---------- ----QKQSQL HVSLSKSACS SNSETSKGSG LSLSRSEVRV NRLKARERAK ERAQKEKEKE QDSSRITDHN LSSMTRNSSF
MELO3C002754  PEMELDG--- ---------D PKKHQQNPRQ QLSLTRSACS SNSETSKGSG LSLSRSEILV NRAKARDRAR ERTAKEKERE QES-----RD AHCIPNNPSF
MELO3C002636  ---------- ---------- ---------- ---------- ---------- ---------- ---------- ---------- ---------- ----------
MELO3C019745  ---------- ---------- --TTAPSSVP VETETNGIDV VVDRSECCKN HDFLFQSQFS ENLIP----- ------PPPP PPVESGVGFR ----------
MELO3C025629  IPVSIHPSEE SATRISNRRA NFVVGDGGIS ECTMQNLQNI STEDNHNSDN SNFLPPSFDS DSIVDTFKSF LPV-TPTAAA ETPSSIFEFD TFPPDLLSRT
MELO3C014426  ---------- ---------- ---------- ---------- ---------- ---------- ---------- ---------- ---------- ----------
MELO3C016092  NLLSVQR--D VFNSAGNRRA TVLGSDSRVS EFPLQNLQHT QMGEGPNSST SSFLPPSLDS DSIADTIKSF FPIGTAAAAA ETTSSSIQFQ NYPQDLLSRT
MELO3C022091  ---------- ---------- ---------- ----NNIITS SSSSKFHFDH DFEECEVISN DDDDEE---- ---------- ---------- ---AEKIIML
MELO3C023772  ---------- ---------- ---------- -AKKLSLAAA SSDVEEEYDD DKGWELKMKS MLRID----- ---------- ---------- ----------
MELO3C005358  ---------- ---------- ---------- -----SNSGQ SSEVVSEIND NNGVVNVAAS NHLDQ----- ---------- ---------- ----------
MELO3C012212  ---------- ---------- ---------- --------SK SSCYSSTDLV TNLCKEVLPS GKKRR----- ---------- ---------- ----------
MELO3C017286  ---------- ---------- ---------- -----NGDTR SPSSVSDGEV VSGIIDETPT VVNNN----- ---------- ---------- ----------
MELO3C002076  ---------- ---------- ---------- -----NLNTK DNNIKINKKK IKKKKIMRAK QLPIR----- ---------- ---------- -------KTM

                      410        420        430        440        450        460        470        480        490        500
              ....|....| ....|....| ....|....| ....|....| ....|....| ....|....| ....|....| ....|....| ....|....| ....|....|
MELO3C001972  VMGFNSEAMN N-MVQITTQL ---------- ---------- ---PKEDHQY HLHQS----- -----TTTGV VPPTN----I PANFWMLSDP NNQLLASGVD
MELO3C017168  LLNFQSTNIG NSLLQAKPEI RDTPSSLDLS DAAEEISIGR KRRPSTEQEL SSSSSQHQMG GYLLQSSTGT IPASHGGAQV PANFWMLTNT NNQVMGG--D
MELO3C012086  FLNFNSSSYV NVFKQDQGGI D--------- ---------- --VVETGKKR RSEQSESSSE NYMVQTSTGS IPASQS-SQL PATFWMVTNP TNQ--GS--D
MELO3C011835  LNLMTADQIG EGLPSGGGGE STDNYMRKRF REDLFKEDTQ SPGESSGGGG SPKAMKSDLQ LEKQQQQQPS ASSSSGLLRP GAAMWAVATP GPSSGAS--N
MELO3C011754  LSRPQALSGP TPFILGKRVR S--------- ---------- -----DDD-- AKDD------ GLGVGQAVGS IVGPTG---- PGGYWAIPAR PDYG------
MELO3C025357  VSDHKPLLPP APFILGKRVR T--------- ---------- -----DDDVA NKDDTGGAGA GISVGPSIGS IMGPAV---- AGGYWAIPAR SDFG------
MELO3C025049  VFDHKPLLPP APFILGKRVR T--------- ---------- -----DDDEA NKDDTGGDGP GISVGPSIGS IMGPAV---- AGGYWAIPAR SDFG------
MELO3C006863  -RRRKGPNSE SNDCNDQASV S--------- ---------- --------SG LTPIAP---- MAAYGAGLVP FWGSAGGVTE P--FFMVPGT SNN----HQP
MELO3C022520  -RRKRPANSD YVDVNDALSV S--------- ---------- --------IS ASNVAGSPAV QQAFPPGFVP VWAIPSNAII PGAVFMVPSV ACSS---TRP
MELO3C024459  -SGQGMFSMP PPSCRLDLCQ P--------- ---------- ---------- ---------- ---------- ---------- ---------V GMEYS-----
MELO3C014903  -GGG--GSSR ANWGRPHLAA TP-------- ---------- ------TALW PSSVTGFGFQ SSTAAAAVAA QSATNLPNET SNYFHKLGFP PFDLTPTNMG
MELO3C022331  -PRTNWAMMG ANLSRSHMAT P--------- ---------- -------GFW P-SLGGIG-- ----TGFVSE NPGSIMP--- -----KFGFH GFEFQGMNLG
MELO3C026920  -----LGGNN YAFPTVNECI P--------- ---------- ---------- ---------- ---------- ---------- ---------- ----------
MELO3C017230  IDGSNNDSNS TQQVEDYNEQ ISSQ------ -----KLLSM AICSSNLPNM MNNNNSTPYN NYNSSHDHHH HRQQPSCLSL SQFGNNNNSN GIFPSQLVEA
MELO3C002038  ----NEETDQ TLLVQT---- ---------- ---------- --------ND INGDSS---- ---------- HFEPPS---- ---------- --FP------
MELO3C010371  YWDPNPPSSS SSTTNN---- ---------- --------NL PFLTSQAGDN NNNNNLHTFN -------HLN LSLPPSPLSL STASPFLEFS QLHQNFFIN-
MELO3C007121  TELLAG---- -GAASVSTHR DAG------- -------VAA ERQWQSSTVA MDYFSS-GIL EP-----STS RTHH--SSGF SDQMNLGTSL PQSMSSTPLF
MELO3C002754  TELLTGGINN NGANNTTNRT NSAS------ -----QNNVV EPNLFDKATA MDYLASSGII AQQPSSSSSS RPGHNQSAGF SGQIFLGNPH QLPVS-IPQF
MELO3C002636  ----NDCRKQ IFSLPVPSS- ---------- ---------- ---------- ---------- ---------- ---------- ---------- ----------
MELO3C019745  --SCRNLGLS LQTQNSPHNV H--------- ---------- -----HQTVF YTANS----- ---------G NSGPGFDDG- FEKIVAWNTN ----------
MELO3C025629  SSRTQDLRLS LQSLHGPTSK LESE------ -----QTQ-- ----QNDNLY FSGTTPLGCF D----GWSEQ QPSPTMEISR FQRILNWSTV SADHSGGGDG
MELO3C014426  ---------- ---------- ---------- ---------- ---------- ---------- ---------- ---------- ---------- ----------
MELO3C016092  SSQNQDLRLS LQSFQDPIAI HRHH------ -----HAQHQ SQGHQNEHVL FSGTAPLSGF DVTTAGWTEH NSLNPAEISR FPRITSWNAS GAETGSGGGG
MELO3C022091  GKSSSNCKKN LGFHDVLAKE S--------- ---------- ------RAKA RARARERTKE KMIMNQQFRS PPPPPPQPSM PTPVVQKVAG SDQENNNIIG
MELO3C023772  -EQEKVSKEK VEIFNLVAKE S--------- ---------- ------RAKA RARARERTME KKQVDDSKVY GHQKGGAQEV SDHWSKHLNH STETSNLSME
MELO3C005358  QQDQEDVAFT GGFKDKIKSR ---------- ---------- ---------- ---------- ---------- ---------- ---------- ----------
MELO3C012212  -RRFRIVDKE SRYKARVRAR ER-------- ---------- -----TRAKS IRRGLEVSKS DLEGNPNGVR KLGDVSEENS SNKSKISTHV SLCEN-----
MELO3C017286  --DINRKELE IGRKATTKKE K--------- ---------- ------RSRV GRKMAFHPFT RECREKARAR ARARAREKQL QIEGTSTTTK LQDVS-----
MELO3C002076  AKELREKARE RARARTLEKQ K--------- ---------- ------NQLF IRDSACCSYD HQPNFTNINS SSSWSSPFET TGGEESAGTT HQSYN-----

                      510        520        530        540        550        560        570        580        590        600
              ....|....| ....|....| ....|....| ....|....| ....|....| ....|....| ....|....| ....|....| ....|....| ....|....|
MELO3C001972  PLWRLQSANQ DGSLFRSG-- -----QSSPS G--GLRLSNN I-------LG PAPLTLLPGK PLSLGPTAGA ARRGNLNDHD DDDDDDDGHL NIYAAQNPPH
MELO3C017168  PIWTFPSVNN SG-LYRGT-- -----MSS-- ---GLHFMN- ---------F PAPVALVPGQ --QFGSSTGG G--SNNNNNN NNSSSTEGHL NILAGLN-PY
MELO3C012086  QMWTFPSVGN SS-MLRGT-- -----GGGGG GGGGFHFMN- ---------L PPPMALLPSG Q-QLGTGIGG G--------- -GGPATDSHL GMLAALSAYR
MELO3C011835  TFWMLPVTAG SGGGNVGNSS GGGSGGGGGG GGGGLEAHQM WPFSTSGNTL QAPLHLMPRF NIPGGVEIQA AAAGG----G RGGGGQLQLG SMLMQQATGG
MELO3C011754  QVWSFAAAAP S--------- ---------- ---EMVVQQG G--------I SQQASLFAQQ QQPIREASAA R--------- -VGNYLPGHL NLLASLSGG-
MELO3C025357  QVWSFAAAAA AA-------- ---------- AAPEMVIQPT A--------V SHQASLFVQQ Q-SMGEASAA K--------- -VGNYLPGHL NLLASLSGGP
MELO3C025049  QLWSFAAAAA A--------- ---------- --PEMVIQPT A--------V SHQASLFVQQ Q-SMGEASAA K--------- -VGNYLPGHL NLLASLSGGP
MELO3C006863  QLWAVPARPL SNLVSSMN-- ---------- --PGLQFGGV V-------PV LTRAVSNGSS GLESGSSPAM V--------- -SPSLIPGTV SAPASTSG--
MELO3C022520  QLFSFSAPAA PVINTTAR-- ---------- --PISAFPSK P-------DT SHIQSSPTNQ SSSTSSSSAS K--------- -SQSVMAPIS SSSVQRSGKT
MELO3C024459  ------AAGN D--------- ---------- ---------- ---------- ---YRHMPFT ALLLQPSTG- ---------- --EETEERQE EEVFRE----
MELO3C014903  PMSFTSILGA N--------- ---------- ---------- ---------- ---TQQLPGL ELGLSQDGH- ---------- -IGVINPQAL NQIYQQMGQP
MELO3C022331  SVNFPGMIGN Q--------- ---------- ---------- ---------- ---HQQIPGL ELGLSQDGNN L--------- -RGMLNPPSF SQIYHQIGQN
MELO3C026920  ---------- ---------- ---------- ---------- ---------- ------LPNY EFDLVSDFDM E--------- ----LFAHHI ATLQEVAENS
MELO3C017230  AQASTSSSSS SSSLFFTPYA SYITNPFESH IHFLSSNSHA LSNNPFMSSS LHPFATSQNL KPFPPPSFNL K--------- --QLFHAADT NNNAGGSAGA
MELO3C002038  ---------- ---------- ---------- ---------- ---------- -FPFMYQTNS SPF---SF-- ---------- ---------- ----------
MELO3C010371  ---------- ---------- ---------- ---------- -------NNN NNNPSFHPNV RPFH-FSMAT K--------- --FLSHQDKN NNNVDPTSKD
MELO3C007121  SSVSTGDSNA EQLHQFSFVH DGNIVPVATT QPGGGNDYSL NFTISSNLPG YYRGTLQSNS S--LLPHLQR F--------- --SPVDGSNL PFLFGAA--T
MELO3C002754  NISAAENTQQ DRLQHFSFVP DN-------- ----LNDYNL NFTISAGLAG CNRGTLQSNI SPSLLPYLQR F--------- --PHLEGSNV PFFIGAASPT
MELO3C002636  ---------- ---------- ---------- ----SSSSQ- -----FQSYP LQNFSQKGNL SVFSPMILNV T--------- --TAGFSGTQ PPCPSQSNVG
MELO3C019745  ---------- ---------- --VFGSIPIV N---QSSTA- -----FPQRG PLQSSLESTV PHYTWNDLSI V--------- --SAGKRHPN PIQENPSPSI
MELO3C025629  KAGGVDGEFL YNSQPTSSSF PPPSPPLPIL QPLFGENQL- -----VSQRG PLQSSYTPSI RAWIDPSIAF M--------- --DNQQQLIP PSIYQSSFPG
MELO3C014426  ---------- ---------- ---------- ---------- ---------- ---------- ---------- ---------- ---------- ----------
MELO3C016092  --GIRSAGYV FN----SPHL PTALPPSQLM QPLFGENQF- -----FSQRG PLQSSNTPSI RAWIDPSLTP T--------- --DHHQHQMP PSIHQSSYAG
MELO3C022091  IMWKPSPMVT SS-------- ---------- ---------- --------SS YQKNLVISKG ESCNYNNYCN N--------- -FYFPASNLT PNWDINIDST
MELO3C023772  ESSFMNKRKI YA-------- ---------- ---------- --------KK FINHDNYTTV KSCRDDENAE TS-------- HWKLLDQMKA SKRKWKPSII
MELO3C005358  ---------- ---------- ---------- ---------- ---------- -TLRTGSREA RDRARARARQ R--------- --TLLKNTLL PNPPITSSIS
MELO3C012212  ---------- ---------- ---------- ---------- ---------- --EKDLGSEE KNSDSLIMDK R--------- ---VSKEICS EDIHLLPDSS
MELO3C017286  ---------- ---------- ---------- ---------- ---------- -KISLWMSST QMENNGNDEQ LR-------- --TRNEGKII IDHETTDDCL
MELO3C002076  ---------- ---------- ---------- ---------- ---------- ----NNNHPI TSQFDSQFHQ N--------- --FSSFDAIL PKWSPSDDST
```

```
                      610        620        630        640
             ....|....| ....|....| ....|....| ....|....| ....
MELO3C001972 GAAPDTGVS- ---------- ---------- ---------- ----
MELO3C017168 RSVSNTGVME PQGSGSQSHH GGGGDDRHDT TSHQS----- ----
MELO3C012086 TTMPGLSVSE SPTRGVGGVT SNNPDQS--- ---------- ----
MELO3C011835 GGGGGLGVSE TNLGMLAALN AAYNSRSGSG FKYQFR---- ----
MELO3C011754 GGSGRREDDP R--------- ---------- ---------- ----
MELO3C025357 GSSGRRDNDH R--------- ---------- ---------- ----
MELO3C025049 GSSGRRDNDH R--------- ---------- ---------- ----
MELO3C006863 STQMLRDFSL EIYEKKELEF MGRPSPAKSK TPCSKP---- ----
MELO3C022520 TTQMLRDFSL EIYDKEELQF MTRSS---SK H--------- ----
MELO3C024459 ---------- ---------- ---------- ---------- ----
MELO3C014903 RVHQHHPQ-- -QPPAKDDSQ DSA------- ---------- ----
MELO3C022331 RESSAPPLEQ DQAPEKDDSQ ESR------- ---------- ----
MELO3C026920 EQETEDGDD- ---------- ---------- ---------- ----
MELO3C017230 GSSAKDGAAG NS-------- ---------- ---------- ----
MELO3C002038 ---------- ---------- ---------- ---------- ----
MELO3C010371 HGFASHQ--- ---------- ---------- ---------- ----
MELO3C007121 SAAPQLENHN HYQFSPA-FD GRLQLCYGGG NRQSEQKGKG KD--
MELO3C002754 TATTPVQLEN HHQLTPALFD GSWQLCYADG SHHSDQKGKG KN--
MELO3C002636 AGFPFFPMWE ---------- ---------- ---------- ----
MELO3C019745 PRFISNELPH LDVHGRFQND GDGDDDGNGN GGKASSRSSS YSRR
MELO3C025629 LGFATGGFSR FLIPTRIP-- --GEEEHDGI SEKPSS-ASS NSRH
MELO3C014426 ---------- ---------- ---------- ---------- ----
MELO3C016092 LGFASGGFSG FHIPTRIQ-- --GEEEHDGI SDKPSS-ASS DSRH
MELO3C022091 FARTQSGFCS IANMDSLPEF QVCAKPWDSC NNNQHLH--- ----
MELO3C023772 SGSSQRNLLI SIDDIPINLP RNLDTNNSAG YRFAK----- ----
MELO3C005358 QHFDHLSVLK LP-------- ---------- ---------- ----
MELO3C012212 MDMPSLSLDT MKASSAGLKL LELDEVDGVL IFYGE----- ----
MELO3C017286 MIMGRWSPSN SIYCNSLNNN NNGSPQEVNS KHQQT----- ----
MELO3C002076 FNFLHQQVYG LVASMLQVNN MGFYSQFSQG N--------- ----
```
