## Supplementary File 2 for "Genome-Wide Analysis of *TCP* Family Genes and Their Constitutive Expression Pattern Analysis in the Melon (*Cucumis melo*)"

### Cucumber TCP Proteins

**>Cucsa.014000**

GNQSSNNAATPLPVKKPPSKDRHSKVDGRGRRIRMPIVCAARVFQLTRELGHKSDGQTIEWLLRQAEPSIIAATG
TGTTPASFSTISASLRTPPAASLSDHKPLLPPAPFILGKRVRTDDDGANKDDTGGAGAGISVGPSIGSIMGPAVA
GGYWAIPARSDFGQVWSFAAAAAAAAAPEMVIQPTAVSHQASLFVQQQSMGEASAAKVGNYLPGHLNLLASLSGG
PGSSGRRDNDHR

**>Cucsa.031600**

MVIPSAPGSGTTGMGKKDRHSKIYTAQGLRDRRVRLSIDIARKFFDLQDMLGYDKASKTLEWLFSKSKKAIKELS
RTKSGNIGVHGGAKKFSLVADSSDVEEEYDDDKEGWELKMKSMLSIDEQEKVSKEKVEGFNLLAKESRAKARARA
RERTMEKKQVDNRKVYGHQKGGAQEVSNHWSKHLNHSTETSNLSMEESSFINKRKIIYSKKFINHDNYSDKSRRD
DENAETSQRKLLDQMKASKRKWKPSIISGSSQRNLLISIDGIPINLTQNLDTNSNPNYPIAK

**>Cucsa.047860**

MEVVDNIQEQPRKFPRLANGRSSDSNEIGLKGVENIHTGDEKDKESKRDGGITAGVGGGGLGGADDETNRLRGWH
HSRIIRVSRASGGKDRHSKVWTSKGLRDRRVRLSVATAIQFYDLQDRLGFEQPSKAVEWLIKAAADAIKELPSLN
ASFPETPKQLSGEKISVTDRPGPLDSVEQKHSQLHVSLSKSACSSNSETSKGSGLSLSRSEVRVNRLKARERAKE
RAQKEKEKEQDSSRITDHNLSSMTRNSSFTELLAGGAASVSAHRDAGVAAERQWQSSTVAMDYFSSGILEPSTSR
THHSSGFSDQMNLGTSLPQTMSSTPLFSSVSTGDSNAEQLHQFSFVHDGNIVPVATTQPGGGNDYSLNFTISSNL
PGYYRGTLQSNSSLLPHLQRFSPVDGSNLPFLFGAATSAAPQLENHNHYQFSPAFDGRLQLCYGGGNRQSEQKGK
GKD

**>Cucsa.054000**

MGENKPAEIKDFQIAIAEKEEGKKQLAPKRSSNKDRHTKVEGRGRRIRMPALCAARIFQLTRELDHKSDGETIQW
LLQQAEPSIIAATGTGTIPASALAAAAGSVSQQGASLSAGLHQKLEELGGGGSSRANWGRPHLAATPTALWPSSV
TGFGFQSSTAAAAVAAQSATNLPNETSNYFHKLGFPPFDLTPTNMGPMSFTSILGANTQQLPGLELGLSQDGHIG
VLNPQALNQIYQQMGQPRVHQHHHQQPPAKDDSQDSA

**>Cucsa.076020**

MELTDMQQQSNKQQQQQQQHHHHHHLHHQNQTQKLGSSQSQSQSQSQSPSPSPSHGHPFDARSPPSSAGGGGGGA
GGAGGGGHPTFMASTSASSAATHHLDASLVIATRSDTTAAADPSPRSTSTANAANKRSTKDRHTKVEGRGRRIRM
PATCAARVFQLTRELGHKSDGETIEWLLQQAEPAIIAATGTGTIPANFSSLNISIRSSGSTLSAPPSKSAPHSFH
GAALALAHHPHAYEEGFAHAALLGFHHQQQQQQQHHLNLMTADQIGEGLPSGGGESTDNYMRKRFREDLFKEDTQ
SPGESSGGGGSPKAMKTDLQLEKQQQQPSASSSGLLRPGAAMWAVAAPGPSSGASNTFWMLPVTAGSGGGNVGNS
SGGGGGGGGGGGGGLEAHQMWPFSTSGNTLQAPLHLMPRFNIPGGVEIQAAAAAAAAAAAGGGRGGGGQLQLGS
MLMQQAAGGGGGGLGVSETNLGMLAALNAAYNSRSGSGFNINSDNHPLEHQQHQHQHHHHHHHHHNPAHNTNTDH
NSGDEAPNTSQ

**>Cucsa.111880**

MEVLDIQEQACRFPRIGNGRNDSNKLGQKGIVQYTPDDEEDKEFSKRVAVVTSATVNPTGRCEPGGIGGVGANRL
HGWHHSSRIIRVARASGGKDRHSKVWTSKGLRDRRVRLSVATAIQFYDLQDRLGFEQPSKAVEWLIEAASDAIAE
LPSLSGSFPETPRQLSDEKMGSDGADQGFDSPEMELDGDPKKHQQNPRQQLSLARSACSSNSETSKGSGLSLSRS
EILVNRAKARDRARERTAKEKEREQESRDAHCIPNNPSFTELLTGGINNNGANNTTNRTNSASQNNVVEPNLFDK
ATAMDYLASSGIIAQQPSSSSSSRPGHNQSAGFSGQIFLGNPHQLPVSIPQFNISAAENTQQDRLQHFSFVPDNL
NDYNLNFTISAGLAGCNRGTLQSNISPSFLPYLQRFPHLEGSNVPFFIGAASPTTATTPVQLENHHQLTPALFDG
SWQLCYADGSHHSDQKGKGKN

**>Cucsa.112620**

MEGNDQLHYHQNHRSNASFPLQLLEKKEYDPQPPLEIVSKKPSPKRTSTKDRHTKVDGRGRRTRMPALCAARVFQ
LTRELGHKSDGETIEWLLQQAEPAVIAATGTGTIPANFNSLNTSLRRSATSISLPSQLRSSSYYHPGVMGFNSEA
MNNMVQITTQLPKEDHQYHLHQSTTTGVVPPSTIPANFWMLSDPNNQLLASGVDPLWRLQSANQDGSLFRSGQPS
PSGGLRLSNNILGGPAPLTLLPGKPLSLGPTAGAARRGNLNNDHDDDDDDHNGHLNIYATQNPPHGAAPDTGVSQ
CIHGHHLSIIKSID

**>Cucsa.113300**

MKKARTCSRQGQFLRGNPRIFRVSPFFGGKDRHSKVCTIKGLRDRRIRLSIPTAIQLYDLQNKLGLSQPSKVIDW
LIDVTRFEIDKLPPLPFPKDFDPNASILHHSDIGDAAFKAKNEETDHTLL

**>Cucsa.113660**

SGNTKNVHDPFGNLSHDLDHSFHLPISFFPFPSPFEVDDDRDLNFTTNSPSNSSQNYNNNPPPIISTYQKMKKMM
KKVPQRKRLSKKDRHSKITTAKGVRDRRMRLSLPVAKQFFGLQDMLGVDKGSKTVEWLLIQAKPEILKLATEKKN

HNCFTISNSSSNYSNSSELDNEVGVIIGDQNLNTKDNNIKISKKKIKKKKIMRAKQLPIRKTMAKELREKARERA
RARTLEKLKSQMFIRDSACCSDDHQPNFTNINSSSSWSSPFETTGGEESAGTTTHQSNNNNNPITSQFDSQFHQI
FSSFDFILPKWSPSDDSTFNFLHQQVSFEVNLLLFFLFNLNQTVYGLVASMLQVSTTWDFDS
**>Cucsa.122730**
MGEFSNQRPAKSAKPGIRTGGDGGGSEIVEVQGGRVVRSIGRKDRHSKVCTAKGPRDRRVRLSAHTAIEFYDVQD
RLGYDRPSKAVDWLIKKAKPAIDKLRELPGWNPNVLDMSTQKLTKQAEKNSENKIPVSIHPSEESATRISNRRAN
FMVGDGGISKCTMQNLQNISTEDNHNSDNSNFLPPSFDSDSIVDTFKSFLPVTTAAAAETPSSIFEFDTFPPDLL
SRTSSRTQDLRLSLQSLQGPTSKLESEQTQQNDHLYFSGTTPLGCFDGWSEQQPPPTMEISRFQRILQWSTVSAD
HSGGGDDKAGGVDGEFLYNSQPTSSIFPPPSPPLPILQPLFGENQLVSQRGPLQSSYTPSIRAWIDPSIAFMDNQ
QQLSPPSIYQSSFSGLGFATGGFSKFLIPTRIAGEEEHDGISEKPSSASSNSRH
**>Cucsa.124600**
MGSDIVLHKFPSSSDPHSQPPTASHSVVGSNSSPVTITRQSLLPRKSNTPLSSSSSKDRHIKVNGRGRRVRMPAL
CAARIFQLTRELGHRSEGETIEWLLRQAEPSIIAATGTGTVPSGPISTVSSAMASSGRSVSCRVQPVSVGGSGQG
MFAMPPPSCRLDLCQPVGMEYSAAGNDYRHMPFTALLLQPSTAEETEERQEEEVFRE
**>Cucsa.127020**
NTTKIARRNRRLSSTADAGAGVGKVYTSKGLRARRLRLSAPTAIKFYDLQDRLGCGRPTEAIDWLLLNAKSAIDA
LSRPLQTEDNDCRKQSFSLPVPSSSSSSSSSSSSSQFQSYPLQNFSQKV
**>Cucsa.166230**
MSNSDHTNGVSNGATIDTHKNHSSSNGALVVKKPPSKDRHSKVDGRGRRIRMPIICAARVFQLTRELGHKSDGQT
IEWLLRQAEPSIIAATGTGTTPANFSSVSLSVRGNGGGSASLSSPSSTTSTLSLSRPQPLSGPTPFILGKRVRSD
DDAKDDALGVGQAVGSIVGPTGPGGYWAIPARPDYGQVWSFAAAAPSEMVVQPGGIAQQASLFAQQQQPIREASA
ARVGNYLPGHLNLLASLSGGGGSGRREDDPR
**>Cucsa.197650**
MEGGDELHHHHHHHHQNHHHQHQHQHQHQHQHRPTFPFQLLEKKDDDAASCSTSPYPSLAISADTTTTTTTTTT
TSSVVAAETSKKPPPKRTSTKDRHTKVDGRGRRIRMPALCAARVFQLTRELGHKSDGETIEWLLQQAEPAVIAAT
GTGTIPANFTSLNISLRSSGSSMSVPSQLRSSSYYNPNFSLHQQRRTLFPGFGLSSETSSTAALLNFQSTNIGNS
LLQAKPEIRDTPSSLDLSDAAEEISIGRKRRPSTEQELSSSSSQHQMGSYLLQSSTGTIPASHGGAQVPANFWML
TNTNNQVMGGDPIWTFPSVNNSGLYRGTMSSGLHFMNFPAPVALVPGQQFGSGTGGGSNNNNNNNNNNSSSEGHL
NILAGLNPYRSVSSSGVMEPQGSGSQSHHGGGGGGDDRHDTTSHQS
**>Cucsa.198300**
MLRNNSKDQKGCEAKEEGVLVRSSSNNNNDGKYSFPKAPTSSSRQWSGFRNPRIVRVSRSFGGKDRHSKVCTIRG
LRDRRIRLSVPTAIQLYDLQDKLKLGQPSKVIDWLLDVTKLDIDNLPPLQIPPGFSPFHQQMLFPQYVDPHHHYY
TSSSSSSSSSSSSSRFCNSLPPPFMLDVKSTDAYQTTMTVEKTKLWDMDLAVLQDKAGSATTAATTTKGKWIDGS
NNDSNSTQQVEDYNEQISSQKLLSMAICSSNLPNMMNNNNSTPYNNYNSSHDHNHRQQPSCLSLSQFGNNNNSNG
VFPSQLVETAAQGSNSSSSSSSLFFTPYASYITNPFESHIHFLSSNSHALSNPFMSSSLHPFATSQNLKPFPPPS
FNLKQLFHAADTNNAGGSAGAGSSAKDGAAGNS
**>Cucsa.198860**
MFVFNSCSNNNHGNDCISFPDHSFLHFPSPFDDDGNPTNSLLLQQQSQHDIFLHHHIPLNEPSPPPPSSTFVNAL
RSSETINNDVFHQDLVSQRKKSSSKRDRHSKINTLHGPRDRRMRLSLPVAKEFFGLQDMLGVDKASKTVEWLLFQ
ARHAIKKLSKDQQSFHIDGNGDTRSPSSVSDGEVVSGIIDETSTVVNNNDMISTKELEIGRKSTTKKEKRSRVGR
KMPFNPLTRECREKARARARARAREKQQQIKGTSTTTKLQDVSKISSPWMSSTQMENNGIDEQLRTRNEGRIIMD
HETTDDCLIMGRWSPSNSIYCNSLHNNNNGSPQEHPYGDFQFLVKSWEPYNNHSIC
**>Cucsa.215290**
MPCGNKKRGFFFSLKKKNQYAGTRFEASCSIINLEEQLLKAQEHTSGYQCLCVKNSPPPKLAAGSSSSSATPWLR
LKDPRIVRVSRAFGGKDRHSKVCTIRGLRDRRVRLSVSTAIQLYDLQDRLGLNQPSKVVDWLLNAAKDEIDELPP
LPIPSASLGLHYQSMIPTTTTVVHPHRSEFKIIDKAGVEDETEQKQHKSNSNPSHPNSSFSALLNNVSAPPFGFY
WDHNPPSSSSSTTNNNLPFLTSQAGDNNNNNNLHTFNHLNLSLPPSPLSLSTASQFLEFNHLHQNFFINNNSNNN
NPSFHPNVRPFHFSMATKFLPHQDKNNNNVDPPX
**>Cucsa.238090**
MFSSTNNLHLFPLQQYFPSSSSSSPSYHHHLVAPPPSPPPHHHHSEPVYIGPLQDPVLSGVVAPIFLQVQEEDRD
QARMKNEQQEVAVLMSQNGNNNNNNNNGQCFSSSTTLNNFTKKGSVKKDRHSKIYTAQGLRDRRVRLSIDISRKFF
DLQDMLGYDKASKTLDWLLTKSRKAIKELTKKTQYHNNNNNIITSSSSSSKFHFDHDFEECEVISNDDDDEEAEK
IIMLGKSSSNCKKNLGFHDVLAKESRAKARARARERTKEKMIMNQQFRSPPPPPPQPSMPTPVVQKAAGPDQENN

NIIGIMWKPSPMVTSSSSYQKNLVISKGESCNYNNYCNNFYFPASNLTPNWDINIDSTFARPQSGFCSIANMDSL
PEFQVCAKPWDSCNNNQHLH

**>Cucsa.251110**

MPPLTLSQVPSTTLAKQTKRPSKDRHTKVEGRGRRIRMPAACAARIFQLTRELDHKSDGETIRWLLEHAEPAIIE
ATGTGTVPAIAVSVGGTLKIPTTSPARPNGEISEAPRKRRRKGTNSESNDCNDQASVSSGLTPIAPMAAYGAGLV
PFWGSAGGVTEPFFMVPGTSNNHQPQLWAVPARPLSNLVSSMNPGLQFGGVVPVLTRAVSNGSSGLESGSSPAMV
SASLIPGTVSAPASTSGSTQMLRDFSLEIYEKKELEFMGRRSPAKSQTPCSKP

**>Cucsa.273460**

MGESHRQTSTSSRLGMRTGGGGGEIVEVQGGHIVRSTGRKDRHSKVCTAKGPRDRRVRLSAHTAIQFYDVQDRLG
YDRPSKAVDWLIKKAKAAIDELAELPAWHPSTGNASTQGEEQQSLNDENENLLSVQRDVFNSTGPNSSTSSFLPP
SLDSDSIADTIKSFFPIGTSAAAAETTSSSIQFQNYPQDLLSRTSSQNQDLRLSLQSFQDPIAIHRHHHAQHQSQ
GHQNEHVLFSGTAPLSGFDVTTAGWSEHNSLNPAEISRFPRITSWNASGAETGSGGGGGIRSAGYVFNSPHLPTA
LPPSQLMQPLFGENQFFSQRGPLQSSNTPSIRAWIDPSLTHTDHQQHQIPPSIHQSSYAGLGFASGGFSGFHIPT
RIQGEEEHDGISDKPSSASSDSRH

**>Cucsa.280880**

MDDHQQQQHQEDQILINSYDDHDFHYFPHPFLDDDDLFLSHFLSQQQQFFEQRDQQLDETTKKMKKKKKLCGSSS
SRKNGSNSTKRRGKKDRHSKIYTAQGPRDRRMRLSLQIARKFFDLQDMLGFDKASKTIEWLLLNSNSAIKDLKQA
YFFKYSNSGQSSEVVSEINDNNGVFNVAASNLYQEQDQEDVAFTGGFKDKIKSRTLRTVAREARDRARARARQRT
LLKNTLLPNPPITSSISQHFDHLSX

**>Cucsa.283930**

MSRLFEQQIHPPSLQLIDCSNYNYNNNYFNPFLDDHQEFILGHILYSQKHQLLANNSDQIGVIETKTDHSSQKIP
ISRKRSSSVMKKKDRHSKICTSKGPRDRRMRLSLEIARNFFDLQDMLGFDKASKTVEWLFTKSRSAIKEVKEKYL
SEAKSKSSCYSSTDLVTNLYKEVLPSGKKRRRRLRIVDKESRYKARARARERTRAKSIQRGLEVSKSNLEGNPDG
VRKMGDFSEEISGNKRKTSTHVSLCENEKVLGSEEKNSDSLIMDKRVSKEICSDEIHLLPDSSMDMPTLSLDTMK
ASSAGLKLLELDEVDGVLIFYGE

**>Cucsa.285170**

MIMDGENGIRRPNFPLQLLEKKKEEEEEQQQHPCSSSANNGGTKTTTTITTTPNSNLQSSSSPKKPPPKRATTKD
RHTKVDGRGRRIRMPALCAARVFQLTRELGHKSDGETIEWLLQQAEPAVIAATGTGTIPANFTSLNISLRSSGST
ISAPSHLRTSYFSPTQFAVRTRSDLWDRTNNHNHNLVDDSSTFLNFNSSNYVNVFKQDQGGIDVVETGKKRRSEQ
SESSSENYMVQTSTGSIPASQSSQLPATFWMVTNPTNQGSDQMWTFPSVGNSSMLRGAGGGGGGGGGGGFHFMNL
PPPMALLPSGQQLGTGIGGGGGGPATDSHLGMLAALSAYRTTMPGLSVSESPTRGVGGVTSNNPDQS

**>Cucsa.349500**

MADQNNNNNNKHHPLLPSPATSSSPLTTPPLPTTSSDIKDFTLIISNKDQNNSNSNNNINNKQLAPKRTSNKDRH
KKVDGRGRRIRMPALCAARVFQLTRELGHKTDGETIQWLLQQAEPSIIAATGSGTIPASALHAVGVSLSDHDSSV
SVSASLTPKIEAVPRTNWAMMGANLSRSHMASQGFWPSLGGIGTGFVSENPGSIMPKFGFHGFELQGMNLGSVNF
PAMIGNQHQQIPGLELGLSQDGNNLRGMLNPPSFSQIYHQIGQNRDSSAQPLEQDQAPEKDDSQESR

**>Cucsa.363020**

RPSTKDRHTKVEGRGRRIRIPATCAARIFQLTRELGHKSDGETIRWLLERAEPAIIAATGTGTIPAIAMSVNGTL
KIPTTTSSSNQDSADPSAVKKRRKRPANSDYVDVNDALSVSISASNVAGSSAVQQAFPPGFVPVWAIPSNAIIPG
AVFMMWAKGGPLGLQEEQQQQQQQQHHVVVPSPHIPGSCMGGG

**>Cucsa.384890**

MDSNKPFLNSTLEQHSTPTTTMGINTIPGQNIHIPGGGLVVRSTARKDRHSKVYTSRGLRDRRVRLSAHTAIQFY
DVQDRLGFDRPSKAVDWLIKKAKSAIDKLSADLPPCNTNACFSVPVETQTNGIDVVTVFYTASSGNSGPGFDDGF
EKIVAWNTNVFGSIPILNQSSTAFPQRGPLQSRLESTVPHYTWNDLSVVAAGKRHPNPIQENSSPSIPRFISNEL
PHLDVHGQFQNDGDGKDNGGKSSSHSSSYSRR

**>Cucsa.385930**

MRVKRPTSNSPRNAPKPNSPIASESKLIPSKAPKDRHAKVHGRDRRIRLPPLCAARVFQLTRELGNKTDGQTVEW
LLKKAEPSIIALTGKNIASATLDLPGCSKPCDPSSSNTNSTTTSYFEEETLGGNNYAFPM

#### Arabidopsis TCP Proteins

**>AT1G30210**

MEVDEDIELQKHQEQQSRKLQRFSEDNTGLMRNWNNPSSRIIRVSRASGGKDRHSKVLTSKGLRDRRIRLSVATA
IQFYDLQDRLGFDQPSKAVEWLINAASDSITDLPLLNTNFDHLDQNQNQTKSACSSGTSESSLLSLSRTEIRGKA
RERARERTAKDRDKDLQNAHSSFTQLLTGGFDQQPSNRNWTGGSDCFNPVQLQIPNSSSQEPMNHPFSFVPDYNF
GISSSSSAINGGYSSRGTLQSNSQSLFLNNNNNITQRSSISSSSSSSSPMDSQSISFFMATPPPLDHHNHQLPET
FDGRLYLYYGEGNRSSDDKAKERR

**>AT1G35560**

MESHNNNQSNNNTTGSAHLVPSMGPISGSVSLTTTAPNSTTTTVTAAKTPAKRPSKDRHIKVDGRGRRIRMPAIC
AARVFQLTRELQHKSDGETIEWLLQQAEPAIIAATGTGTIPANISTLNISLRSSGSTLSAPLSKSFHMGRAAQNA
AVFGFQQQLYHPHHITTDSSSSSLPKTFREEDLFKDPNFLDQEPGSRSPKPGSEAPDQDPGSTRSRTQNMIPPMW
ALAPTPASTNGGSAFWMLPVGGGGGPANVQDPSQHMWAFNPGHYPGRIGSVQLGSMLVGGQQLGLGVAENNNLGL
FSGGGGDGGRVGLGMSLEQKPQHQVSDHATRDQNPTIDGSP

**>AT1G53230**

MAPDNDHFLDSPSPPLLEMRHHQSATENGGGCGEIVEVQGGHIVRSTGRKDRHSKVCTAKGPRDRRVRLSAPTAI
QFYDVQDRLGFDRPSKAVDWLITKAKSAIDDLAQLPPWNPADTLRQHAAAAANAKPRKTKTLISPPPPQPEETEH
HRIGEEEDNESSFLPASMDSDSIADTIKSFFPVASTQQSYHHQPPSRGNTQNQDLLRLSLQSFQNGPPFPNQTEP
ALFSGQSNNQLAFDSSTASWEQSHQSPEFGKIQRLVSWNNVGAAESAGSTGGFVFASPSSLHPVYSQSQLLSQRG
PLQSINTPMIRAWFDPHHHHHHHQQSMTTDDLHHHHPYHIPPGIHQSAIPGIAFASSGEFSGFRIPARFQGEQEE
HGGDNKPSSASSDSRH

**>AT1G58100**

MDLSDIRNNNNDTAAVATGGGARQLVDASLSIVPRSTPPEDSTLATTSSTATATTTKRSTKDRHTKVDGRGRRIR
MPALCAARVFQLTRELGHKSDGETIEWLLQQAEPAIVAATGTGTIPANFSTLSVSLRSSGSTLSAPPSKSVPLYG
ALGLTHHQYDEQGGGGVFAAHTSPLLGFHHQLQHHQNQNQNQDPVETIPEGENFSRKRYRSVDLSKENDDRKQNE
NKSLKESETSGPTAAPMWAVAPPSRSGAGNTFWMLPVPTTAGNQMESSSNNNTAAGHRAPPMWPFVNSAGGGAGG
GGGAATHFMAGTGFSFPMDQYRGSPLQLGSFLAQPQPTQNLGLSMPDSNLGMLAALNSAYSRGGNANANAEQANN
AVEHQEKQQQSDHDDDSREENSNSSE

**>AT1G67260**

MSSSTNDYNDGNNNGVYPLSLYLSSLSGHQDIIHNPYNHQLKASPGHMVSAVPESLIDYMAFKSNNVVNQQGFEF
PEVSKEIKKVVKKDRHSKIQTAQGIRDRRVRLSIGIARQFFDLQDMLGFDKASKTLDWLLKKSRKAIKEVVQAKN
LNNDDEDFGNIGGDVEQEEEKEEDDNGDKSFVYGLSPGYGEEEVVCEATKAGIRKKKSELRNISSKGLGAKARGK
AKERTKEMMAYDNPETASDITQSEIMDPFKRSIVFNEGEDMTHLFYKEPIEEFDNQESILTNMTLPTKMGQSYNQ
NNGILMLVDQSSSSNYNTFLPQNLDYSYDQNPFHDQTLYVVTDKNFPKGKVWIQDSFVN

**>AT1G68800**

MFPSLDTNGYDLFDPFIPHQTTMFPSFITHIQSPNSHHHYSSPSFPFSSDFLESFDESFLINQFLLQQQDVAANV
VESPWKFCKKLELKKKNEKCVDGSTSQEVQWRRTVKKRDRHSKICTAQGPRDRRMRLSLQIARKFFDLQDMLGFD
KASKTIEWLFSKSKTSIKQLKERVAASEGGGKDEHLQVDEKEKDETLKLRVSKRRTKTMESSFKTKESRERARKR
ARERTMAKMKMRLFETSETISDPHQETREIKITNGVQLLEKENKEQEWSNTNDVHMVEYQMDSVSIIEKFLGLTS
DSSSSSIFGDSEECYTSLSSVRGMSTPREHNTTSIATVDEEKSPISSFSLYDYLCY

**>AT1G69690**

MDPDPDHNHRPNFPLQLLDSSTSSSSTSLAIISTTSEPNSEPKKPPPKRTSTKDRHTKVEGRGRRIRMPAMCAAR
VFQLTRELGHKSDGETIEWLLQQAEPAVIAATGTGTIPANFTSLNISLRSSRSSLSAAHLRTTPSSYYFHSPHQS
MTHHLQHQHQVRPKNESHSSSSSSSQLLDHNQMGNYLVQSTAGSLPTSQSPATAPFWSSGDNTQNLWAFNINPHH
SGVVAGDVYNPNSGGSGGGSGVHLMNFAAPIALFSGQPLASGYGGGGGGGGEHSHYGVLAALNAAYRPVAETGNH
NNNQQNRDGDHHHNHQEDGSTSHHS

**>AT1G72010**

MNQNSSVAEATLQLNSGEKPSPGSIPFISSGQHGNISTSATSSTSTSSGSALAVVKSAVKKPTKDRHTKVDGRGR
RIRMPAMCAARVFQLTRELGHKSDGETIEWLLQQAEPAIIASTGTGTIPANFSTLNASLRSGGGSTLFSQASKSS
SSPLSFHSTGMSLYEDNNGTNGSSVDPSRKLLNSAANAAVFGFHHQMYPPIMSTERNPNTLVKPYREDYFKEPSS
AAEPSESSQKASQFQEQELAQGRGTANVVPQPMWAVAPGTTNGGSAFWMLPMSGSGGREQMQQQPGHQMWAFNPG
NYPVGTGRVVTAPMGSMMLGGQQLGLGVAEGNMAAAMRGSRGDGLAMTLDQHQHQLQHQEPNQSQASENGGDDKK

**>AT2G31070**

MGLKGYSVGEGGGEIVEVQGGHIIRATGRKDRHSKVFTSKGPRDRRVRLSAHTAIQFYDVQDRLGYDRPSKAVDW
LIKKAKTAIDKLELGETTTTTTRQEPVNTKPESPTLVFQRENNDQTQFVAANLDPEDAMKTFFPATTTTNGGGGT
NINFQNYPHQDDNNMVSRTTTPPPNLSQDLGLSLHPFQGNNNTVVVPETNNFTTTHFDTFGRISGWNHHDLTMTS
SSSSEHQQQEQEERSNGGFMVNHHPHHHHHQPSMMTLLNSQQQQVFLGGQQQQQQRGTLQSSLFPHSFRSWDHHQ
TTSDHHHHQNQASSMFASSSQYGSHGMMMMQGLSFPNTTRLLHGEEATQPNSSSSPPNSHL
**>AT2G37000**
MIFQNVCRNESNFNAIASESRSQTQFGVSKSSSSGGGCISARTKDRHTKVNGRSRRVTMPALAAARIFQLTRELG
HKTEGETIEWLLSQAEPSIIAATGYGTKLISNWVDVAADDSSSSSSMTSPQTQTQTPQSPSCRLDLCQPIGIQYP
VNGYSHMPFTAMLLEPMTTTAESEVEIAEEEERRRRHH
**>AT2G45680**
MATIQKLEEVAGKDQTLRAVDLTIINGVRNVETSRPFQVNPTVSLEPKAEPVMPSFSMSLAPPSSTGPPLKRAST
KDRHTKVEGRGRRIRMPATCAARIFQLTRELGHKSDGETIRWLLENAEPAIIAATGTGTVPAIAMSVNGTLKIPT
TTNADSDMGENLMKKKRKRPSNSEYIDISDAVSASSGLAPIATTTTIQPPQALASSTVAQQLLPQGMYPMWAIPS
NAMIPTVGAFFLIPQIAGPSNQPQLLAFPAAAASPSSYVAAVQQASTMARPPPLQVVPSSGFVSVSDVSGSNLSR
ATSVMAPSSSSGVTTGSSSSIATTTTHTLRDFSLEIYEKQELHQFMSTTTARSSNH
**>AT3G02150**
MNIVSWKDANDEVAGGATTRREREVKEDQEETEVRATSGKTVIKKQPTSISSSSSSWMKSKDPRIVRVSRAFGGK
DRHSKVCTLRGLRDRRVRLSVPTAIQLYDLQERLGVDQPSKAVDWLLDAAKEEIDELPPLPISPENFSIFNHHQS
FLNLGQRPGQDPTQLGFKINGCVQKSTTTSREENDREKGENDVVYTNNHHVGSYGTYHNLEHHHHHHQHLSLQAD
YHSHQLHSLVPFPSQILVCPMTTSPTTTTIQSLFPSSSSAGSGTMETLDPRQM
**>AT3G15030**
MSDDQFHHPPPPSSMRHRSTSDAADGGCGEIVEVQGGHIVRSTGRKDRHSKVCTAKGPRDRRVRLSAHTAIQFYD
VQDRLGFDRPSKAVDWLIKKAKTSIDELAELPPWNPADAIRLAAANAKPRRTTAKTQISPSPPPPQQQQQQQQLQ
FGVGFNGGGAEHPSNNESSFLPPSMDSDSIADTIKSFFPVIGSSTEAPSNHNLMHNYHHQHPPDLLSRTNSQNQD
LRLSLQSFPDGPPSLLHHQHHHHTSASASEPTLFYGQSNPLGFDTSSWEQQSSEFGRIQRLVAWNSGGGGGATDT
GNGGGFLFAPPTPSTTSFQPVLGQSQQLYSQRGPLQSSYSPMIRAWFDPHHHHQSISTDDLNHHHHLPPPVHQSA
IPGIGFASGEFSSGFRIPARFQGQEEEQHDGLTHKPSSASSISRH
**>AT3G18550**
MNNNIFSTTTTINDDYMLFPYNDHYSSQPLLPFSPSSSINDILIHSTSNTSNNHLDHHHQFQQPSPFSHFEFAPD
CALLTSFHPENNGHDDNQTIPNDNHHPSLHFPLNNTIVEQPTEPSETINLIEDSQRISTSQDPKMKKAKKPSRTD
RHSKIKTAKGTRDRRMRLSLDVAKELFGLQDMLGFDKASKTVEWLLTQAKPEIIKIATTLSHHGCFSSGDESHIR
PVLGSMDTSSDLCELASMWTVDDRGSNTNTTETRGNKVDGRSMRGKRKRPEPRTPILKKLSKEERAKARERAKGR
TMEKMMMKMKGRSQLVKVVEEDAHDHGEIIKNNNRSQVNRSSFEMTHCEDKIEELCKNDRFAVCNEFIMNKKDHI
SNESYDLVNYKPNSSFPVINHHRSQGAANSIEQHQFTDLHYSFGAKPRDLMHNYQNMY
**>AT3G27010**
MDPKNLNRHQVPNFLNPPPPPRNQGLVDDDAASAVVSDENRKPTTEIKDFQIVVSASDKEPNKKSQNQNQLGPKR
SSNKDRHTKVEGRGRRIRMPALCAARIFQLTRELGHKSDGETIQWLLQQAEPSIIAATGSGTIPASALASSAATS
NHHQGGSLTAGLMISHDLDGGSSSSGRPLNWGIGGGEGVSRSSLPTGLWPNVAGFGSGVPTTGLMSEGAGYRIGF
PGFDFPGVGHMSFASILGGNHNQMPGLELGLSQEGNVGVLNPQSFTQIYQQMGQAQAQAQGRVLHHMHHNHEEHQ
QESGEKDDSQGSGR
**>AT3G45150**
MDSKNGINNSQKARRTPKDRHLKIGGRDRRIRIPPSVAPQLFRLTKELGFKTDGETVSWLLQNAEPAIFAATGHG
VTTTSNEDIQPNRNFPSYTFNGDNISNNVFPCTVVNTGHRQMVFPVSTMTDHAPSTNYSTISDNYNSTFNGNATA
SDTTSAATTTATTTV
**>AT3G47620**
MQKPTSSILNVIMDGGDSVGGGGGDDHHRHLHHHHRPTFPFQLLGKHDPDDNHQQQPSPSSSSSLFSLHQHQQLS
QSQPQSQSQKSQPQTTQKELLQTQEESAVVAAKKPPLKRASTKDRHTKVDGRGRRIRMPALCAARVFQLTRELGH
KSDGETIEWLLQQAEPSVIAATGTGTIPANFTSLNISLRSSGSSMSLPSHFRSAASTFSPNNIFSPAMLQQQQQQ
QRGGGVGFHHPHLQGRAPTSSLFPGIDNFTPTTSFLNFHNPTKQEGDQDSEELNSEKKRRIQTTSDLHQQQQQHQ
HDQIGGYTLQSSNSGSTATAAAAQQIPGNFWMVAAAAAAGGGGGNNNQTGGLMTASIGTGGGGGEPVWTFPSINT
AAAALYRSGVSGVPSGAVSSGLHFMNFAAPMAFLTGQQQLATTSNHEINEDSNNNEGGRSDGGGDHHNTQRHHHH
QQQHHHNILSGLNQYGRQVSGDSQASGSLGGGDEEDQQD
**>AT4G18390**

MIGDLMKNNNNGDVVDNEVNNRLSRWHHNSSRIIRVSRASGGKDRHSKVLTSKGPRDRRVRLSVSTALQFYDLQD
RLGYDQPSKAVEWLIKAAEDSISELPSLNNTHFPTDDENHQNQTLTTVAANSLSKSACSSNSDTSKNSSGLSLSR
SELRDKARERARERTAKETKERDHNHTSFTDLLNSGSDPVNSNRQWMASAPSSSPMEYFSSGLILGSGQQTHFPI
STNSHPFSSISDHHHHHPHHQHQEFSFVPDHLISPAESNGGAFNLDFNMSTPSGAGAAVSAASGGGFSGFNRGTL
QSNSTNQHQSFLANLQRFPTSESGGGPQFLFGALPAENHHHNHQFQLYYENGCRNSSEHKGKGKN

**>AT5G08070**

MGIKKEDQKSSLSLLTQRWNNPRIVRVSRAFGGKDRHSKVCTVRGLRDRRIRLSVMTAIQVYDLQERLGLSQPSK
VIDWLLEVAKNDVDLLPPLQFPPGFHQLNPNLTGLGESFPGVFDLGRTQREALDLEKRKWVNLDHVFDHIDHHNH
FSNSIQSNKLYFPTITSSSSSYHYNLGHLQQSLLDQSGNVTVAFSNNYNNNNLNPPAAETMSSLFPTRYPSFLGG
GQLQLFSSTSSQPDHIE

**>AT5G08330**

MADNDGAVSNGIIVEQTSNKGPLNAVKKPPSKDRHSKVDGRGRRIRMPIICAARVFQLTRELGHKSDGQTIEWLL
RQAEPSIIAATGTGTTPASFSTASLSTSSPFTLGKRVVRAEEGESGGGGGGGLTVGHTMGTSLMGGGGSGGFWAV
PARPDFGQVWSFATGAPPEMVFAQQQQPATLFVRHQQQQQASAAAAAAMGEASAARVGNYLPGHHLNLLASLSGG
ANGSGRREDDHEPR

**>AT5G23280**

MSINNNNNNNNNNNDGLMISSNGALIEQQPSVVVKKPPAKDRHSKVDGRGRRIRMPIICAARVFQLTRELGHKSD
GQTIEWLLRQAEPSIIAATGTGTTPASFSTASVSIRGATNSTSLDHKPTSLLGGTSPFILGKRVRADEDSNNSHN
HSSVGKDETFTTTPAGFWAVPARPDFGQVWSFAGAPQEMFLQQQHHHQQPLFVHQQQQQQAAMGEASAARVGNYL
PGHLNLLASLSGGSPGSDRREEDPR

**>AT5G41030**

MVMEPKKNQNLPSFLNPSRQNQDNDKKRKQTEVKGFDIVVGEKRKKKENEEEDQEIQILYEKEKKKPNKDRHLKV
EGRGRRVRLPPLCAARIYQLTKELGHKSDGETLEWLLQHAEPSILSATVNGIKPTESVVSQPPLTADLMICHSVE
EASRTQMEANGLWRNETGQTIGGFDLNYGIGFDFNGVPEIGFGDNQTPGLELRLSQVGVLNPQVFQQMGKEQFRV
LHHHSHEDQQQSAEENGS

**>AT5G51910**

MESNHEGNAIQVIDQVTTMTHLSDPNPKTKPGMMLMKQEDGYLQPVKTKPAPKRPTSKDRHTKVEGRGRRIRMPA
GCAARVFQLTRELGHKSDGETIRWLLERAEPAIIEATGTGTVPAIAVSVNGTLKIPTSSPVLNDGGRDGDGDLIK
KRRKRNCTSDFVDVNDSCHSSVTSGLAPITASNYGVNILNVNTQGFVPFWPMGMGTAFVTGGPDQMGQMWAIPTV
ATAPFLNVGARPVSSYVSNASDAEAEMETSGGGTTQPLRDFSLEIYDKRELQFLGGSGNSSPSSCHET

**>AT5G60970**

MRSGECDEEEIQAKQERDQNQNHQVNLNHMLQQQQPSSVSSSRQWTSAFRNPRIVRVSRTFGGKDRHSKVCTVRG
LRDRRIRLSVPTAIQLYDLQDRLGLSQPSKVIDWLLEAAKDDVDKLPPLQFPHGFNQMYPNLIFGNSGFGESPSS
TTSTTFPGTNLGFLENWDLGGSSRTRARLTDTTTTQRESFDLDKGKWIKNDENSNQDHQGFNTNHQQQFPLTNPY
NNTSAYYNLGHLQQSLDQSGNNVTVAISNVAANNNNNLNLHPPSSSAGDGSQLFFGPTPPAMSSLFPTYPSFLGA
SHHHHVVDGAGHLQLFSSNSNTASQQHMMPGNTSLIRPFHHLMSSNHDTDHHSSDNESDS

**Rice (Indica) TCP Proteins**

**>BGIOSGA000770**

MILGSNQAAAAAAAAAAAEEEAAELARKHTAAVATSRQWSAQTESRIVRVSRVFGGKDRHSKVKTVKGLRDRRVR
LSVPTAIQLYDLQDRLGLNQPSKVVDWLLNAARHEIDKLPPLQFPPQDHLCMGHHHHLPSAMPLMHHHGHHHHAD
DDKYHVAAAAAALAAEKEAAAAGGGGGGGGDDVDGGGGGGAAHIVGRFPAGGYHRFMGLNNPLGMVNSAAGAAMP
FHYAGESWNNGSVQDSGAGSPQVAAAAAHHTSPFPSLLSLAPGPHHQLVFYSSEAEQFTVDNLGSQGLSLSSARA
FHDQTGS

**>BGIOSGA002128**

METRPPAAPAKLSYGIRRGWTRIGAATVAAGKKAAGDLDPRHHHHRVTHGGDGGGVGGGGSGGQEEADEQQQQQ
HDHHRLLQLHHHQGVQQDQEPPPVPVFHLQPASVRQLSGSSAEYALLSPMGDAGGHSHHHQHGFQPQLLSFGGVG
HHHHLHQFTAQPQPPAASHTRGRGGGGEIVPATTTPRSRGGGGGGGGEIVAVQGGHIVRSTGRKDRHSKVCTARG
PRDRRVRLSAHTAIQFYDVQDRLGYDRPSKAVDWLIKNAKDAIDKLDVLPAWQPTAGGAGAGNAAAPPSSSTHPD
SAENSDDQAQAITVAHTAFDFAGGGSGGTSFLPPSLDSDAIADTIKSFFPMGGTAGGEASSSTTAAQSSAMGFQS
YTPDLLSRTGSQSQELRLSLQSLPDPMFHHQQHRHGGGGGGGNGTTQQALFSGAANYSFGGGAMWATEQQAQNQR
MLPWNVPDPGGGGGAAYLFNVSQQAAHMQAAAAALGGHQSQFFFQRGPLQSSNQPSERGWPETVEADNQMSHHQG
GLSPSVSAAIGFAAPGIGFSGFRLPARIQGDEEHNGGGGGNGDKPPPPSSVSSASHH

**>BGIOSGA005085**

MDPKFPPPPPLNKTEPTTTTTNQQHHHDEQQQQHRLQIQVHPQQQEQQDGGGGGGKDQQQQQQMQVVVAAAAGER
RMQGLGPKRSSNKDRHTKVDGRGRRIRMPALCAARIFQLTRELGHKSDGETVQWLLQQAEPAIVAATGTGTIPAS
ALASVAPSLPSPNSALSRSHHHHHHMWAAAPPTASAGFAGAGFSGADSGVIGGIMQRMGIPAGIELQGGGAGGLG
GGGGGGGGHIGFAPMFAGHAAAAAAMPGLELGLSQDGHIGVLAAQSLSQFYHQVGAAGQLQHQHQHHHQQQQQQQ
EDGEDDRDDGESDEESGQ

**>BGIOSGA005319**

MTTRPDGGGGGGGAPDKQLVPASNANGTALAVRKPPSKDRHSKVDGRGRRIRMPIICAARVFQLTRELGHKSDGQ
TIEWLLRQAEPSIIAATGTGTTPASFSTSSPSSLRSNSTSNSNDLLLPRAAPFILGQAPPCRRRPHHFARARSRR
HRPHPGLLGAPGQG

**>BGIOSGA008683**

MSSRDAAATFHVYQPVQIPTATVAPAAAVSAAPAEAVAQLVPAPSKKAAGAAGGKDRHSKVNGRGRRVRMPIVCA
ARVFQLTRELGLKSDGQTIEWLLRQAEPAILAATGTGTTPAAFVSSSAPSTSSHQHTLLGKRQRQESAAADAVSV
AGAASAFWAALPAPGRPDAWGFSPLDAQPTYVPMAQAHHHHLNLLAALSGAARRAEEESR

**>BGIOSGA009049**

MDVAGDAGGGRRPNFPLQLLEKKEEQPCSSSAAGGGTGPSSAGGNGNNGSGPGGAGGEMQLRKAAPKRSSTKDRH
TKVEGRGRRIRMPALCAARVFQLTRELGHKTDGETIEWLLQQAEPAVIAATGTGTIPANFTSLNISLRSSGSSLS
APAHLRALPSPAADDRFGSRADAWDRVVSLGGESIWTFPQMSSAAAAAAVYRGSVPSGLHFMNFPAPMALLPGQQ
LGLGPVGGGGGGGGGGEGHMGILAALNAYRTQAATDAAGQQGGGGGGGGSSQQQHGGGGGGGERHQSISTSDS

**>BGIOSGA009051**

MEEVIVGKECKRPRYALVGVDGHGSDVADEPGGVRARRTSTTRLSVPTSIAFYDIQNRLGVNRPSKSIEWLICAA
LVARLFLRPPQHRPAPHPRRPSSGRPSEEQARTRKATVAASPGKSGEKRRKTRSGCAAWKAERIAKLRVWREGWM
PRREASTSAASGGQSWIRKLSTPGSGDSCSCSDAKPAKRRWAEKAGEVVAREAGGAERGGGGRRR

**>BGIOSGA013417**

MHPFMDLELEPHGQQLAAAEEDGAGGQGVDAGVPFGVDGAAAAAAAARKDRHSKISTAGGMRDRRMRLSLDVARK
FFALQDMLGFDKASKTVQWLLNMSKAAIREIMSDDASSVCEEDGSSSLSVDGKQQQHSNPADRGGGAGDHKGAAH
GHSDGKKPAKPRRAAANPKPPRRLANAHPVPDKESRAKARERARERTKEKNRMRWVTLASAISVEAATAAAAAGE
DKSPTSPSNNLNHSSSTNLVSTELEDGSSSTRHNGVGVSGGRMQEISAASEASDVIMAFANGGAYGDSGSYYLQQ
QHQQDQWELGGVVYANSRHYC

**>BGIOSGA013684**

MEAAVGDGEGGGGGGGRGKRGRGGGGGEMVEAVWGQTGSTASRIYRVRATGGKDRHSKVYTAKGIRDRRVRLSVA
TAIQFYDLQDRLGFDQPSKAIEWLINAASPAIDTLPSLDPAAFAAIPHAAAADAAPTRRRSQQQQQLSNKSGCS
STSETSKGSDKEVTVASAPAQAASFTELLIAGVAASSAGGGAIGNGADCVGIAHPGKGGAEGASTYGFSAASSFG
DAPPIGMVPAPPFNFSAPGADMAAHYSLAQDQLAAPPPPAGGDYNLNFSMSSGFLGANRGTLQSNSPSNMSGHHH
HHHQQQLQRLDGSTISFLLGHAAAAAHPAASEGQITSTAALQLWDGFRHSGMKEKSKN

**>BGIOSGA015954**

MVPAGVFQLTREFGHSTDGETIDCLLRQAEPSIIAATGTGVTPEEAPPAPVAIGSSSVAAAAAAAGHGGAFVHVP
YYTPLLMQPPNADEPPMASAASASGTTAADENNN

**>BGIOSGA016791**

MASRDAATFQVYRPMAMPTPAALPPSSQQITMPFTAAPVDAVLPAPRKAAATQGGKDRHSKVNGRGRRVRMPIVC
AARVFQLTRELGLKSDGQTIEWLLRQAEPSILAATGSGTTPAVFSCSSAPSTASSSFLLGKRPRQEDHEAPTFWE
ALQQQPRPAVSSWGALVSPSQEAQAYASSVAQVHHLNLLSALSGAATRRPAQEESR

**>BGIOSGA020173**

MACMAGVDSRGGEALSPAAVASGRGRSNHEQLKMISNNSTNEELGGGGAAAAVASSRHWSASTESRIVRVSRVFG
GKDRHSKVRTVKGLRDRRVRLSVPTAIQLYDLQDRLGLSQPSKVVDWLINAAQAEIDKLPPLQFPPHDHDLVAAA
ASSMAPPPFANGGDGHHGASASSMLEDGDKAAGGGGMKAFMSLSNSLGLLNAATMPATLAAHHHHHHAAAYYAA
AESWGNGGNGGHHHDVSHGVSPSAHNSPFPSLLSLAPGSHHQFVFYSPEGGGFAVKEAAAEQFPVDSLDHSQGQL
TLSSARSFLHSGSQG

**>BGIOSGA022577**

MDVTGDGGGGGQRPNFPLQLLGKKEEQTCSTSQTAGAGGGGVVGANGSAAAAPPKRTSTKDRHTKVDGRGRRIRM
PAICAARVFQLTRELGHKTDGETIEWLLQQAEPAVIAATGTGTIPANFTSLNISLRSSGSSLSIPSHLRLAGLAG
PRFGGGARAADAWDRVVGLGFGGAADAPSSATSSSSSPLLLSFHSGSVGLDVSPPSASTSPAAADLSRKRRWEQE
MQQQQQYQQQMAGYTQSQIPAGTVWMVPSSNAQAAGGGAPPGGGGESIWTFPQSGSGGGGGAATVYRGVPSGLHF
MNFPATPMALLPGGQQLGLAGAGGGGEGHPGILAALNAYRAQAAQPDAGAAAQNGAQGSSQHRQHQHHGGGGGGG
DERHESMSASDS

**>BGIOSGA025162**

MELAGSNNKRGRVRGPNDDDDDAGEPDAKRHHHQLLLPWPQQQQQQHPASRIYRVSRASGGKDRHSKVYTAKGIR
DRRVRLSVSTAIQFYDLQDRLGYDQPSKAIEWLIKAAAAAIDKLPSLDTASFPTNPASSAAVAAAAAPPLPHAER
EQQQQLTKSGCSSTSETSKGSVLSLSRSESRVKARERARERSSAAAAAASKDAGDDAATPTAPTAAPASSQAASF
TELLTGMAAANASPADHKQQQAWQPMTVAAATADYIGFAAAAAPHTQPRKSAAGHHSAMPHTFASPAPHLANITP
IAMAPAQHFTLTPAAAEHHAEMTHYSFDHFMPVHAAAAAAAAAASTPAGGDYNLNFSMSSGLVGVHSRGTLQSNSQ
SHLSSHHHHHHQQQQQQQQLQRLSAPLDAPNIPFLFSPAAAPTAADTQFAAALQLWDGFRHADIKEKGKH

**>BGIOSGA026548**

MEAQAQDKAEEGEEEGTRQQHAQAGPVGAAGGGGGGGAAAVAMSAIPMNSWLVPKPEPVEFFGGMAMVRKPPPRN
RDRHTKVEGRGRRIRMPAACAARIFQLTRELGHKSDGETIRWLLQQSEPAIIAATGTGTVPAIATTVDGVLRIPT
QSSSSSGPASSAVVDGEESSAKRRRKLQPTRAVAGASPLATAAPAAYYPVIADPLLQGSGGAAISVPSGLAPITA
TGAPQGLVPVFAVPATGSPAVAGGNRMIPQATAVWMVPQPAGAAGAGNQPTQFWAIQSAPQLVNFAGAQFPTAIN
VADFQQQQQQQPVSTTIVQNSNSGEHMHFSGADSHEQQRRGRKEGNSGGVVDHPEEDEDDDDDEPVSDSSPEE

**>BGIOSGA028717**

MSAPSSSSSPSTLDEYDARFFFPGADAYTAGHRQDEETLEAVLRQPVTTTAAVAAAAAAVEGGGGGGGGGAGGSP
AAAAAATRRRPFRTDRHSKIRTAQGVRDRRMRLSVGVARDFFALQDKLGFDKASRTVEWLLTQSKHAINRLTLPD
SADAAAAPAFAAAPPPADQHSSAMAAAAASAAKEKGEASSSSTTNASSARARNRDHDGSSPVAPMDERGRRGVEL
DWTAAAAASTEQPMDGLEYYFQYYNHLEEIMSCDPTTTTDE

**>BGIOSGA029420**

MASQLQGKGAETAAAGERVAPGTNAAAFAGLGYPPIQSPVALQEEEGPRDAAFAGYAPIRSPVVSRLQEKGEGEG
EEEEVDKREEAGMAADGSAFAAGMALVPKPEPVAVEFLRGLAVAKPPPRNRDRHVKVEGRGRRIRMPVNCAARIA
QLTRELGHKSDGETIRWLMQQSEPAIVAATGTGTVPAIATTVDGVLRIPTESPSAAARGDEPAPKRRRKLQPTRA
AAGGPVEALAAAPPPAVYYPIVADPLLQANGGGSISISSGLAPASSATPPTATGGGAIPFIAMPATSDGGKQAMS
PATVWMVPPGGAGAVNQPIQYWAFQPNPDHANFAGASSYNVGQNPGVHEASAADHAASTGGGGGGEDDEYEGMTD
SSSDEE

**>BGIOSGA030744**

MQQQLDQQQQQQHQYDHFFSGHGQFNSETLEAVLCRPPRGAAADPAVPAAAAAAVLTAARNGGGHGRARKRPF
RTDRHSKIRTAQGVRDRRMRLSLDVARDFFALQDRLGFDKASKTVDWLLTQSKPAIDRLAADPSSSSHRAAGDTR
MSSAERGGDHHMVAVGAAGSGKGDADKARGPRGGRSAPMELGCELGRLVPAPVLGEYYYELAEMMSNNTGGEGDD
DGDYDDDGDFLDDSGFLCHYRSKKKFSIKDIS

**>BGIOSGA034630**

MAAAWRTSQVARVVAGKKDKHSKVVTSRGLRDRCVRLSVPMAIAFYDIQDRLGVDQPSKSIEWLIRAAAAAIDAF
LSLDCSLVLPNAAQLLTRWRRRPSSGQPSEEQARTRKATVAVSPRKSGEKRRKTRSGCAVRSASRS

**>BGIOSGA037077**

MDHGGGGGGGGAAPSSSNSGGGSGGGGGGGGRENHPHHPFYYSGPAAAAAAAAAAQQQQQTFMGALAITPVVAEQ
PQGSSGGGEKKVVAPTTPAAAGAAATTTLAKRPSKDRHTKVDGRGRRIRMPALCAARLPFSCAAAIDEIMWRRVV
DGINQAKQSNQSFELLSVPVHCSASH

**>BGIOSGA037799**

MEEVVGGGKERKRPRGALVGVGGGGESAATAAAWRTSRVARAAAGGKDRHSKVVTSRGLRDRRVRLSVPTAIAFY
DIQDRLGVDQPSKAIEWLIRAAAAAIDALPSLDCSFALPAAASSPPPPAADDAEVSTSETSKSSVLSLANAPCDN
GGGAFAELLHCSNTNGSKPLQQQQQATLAYYAAAQSAHMAAPMSFEVMAMPPHLAFSQEQQQHATVAAFDRGTLQ
SNASLWPPPPQPPPSQHPFLLQRFAAAPAEVAGLPFFLAGGVGGAAAAAPAATTNGGERRLQLWDFKEERKT

**Tomato TCP Proteins**

**>Solyc00g084870**
MSVHLCMLSIFSIVNVNSETNGKTALAVVPTKKKNELSVSSSKDRHTKVNGRGRRVRMLALCAARVFQLTKELGH
RTDGETIEWLLRNAEPAIIAATGTGGGFFSMTPQSQPNCRLDLCQPSLEFSGNAYRHMPFTALLLQPVTADDGEE
KVAEEDEKQ

**>Solyc00g217560**
MISREVDPSDNNVIMIPKKSSLSSSSSWTRFKDPRIVRVSRAFGGKDRHSKVLTVKGLRDRRVRLSVPTALQVYD
LQDKLGLDQPSKVVDWLLNEAKHDIDELPPLQIRDQTGLPLTGLRKEEEETMVVSDEDRVKLDVRNLDLKGNSNS
NNNKNSFSFGLYKDNHSSQSGIFIY

**>Solyc01g008230**
MSTSVEQNGAVLLDSTTGSGGGVANSNGALTVKKPPAKDRHSKVDGRGRRIRMPIVCAARVFQLTRELGHKSDGQ
TIEWLLRQAEPSIIAATGTGTIPASFSTVSVSLRNSISSVTASAPVDHKLPSPSPSLLHPLISPAPFLLGKRLRS
EDDDNGSGDKDVVPSAGFWAVPARPDFGQVWSFAAPSPEISHSAAAAMNTQPSRFLQQQMEEASAGRVGNYFPIA
QGHLNLLASLSGSGSGPRRDDDGQ

**>Solyc01g103780**
MELTDLQNNNTNSNSKQQTTTNNTTPSPPLHHQQQHQYQLLSQHHHHQQQQQLQQQHGRSSVPFMTSISIQPNAQ
APITNSATNASPSTSSPTPPQPPHLVDASLAIATRSDTLVDPNKKSQLAIQQTQPVQPQQPPPKRQTKDRHTKVD
GRGRRIRMPAACAARVFQLTRELGHKSDGETIEWLLQQAEPAIIAATGTGTIPANFSTLNISLRSSGSTLSAPPS
KSAPHSFHSALALAAAHQTPHPFEEGFSQMLGFHHYTHLLTQNPIAESIPGGGGGGGSSGDGGGVGGQEGADNYL
RKRYREDLFKEEGSSNQQGEASGGSSSPSNKQFKGNSPQLPNKPNSQEGVAGPSSNMLRHTNVIPATGLWAVAPP
PTSGTTGGSPFWMLPVTSGQTLSAPMASTSGTTTLESQMWPFPMGSSNTPLHFINNNHLNILGLEWPKLI

**>Solyc02g065800**
MISREVDPSDNNVIMIPKKSSLSSSSSWTRFKDPRIVRVSRAFGGKDRHSKVLTVKGLRDRRVRLSVPTALQVYD
LQDKLGLDQPSKVVDWLLNEAKHDIDELPPLQIRDQTGLPLTGLRKEEEETMVVSDEDRVKLDVRNT

**>Solyc02g068200**
MDPKQANHNNIKPTHDQIKELQILKNDETNKVAAPKRKDRHTKVEGRGRRIRMPALCAARIFQLTRELGHKSDGE
TIQWLLQQAEPSIIAATGTGTIPASALAAAASVSQQGISVSAGLMIESGANIAGSGSSRSSNSRTNWPMICGNFG
RPHLATAGMWPAPAPVVTSFGFQSSSAPSSASLGSDSSNYYLQKIGFPGFDLPAATSMNPMCFTSILGGSNQQLP
GLELGLSQEGHLGVLNQIYQQARMQHPQQQHQQQQQSPEEDSQGSGH

**>Solyc02g077250**
MSEECTKCTSVIFIYFFPLGPRCMKMNHQFQVSDEEDGSEEEEEEEEEVFQENDIGNLQSHYQNQQMPQSLCKKP
EKWANFTVSEQELNKGTRRMKPKRAKTDVIEGHGGRIIRATGRKDRHSKVSTAKGPKDRRVRLSPNTAIQFYDVQ
DRLGYDRPSKAIDWLIKEAKAAIDALGEFPNNFHSTKLNPQKMQYSFDQEQSPEFSQENRGVPNSECGVQDNQQE
VNYDIPNLFSSSDGFKIPFLSDLQSHPHGHFLNFQSLQDDTVLSSGNHHQGRFFTTTSVNHFPSVLSQNQVFSHR
EPLQSSFFPLMSDPLSTQLETLSYGFSNDGFSGIISSASRIQGEEEQATFFTAS

**>Solyc02g089020**
MITREEKGNVVNDDGNKSKACTTSTSTTSWTRLKDPRIVRVSRAFGGKDRHSKVCTVRGLRDRRVRLSVPTAIQL
YDLQDRLGLNQPSKVVDWLLDAAKNEIDELPPLQIPPGSLNPNLYPMLGNAPQITTNKEGRINWDNNIPLEPPNN
NALGYNPNFFKWDPSNLSLAHSENSLSHQHDHHQNFNLMSSMPLTSHHQPVVYPPGLPQFFHQHNNNINVGASSS
SEFDPKEINFQMLSNNSGSENPLSTSTNSFRHSIDSTNQSIRPFHFLPSQNCDDTEPSN

**>Solyc02g089830**
MFPFGNSSNGGNPILHSSFLNNQILLHQHDLPTHHHYLAAANGHSIDSYATNNVAINNKSKKQVKKDRHTKILTS
QGHRDRRVRLSIGVARKFFDLQDMLGYDKPSKTLDWLFTKSKLAIEDLINDVSKKSTPLSIHNNNNNNNSECDED
MIVPLAKKAKQERDSRAKARARARERTIKKIWTQIAPNREATASHYNNSTRNWNHDDVNPTIMSSMDASTICCTS
LPIVQKAWDSYHGSQI

**>Solyc02g094290**
MKRQNTNNTMEMKDFQIGIAEKDEAKKHQLAPKRKSNKDRHTKVEGRGRRIRMPALCAARIFQLTRELGHKSDGE
TIQWLLQKAEPSIIAATGHGTIQASLYRRLDPLFRNRE

**>Solyc03g006800**
MARVENNQNIEQEDDEVNDCKIDPLLGDSHEYTIAAVGNVTDAEGVDPSPAASTDRVLLLKEEPEENDLRVSTSV
GMNMQLQKVEKQPVKRSSKDRHTKVEGRGRRIRMPAACAARIFQLTRELGHKSEGETIRWLLERAEPAIIAATGT
GTVPAIAVSVNGTLKIPTSNNEGESSRKRRKRAANSEFYEASNFAPVAPIAPQGLVPVWPVGSGNGLIPTTAFTG

GATFYMLPPGTNTTVAATGAQLWATPILNVPYAATGVSAACVSKSDNNGGKLSTVASAMPPSSSSTQMLRDFSLE
IYDKRELQFMVGSGSSEIDQTSSSKS

**>Solyc03g045030**

MFSASNSSTHDNPLPHYISSSFHTSSPFLGFTGNQILLHQYYQNQFSSHYYLAKNNEDYCDNSLRSFPMKKKSKK
RERSCGKILTAQGPRDRRIRLSINMARKFFDLQELLGFDKPSKTIDWLFTHSELALEELTNWSTHQTHRPKISGS
LSKSNQQQGFRKKSQKSKRSNTKRVKEKGKSTS

**>Solyc03g115010**

MNSSSGNNFETKQESNNDSRLASKTTSAPPSSTSRQWGANKNPRIVRVSRTFGGKDRHSKVCTIRGLRDRRIRLS
VPTAIQLYDLQDRLGLSQPSKVVDWLLEATKLDIDTLPPLPVPPEYFTRFQQPSHEEFSDVRAQWLNANTSYLEH
GRNKSIASNEESNQEDNMFTLGNQRSSNLPGMPFNSYNYQWDHQPNSNLSLGHFGHNHSFSSQIDDDQSHQNTSF
PLSLGSSSSPFPSSSQLYFNPITTTFQPIVPSPNYITNHHPIESHDPRANMNHFNFLSSTTSHNNQLVMPTLNLI
SSQMKPFSLNNDPRWMSSRDDEDDDDDDDDDNQPHEGRAI

**>Solyc03g116320**

MDGGGDDHLHQRHQHNHRSIATFPFQLLEKKEDEACSSSSNAAAAYTTSLAISNTDNTHTNPNNTPRSTISTLQI
SASGADTSKKPPPKRTSTKDRHTKVDGRGRRIRMPALCAARVFQLTRELGHKSDGETIEWLLQQAEPAVIAATGT
GTIPANFTSLNISLRSSGSSMSVPSQLRSSYFNPNFSLSQRRGLFPPGIGLSTDTSATTLLNFQSANLSSNIQLQ
TKPELRDNSIDLTESSPAEDNLSRKRRSDLDLEQQQHQQQQQQMGSYLLQSSTGTMPTSHSSIPANFWMVTNPIP
SNQVMGGDPIWPFQSVSNSGALYRGTMPSGLQFMNFPTSVALLPSQQLGGGSSGGGGGNNGLGEGQLGMFAGLNP
YRGGGGVSESQASGSHSHHGGGTDDRHDTTSHHS

**>Solyc03g119770**

MYPSSNYSPNISSSSSFFHINIPSPSMQYEPEFIQYFHDFQFIQPSYDQNTNIPAEEAADSDKLDKIEEDQSIIK
SCNNNKKDEKSSSSTSTIRRKNNKRTTSGSAGVGPSKKDRHSKINTAHGPRDRRMRLSLEIARKFFNLQDLLGFD
KASKTVEWLLTKSKSAVNDLVQKINKDKCSGSENPNIATVSSPSAESCEVIDESAATNTAETQKQQKKKVKSIRR
AIIHPVVAKESRKEARARARERTIIKKSLNDNTNNNNNGDQSMADEDLTRSLRSWNTTFEDHQSGIQGYNNNNNM
NVVDNFNLVDTSNWSPFMFNYHQINTEISQEHQFANFQYSGKLWEA

**>Solyc04g006980**

MFPSSNNHDTFSYTSKTYLERSFTYDHQNPSSSSRQEDNPFFLNFPSPFLDHNESPLSQILPQDHHVKEGNLTHL
SSETSKEEMSIEAKPSSKKRSLSTTPRKRTGKKDRHSKICTAQGVRDRRVRLSLHIARKFFDLQDMLGFDKASKT
IEWLFSKSNNAIKDLSENTPQKEYSDGNKIVINSNNSSSYEGKSDSFMSECEENSINELGKDKEKIMQNNPHKRE
SREKARARARERTKEKMMIKGLEKGNPSNMFDQLGSSRSNSGFLDQDSNNNSYNTSVNQEKGSPHEANSQSLEHH
FPIQNYLGGASNSSTIDVGNCFMSFHGNWEINYAPMKSTNYSTTFAGNPSSIYLAQQYQTLDQEKNLSSKHHRLE

**>Solyc04g009180**

MSTSAEGNGAIIDPQRQQQQAPTGVGTNGALTVKKPPVKDRHSKVDGRGRRIRMPIVCAARVFQLTRELGHKSDG
QTIEWLLRQAEPSIIAATGTGTIPASFSTVSVSVRNSTASLVSSLSAPLDQKSSQMISPAPFILGKRLRSDDENI
ENGNKDDVAVAAGAATAVGPTAGFWAVPARPDFGQIWSFAAAPPPEMMVPTSAAAAAAAAAALSSQSSRFFQQQM
GEASAARVGNYLPMTQGHLNLLASLSGPPQPSSGRRDDDGR

**>Solyc05g007420**

METGHHHHDNIGNSIQRLNFPLQLLEKRDEVDHTTTICSSSLQLHPYNPTSLHMMTTSSSDAPCVNHNQKSKTHQ
DKKQQPQQTKKPTTTKDRHTKVDGRGRRIRMPATCAARVFQLTRELGHKSDGETIEWLLQQAEPAVIAATGTGTI
PANYSSLNISLRSSRHHSASNYLAHNHNNFGHVYHDRNYFNGVGLFSSENNSFFPSGNLNMLQAKEELCDDHDDN
DNDNDNNTGRIKRRSEDQDLLQNNYHMSNMLQSSTYGSIPASHQIGQIPATTLYMMTNNNNNNNNNSSHHHDLNN
CSMWGSSSNNNDSIENSNIRGGILNFMNFHQPILGTRGDGGGTAAEGQWGMLTAAMNSYRQSGGHASGSSNQRNE
GDDHQHHS

**>Solyc05g009900**

MFPKSNIIHDPFSFTSQELLKQSYSTHDQNPNSPSKVVEDEDHPFFLNNFFPSPFLDDHELPLNQIFSQKHHQKQ
EASDNHDINQADPDNTIKDNHSVDNSRSTQLNTKIMGDQSSNPAISSKKRKLSAKPRRRTGKKDRHSKICTAQGV
RDRRMRLSLQIARKFFDLQDMLGFDKASNTIEWLFSKSKNAIKELSRNISQESNSSDQNNDDHRKLRKIGSSDKP
IRAKTREKTKEKMMIKLGHNKKGNQELDETNPMSTIDPKLGSNPKSLEHQFANVGIMERYLGGASYSSITSIFDY
DNNGVIKGNIDISDNCFNMGILENCSMTNEVQIPFSGNNPSSIYLDYSRFHQF

**>Solyc05g012840**

MKPKRAKTDVIEGHGGRIIRATGRKDRHSKVSTAKGPKDRRVRLSPNTAIQFYDVQDRLGYDRPSKAIDWLIKEA
KAAIDALGEFPNNFHSTKLNPKKMQYSFDQEQSPEFSQENRGVPNSECGVQDKQQEVNYDIPNLFSSSDGLKIPF
LSDLQSYPHGHFLNFQSLQDDTILSSGNHHQGSFFTTTSVNHFPSVLSQNQVFSHREPLQSSFFPLMSDPLSK

**>Solyc05g032780**

MNHQCQVSDEEDGSEEEEEEEEEVFQENDIGNLQSHYQNQQMPQSVCENPEKWANFTVSGQELNKGTRRLKPKRA
KTDVIEGHGGRIIRATGRKYRHSNVSTAKGPKDRRVRLSPNTAIQFYDVQDRLGYDRPCKAIDSLIKEAKVTIDA
LGEFPNNFHSTKLNPKKMQYSFDQEQSPEFSQENRGVPNSECGVQDKQQEVNYDIPNMFSLY

**>Solyc06g065190**

MEFRESNSKNNQDGGSNSNNNSNENNSNVISTKIVKKPSKDRHTKVDGRGRRIRMPALCAARVFQLTKELGHKSD
GETIEWLLQQAEPSIIAATGTGTIPANFSTLNVSLRSSGTTISAPPSKSAPLFIHGGAATMLGFHHQIPGNSFGQ
DPDENFMKKRYREDTTAASTSPSSSSAKPERTGVQGHEPDQESKPGSSNPSSYIPTPAMWAVGPAAGNVGNTFWM
LPGTSSTMQRIGGFELPGGGRFSPVQLGSMFLQQPQPVQQLGLGVTETNMGMLASMNAYNSSSSRGSGIDLGMNL
EQHHHHQNQPQGSDSGDENHKDSQS

**>Solyc06g069240**

MYPPSNNNCSPILSSLICQNIPSSPCMQYEHELYFQSFNHDNQYYFQQQQLVPSIDDLSPHILADSCTEIITKPS
NCNHELQGMEEGRGEKKGDDDVMSSRISGRISKNNKRSSNKDRHSKINTARGPRDRRMRLSLDAARKFFRLQDLL
GFDKASKTVEWLLTQSDSAIEELVAAKGNDAQVAQQTSCNTPTTTTGIGAICASNSISESCEVISGTDETSSNDK
NKETAQDEEKKKRKKVVNTARRAVLEPLTKESRNQARARARERTKSKKMSQTGKSKSLANDLNPSGSRRPANKTC
EEPGTHEELNFHQEKNTVDDCNFMVNGNWNPFTIFSYHEQYAGISNETCNFVESYGKARARNFEADKVPSSFCAG
NSKRIKFLLHFVLTRPQQKNVSGLYLLVNVT

**>Solyc06g069460**

MNSRKQETGSGVDTNNSKTTSTTTATTIPSSSSSRQSSSWGGGFKNPRIVRVSRSFGGKDRHSKVCTVKGLRDRR
IRLSVPTAIQLYDLQDRLGLSQPSKVVDWLIDATKDEIDKLPPLQIPPPSLSHFFSTKDHHNLDFVDTGLFGKDR
WIATTNDQESSNHLIFQSSNLGMLNTFSNSWEPNSNLSLGPFGNYQDQYQQQQNMPLIPSSSHHQQQQLYFCPTS
SSTTTLSSLVPPYNSHFHPNSIAPTLQLTTSPVKSSFSLQDHHQHNHKEGSGS

**>Solyc06g070900**

MEGGGGDDHLHHHHHNHQHHQHHQQYRPNNFPFQLLEKKEDEPCSSSSAANNINYPSLAISPSDTNTNINPNSND
LQITVASTETAKKPAPKRTSTKDRHTKVDGRGRRIRMPALCAARVFQLTRELGHKSDGETIEWLLQQAEPAVIAA
TGTGTIPANFTSLNISLRSSGSSMSVPSQLRSSYFNPNFSLSQRRSLFQGIGLSSDRSATTTLLNFQTGNSNLHQ
FQAKQEMRDNSLDLTETSIEESLSRKRRQDLDLQQEQQQNQQQQNEQQMGSYLLQSSSSGTMPTSHSSIPANFWM
LTNNNTQVLGGDPVWTFPSVNNSGAAAAALYRSTMSSGLHFMNFPTPVALLPTQQFGAGSNGSTLAGEGQLGMVT
GLNPYRPCSGVSESQASGSHSHHGGGGGGDDRHDSTSHHNS

**>Solyc07g053410**

MKSATGGGEIVQVEGGHILRSTGRKDRHSKVYTAKGPRDRRVRLAAHTAIQFYDVQDRLGYDRPSKAVDWLIKKA
KNAIDKLDELPPWNPNQIPGNTTEADALVLKQQPEYQLQRELEENNQSRVNNSSNSYLMQGGSGGGGGGGEVQQQ
SLGDTMKAFFPMNLGTSLMNFQNYPHEIMPRSSLQNEDLVIIMRMTILPFSQIKILKQIIQEWLVGIYYHKHFSI
KMLIMHLLKGNPFSPIFHI

**>Solyc07g062680**

MAETSRLGIRNTVGEIVEVQGGHIVRSTGRKDRHSKVCTAKGPRDRRVRLSAHTAIQFYDVQDRLGYDRPSKAVD
WLIKKAKPAIDELAELPAWKPTIGTASAAAATNTNLEQEQAQKQQEDNNFAFQQGNVSLFDNVAGPSSKRAIESN
TASFLPPSLESDAIADTIKSFFPMGSSTSANSSAMQFHSFQEPHMLSRANSQNQDLSLSLQFQDPILLHHQNQQA
QHHNQTNHREQEQVQGQAPAHFGGNTPLGFDTSGWSMHQRLRSWSDSREIGPGGSGGAVTGPGGYLFNSPPAPAL
LQQLFGQNQFFSQRGPLQSSNTSSVRAWMDPSAIAIASGDPSNHHQAALSMYPSTIPGYGFASEVGGFSGFRIPA
RIQGEEEEHDGISDKPSSASSDSRH

**>Solyc08g048370**

MSNKEDEQYDVGEVKKSGDLGGGIGKLYGWPTSRIVRVSRASGGKDRHSKVLTSKGLRDRRVRLSVNTAIQFYDL
QDRLGCDQPSKVVEWLLKAAAPSIAELPPLEDLQDTLQLSNEKRSSEHGFDSADVEMDDDLNYNNQQQPSCSNSE
TSKGSGLSLSRSDSRVKARERARERATEKVANHHRNMHPSSSFTELLTGGMSDNNNNKTSVNDDQNTPRQWSTNP
LEYFTDGQIYLGNTLRPVSPPMFSITGGPFSPILPLLCLVIRGTDHILVPQLDMTTSIQIRKEKKRSNFKFHLVI
LNGSSLKVPKEAPIYAFLCYRLIILY

**>Solyc08g048390**

MEEIRTDECKFPRMSNKEDEQYQQYDVDDVGEVKKSSGLGGIVKFYGRPSSRIVRVSRASGGKDRHSKVLTSKGL
RDRRVRLSVNTAIQFYDLQDRLGCDQPSKAVEWLLKAAAPSIAELPPLEAFPDTLQLSDEKKSSEQGFDSADVEM
DDDLHYNQQQQPCCSNSETSKGSGLSLSRSDSRLKARERAKERATEKEKEKENKSCIVGQHHQNMHPGSSFTELL
TGGMSDNNTSPNGGSIHQNTARQWSTNPLEYFTSGLLGPSATRGIDNQIYLGNPLQPLRPMFSITGEHRAELQNF

PFGGDNLVPGVTSSNTSTNYNEYNLNFSISSSSSSGFNRGTLQSNSSSTLPHYQRFSPTDGSYLGSTTEYDARLH
LFYGACVNLFFL

**>Solyc08g080150**

MSSFHDPDDDCGTSELSNGAAGDLNDQKMDGHAVADDKKLVFSAMKEEPFDSDHHTLVPSMPMPVATAAPPRRSS
TKDRHTKVEGRGRRIRIPATCAARIFQLTRELGHKSDGETVRWLLEQAEQSIIEATGTGTVPAIAVSVNGTLKIP
TSSPASNPEIDENHPQKRRRKSSNSEFIDVGSLNRNAANVSQFAPVTTTPATITTAPPPGFVPVWGMSNGGMMVP
SNAIWMIPATVPPAQVNVTNNNNVVLQYPQLWTPVFNLATRPIPSFVTAATINTNGPPAIMIAGPLSVNNGAATT
TGPKIGSNKSSMAPASLSSSTDKNNNNTNDVKPHMLRDFSLEIYDRKELQLMSRSGNHQQETQVTSSERS

**>Solyc09g008030**

MGSEIVVSGAISGVGNGSVNSETNGKTALAVVPTKKKNALSVSSSKDRHTKVNGRGRRVRMPALCAARVFQLTKE
LGHRTDGETIEWLLRNAEPAIIAATGTGTVPATQVTTTSENIPLSQSQPSVLAPLTRATPVSGFPVGGGFFSMTP
QSQPNCRLDLCQPSLEFSGNAYRHMPFTALLLQPVTADDGEEKVAEEDEKQ

**>Solyc10g008780**

MKHHLEVSNENEDKSSSSSQQDDDDDENKDEFFQEILQSDYQHMQQWPNYFTPPSSTLVPKKQNNTQCPSHKEET
KKVSLKRKPKRAKQDVMEVHGSRIVRATGRKDRHSKVSTATGPKDRRVRLSPNTAIQFYDVQDRLGYDRPSKAID
WLIKEAQRAINALEKAPFQVFSNKINSTNKALNVRNVENEVGQFALEQSWSADKKTELEELARRSSKLNATVQYS
TDQFPEGCYKEQSSKVCQANATQIVNSSANGSKIQSFCQIQTHPQAHLHSQTKRQSPNDAKIQSFCEYQTHPQSH
FHSQTKKQSPNVAKTLSFCEMQTPSQGHFHSQTKNQPTNDAKIQSFCQLQTHPQSHFQSETKNQSPNVAKIQSFG
ELQTTPQSHFHSQTQKQSTNDAKIQSFCELQTHPQGHFHSETKNQSPNVAKVQSFCELQTHPQSHFQSETKKQSP
NVAKIQSFCELQTPPLSHFQSETKNQSTQLGNFFSFQSCHQNEPISFSGDHHHGFSFSPQNFPTVLGQNQMFNYQ
REPLQSTSNFPQTISYSNNIGFANDGLLGLSSAPITQQQQQQDEQGSISTTLLHYQD

**>Solyc10g018710**

MPQSLCKKPEKWANFTVSKQELNKGTRRMKQKRAKTDVIEGHGGRTIRATGRKYRHSKVSTAKGPKDRRVRLSPN
AAIQFYDVQDRLGYDRPSKAIDWLIKEAKAAIDALCEFPNNFHSSKLNPKKMQYSFDQEQCPEFSQENRGVPNSE
CGVQDKQQEVNYHIPILFSLY

**>Solyc11g020670**

MEFMEPHRKKLGDASNHHQHNQEETPSSLQLISPHQSQRDPSTPGSNHPHGPFMGSISMQSISPPSTNSSTPNNN
TISTLKVAKKPSKDRHTKVDGRGRRIRMPALCAARVFQLTRELGHKSDGETIEWLLQQAEPAIIATTGTGTIPAN
FSTLNVSTRSSGTTISAPPSKSAPLFIHGASASASAAMLGFHHHLSTANTGFIQDPDENYMKKRFREDTTTSGAT
SPSPDKPGSDSSKPGSNLIPGQAMWAVAPAGGNVGNGFWMLPVSGGTGSTAVSVGAGQSDHQLWQYKSSRIGGLE
FPGGGRFSPVQLGSMVLQQSHPVQQLGSNMGMLNAYNNNNNNSRVDLGMNLEHHNQTQHQDSDSGDENHNDSQS

**>Solyc11g045640**

MSNLNPPMNHQCQVSDEEDGSEEEEEEEEEVFQENDIGNLQSHYQNQQMPQSVYKKQEKWANFTVSEQELNKGTR
RMKPKRAKTDVIEGHGGRIIRATGRKDRHSKVSTAKDPKDLRVRLSPNTAIQFYDVQDRLGYDRPSKAIDWLIIE
AKAAIDALGEFPNNFHSTKLNPKKMQYSFDQEQSPEFSQENRGVPKSEYGVQEKQQEVNYDIPNLFSLN

**>Solyc12g014140**

MYEDLDTIKQQQEQARSKKRSFFPPPPPRPSCSKQKGVTSSKFGCGEIVEVQGGHIIRSIGRKDRHSKVCTAKGP
RDRRVRLAAHTAIQFYDVQDRLGYDRPSKAVDWLIKKAKASIDELAELPPWKPTTTGVDHALQDDIDVGTASTIM
LGQNNSASSSGENVNANTKADGSFMPHSLDSDAISDTIKSFFPMGGSGSNEGNSFQSFQQHNLMSRSQDLKLSLQ
SFQDPQAQFEASNFFTGFDASVWPQHQQQPVELGRLAAARGDIGGGAIAGAGSYLFNSQPAPPLLQQLFTQNQFL
SQRGPLQSSYSPSIRAWIDPSAIAIATADPIHQNQNHHQAVFPMYSTSLSGIGFASELGGFSGFRIPTRIQGEIE
EEHDGVSDKPSSASSDSRH
