## Supplementary File 3 for "Genome-Wide Analysis of *TCP* Family Genes and Their Constitutive Expression Pattern Analysis in the Melon (*Cucumis melo*)"

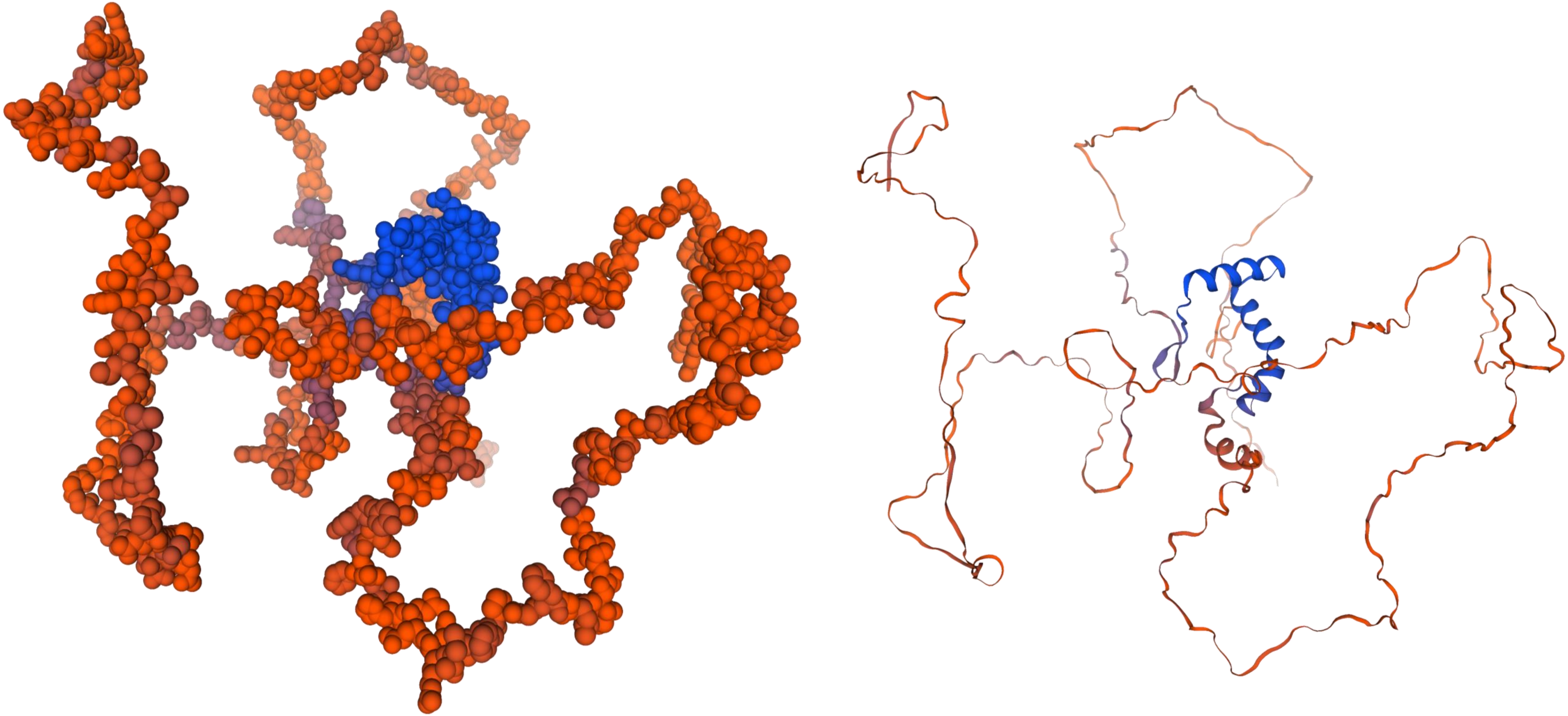

***CmTCP-01***

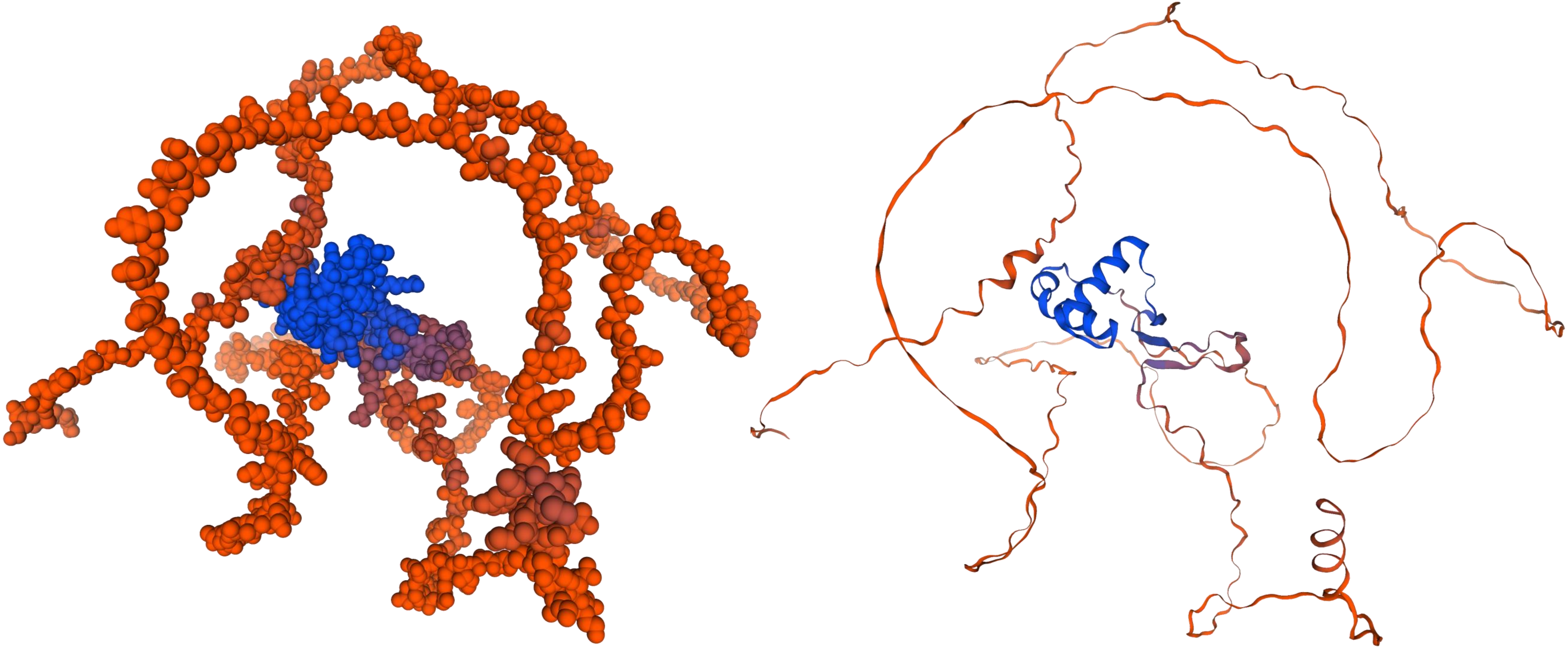

*CmTCP-02*

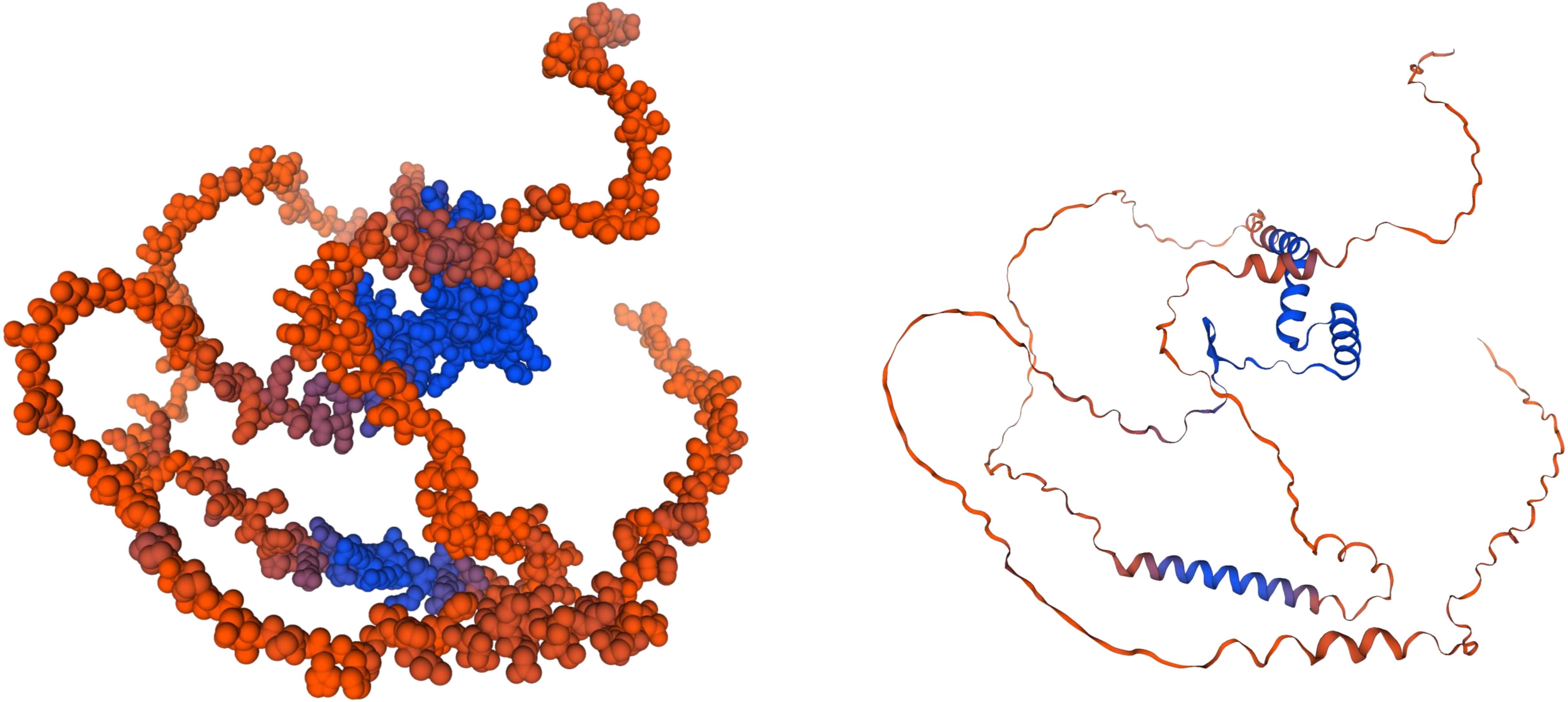

***CmTCP-03***

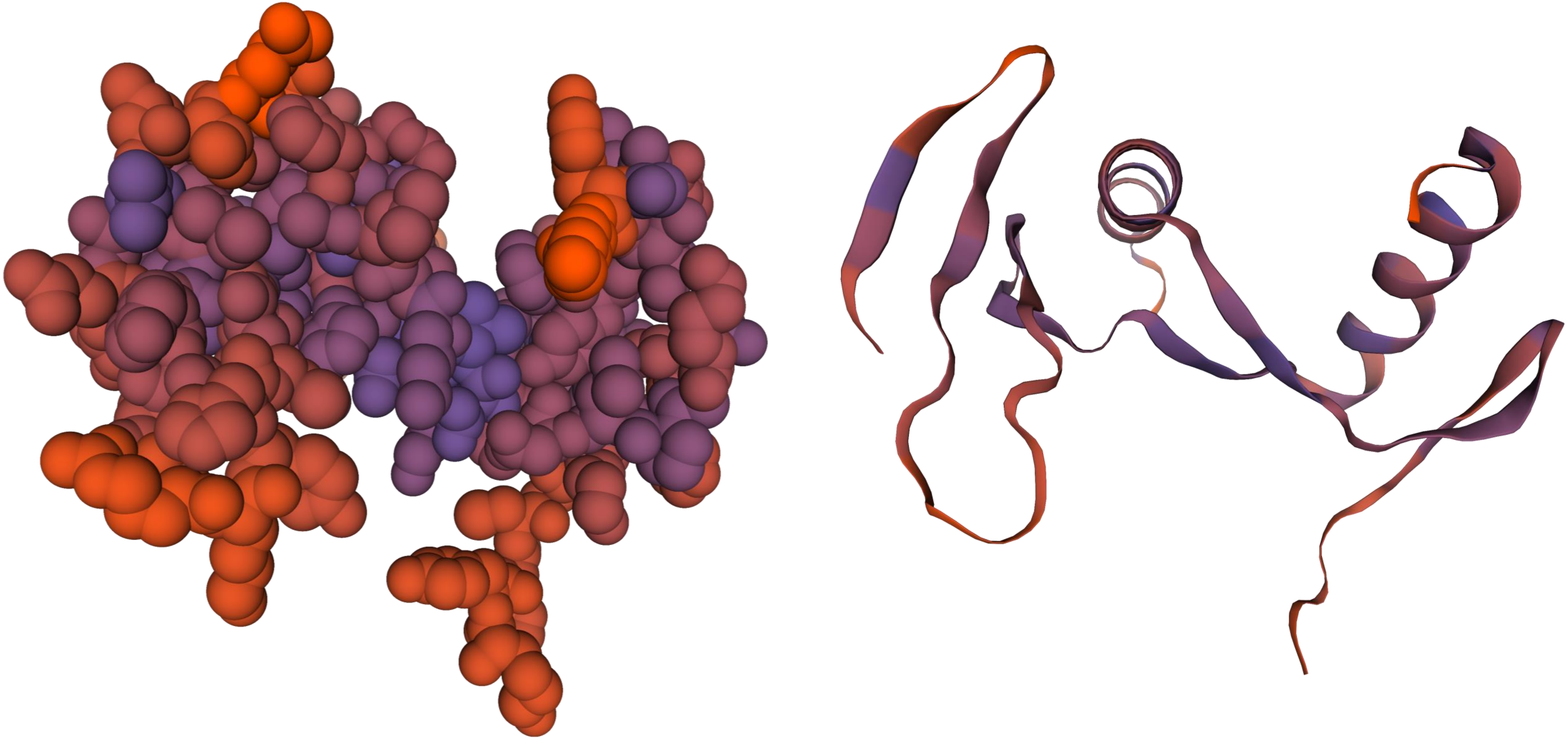

***CmTCP-04***

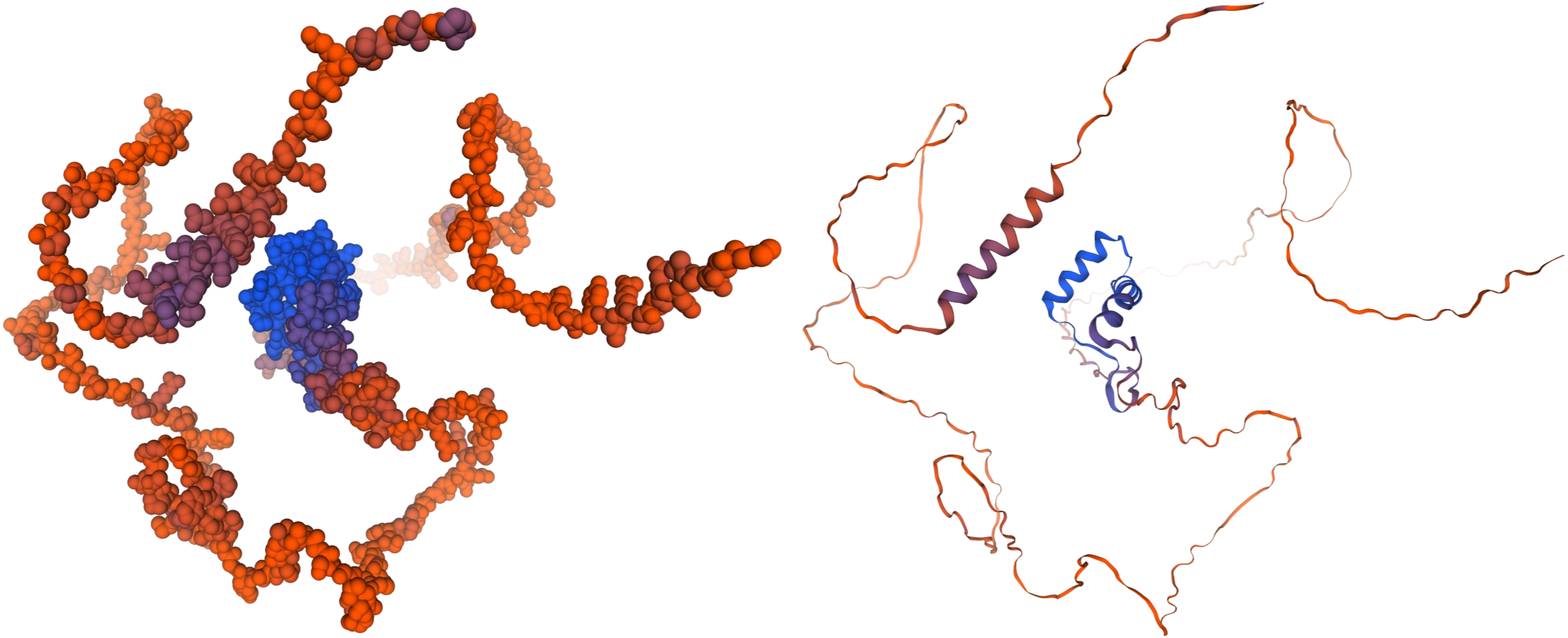

***CmTCP-05***

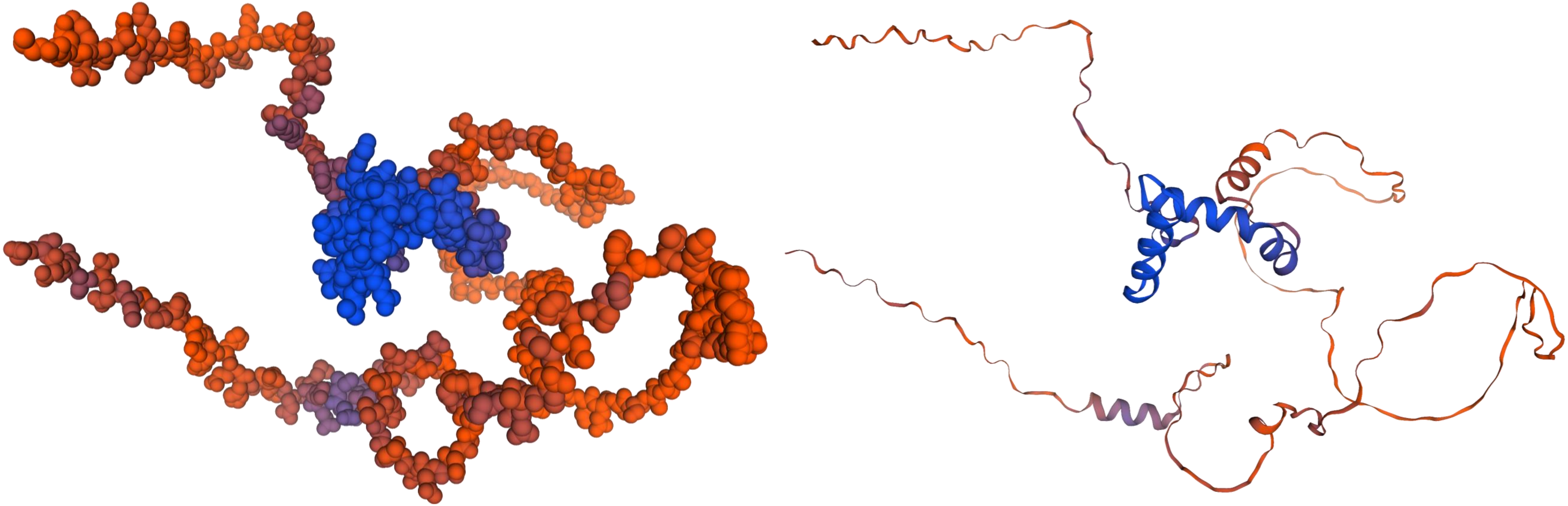

*CmTCP-06*

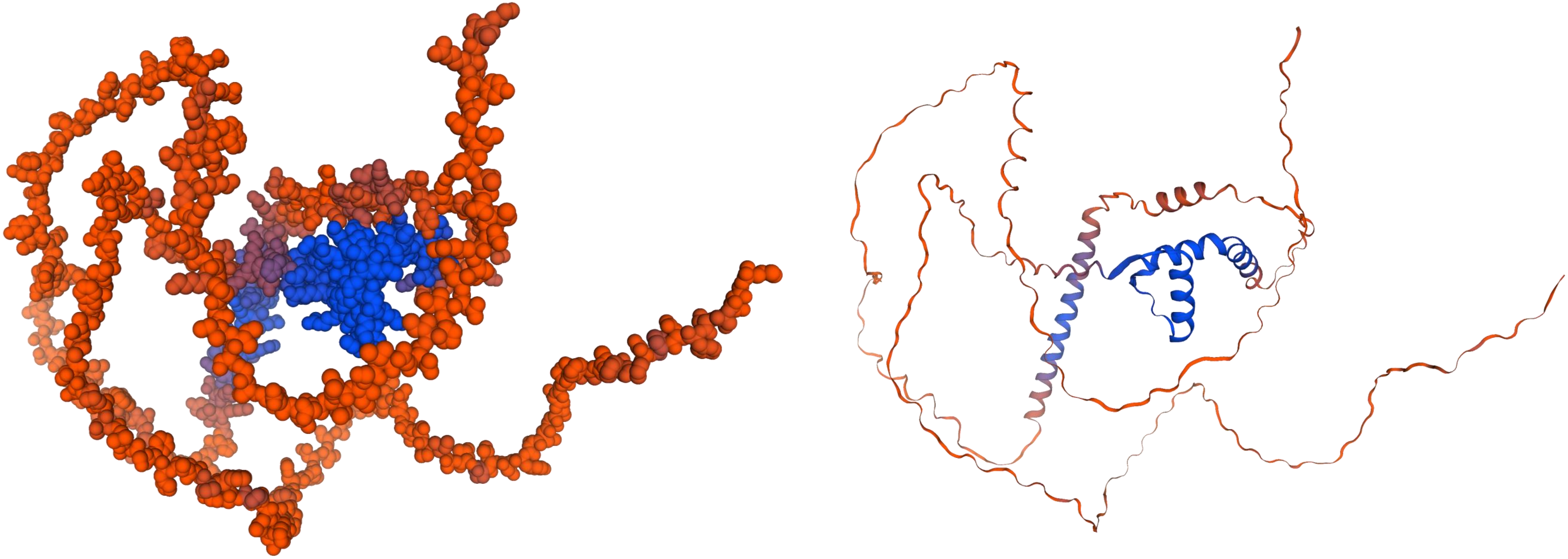

***CmTCP-07***

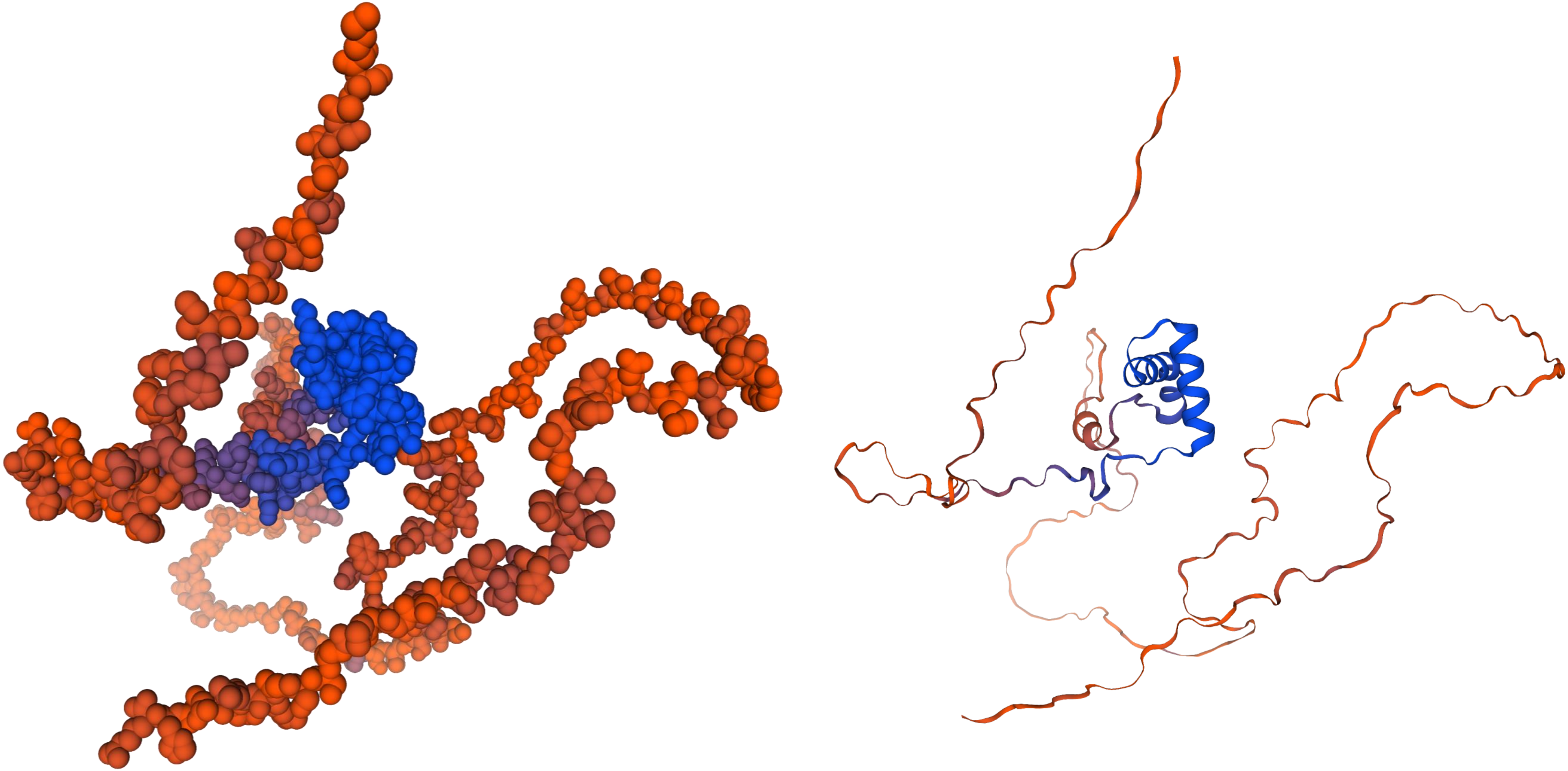

*CmTCP-08*

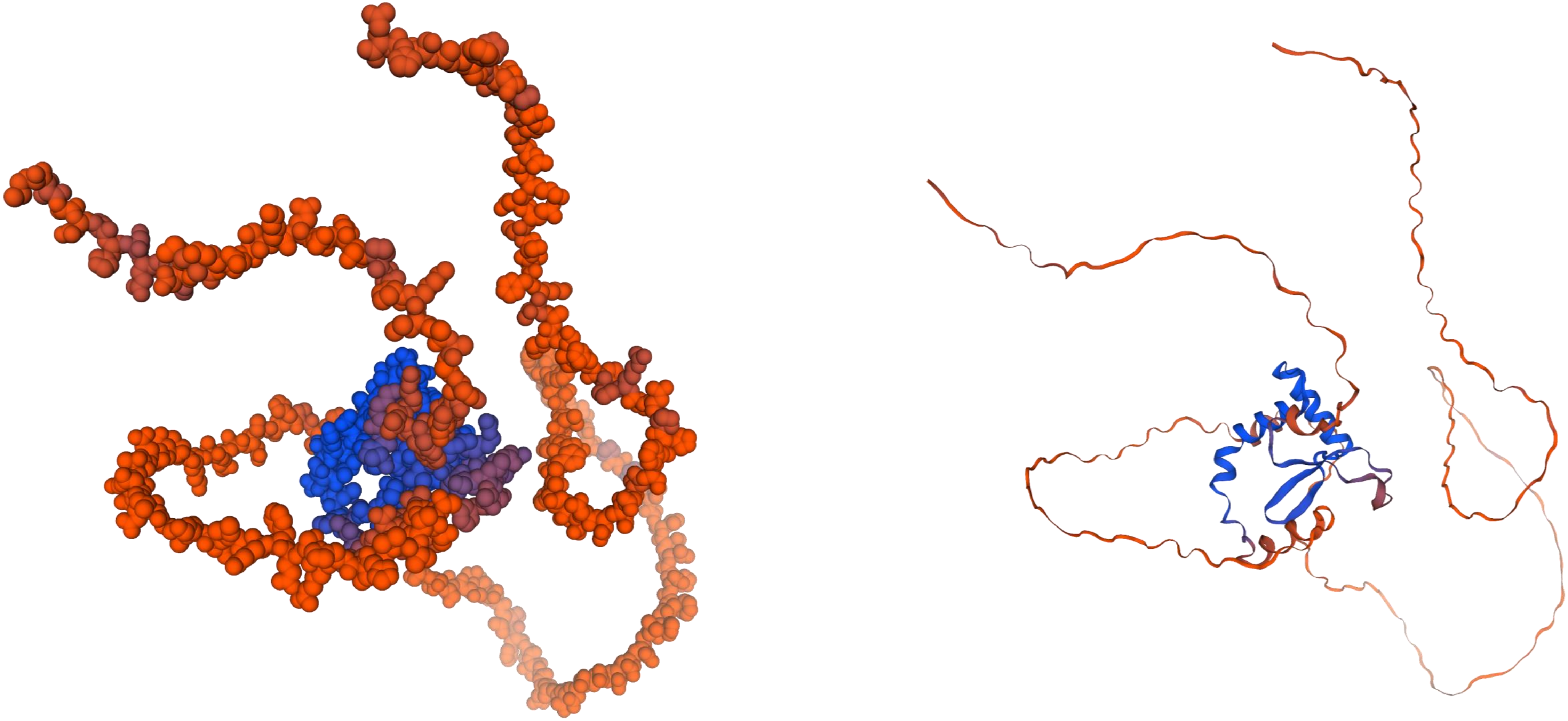

***CmTCP-09***

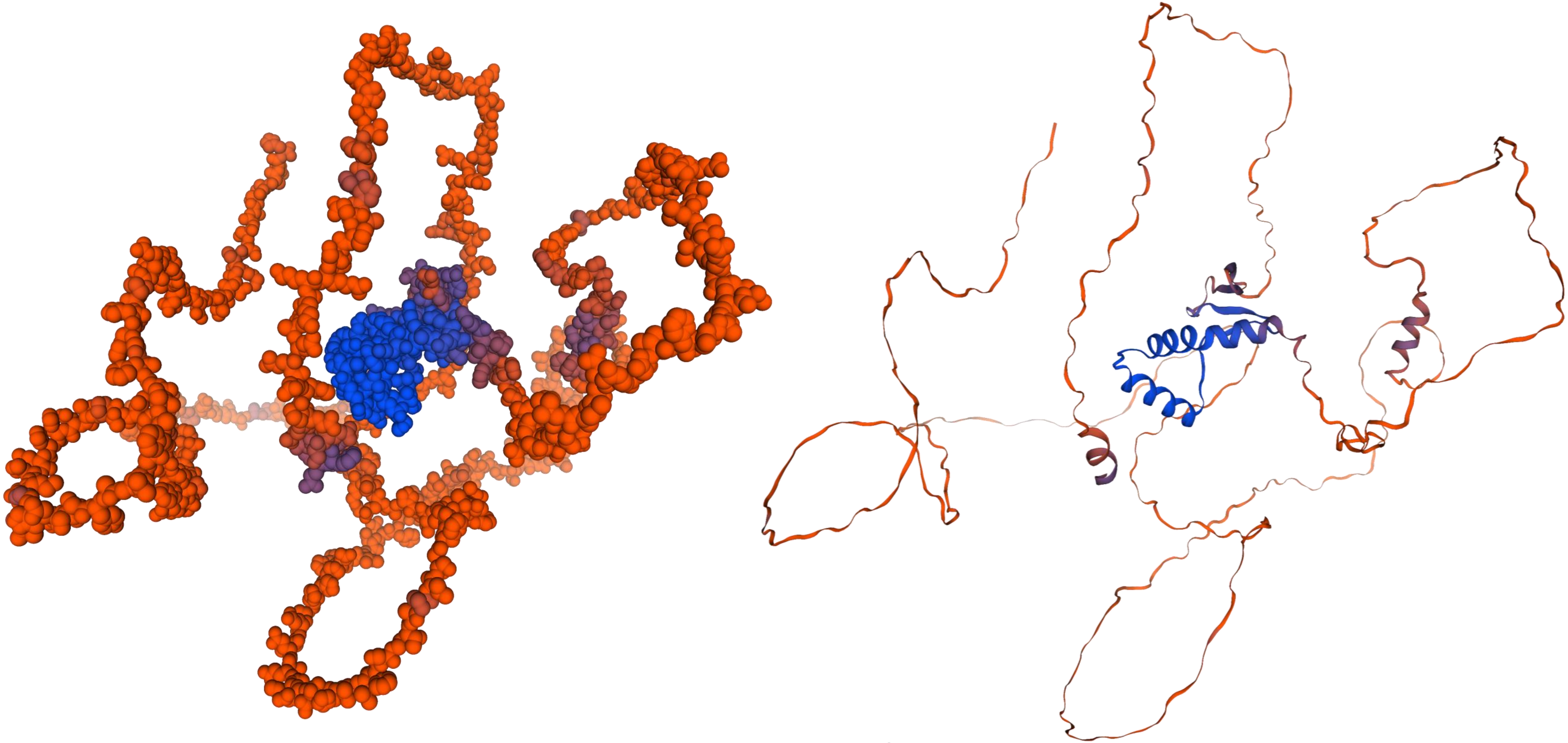

***CmTCP-10***

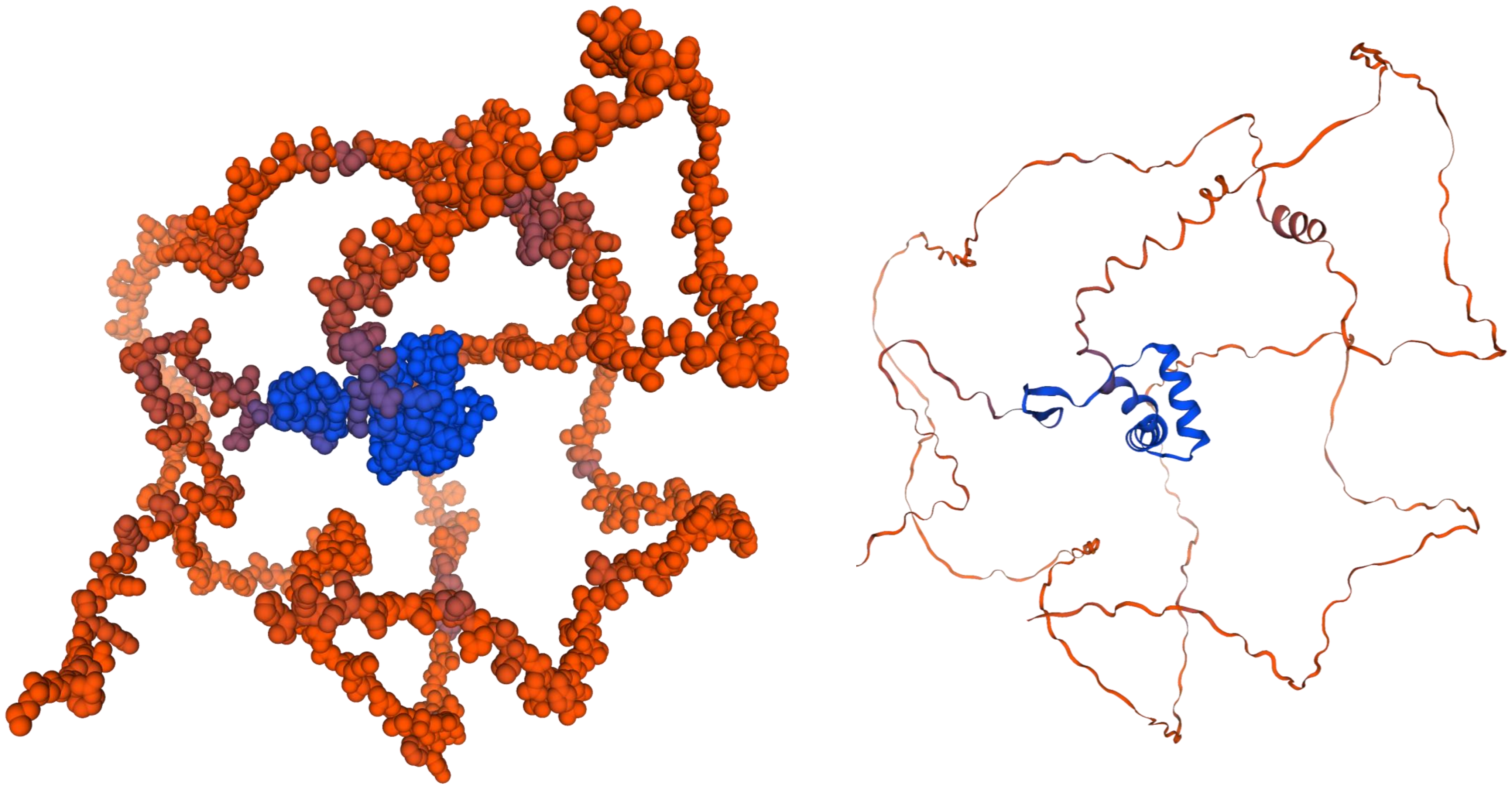

***CmTCP-11***

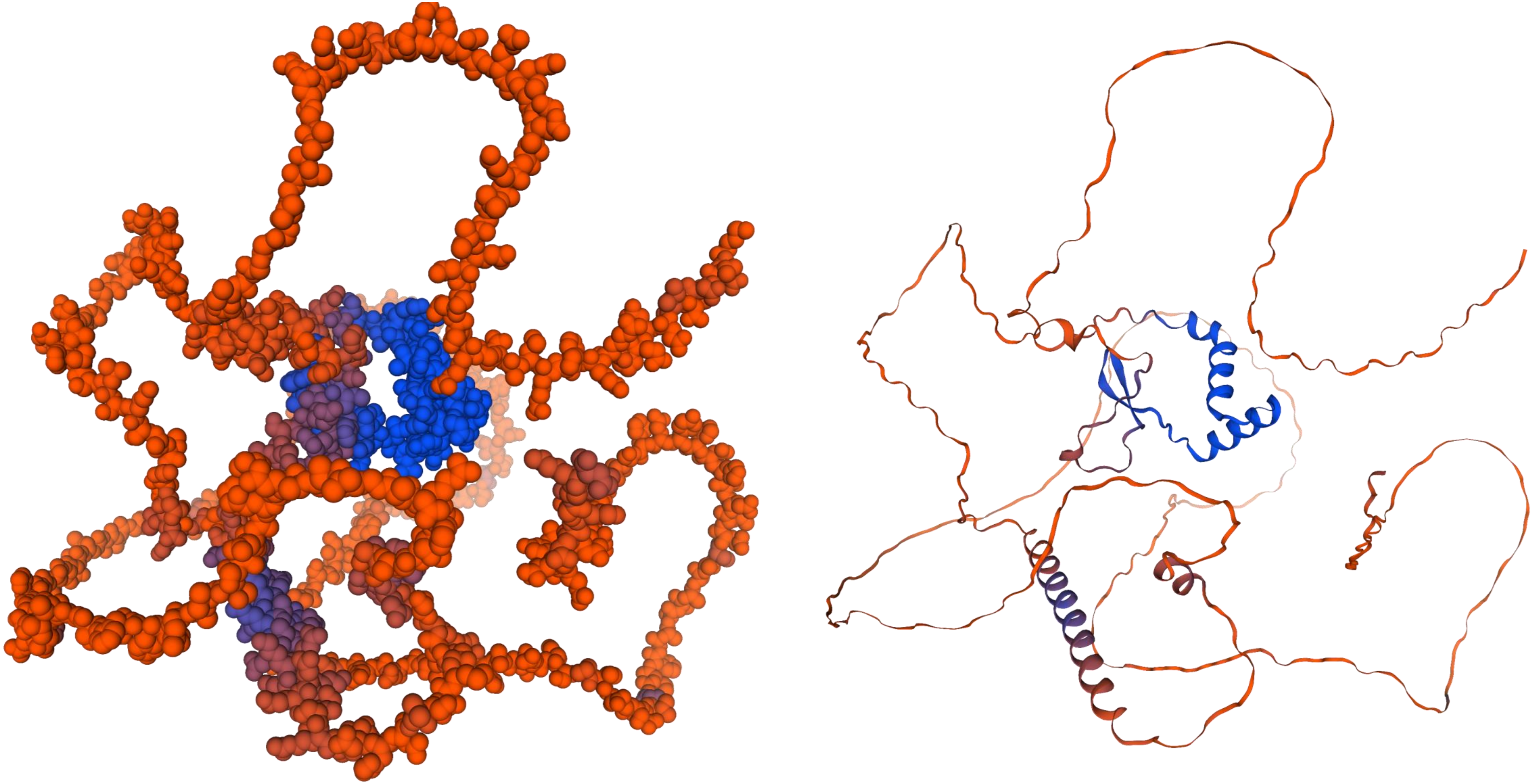

*CmTCP-12*

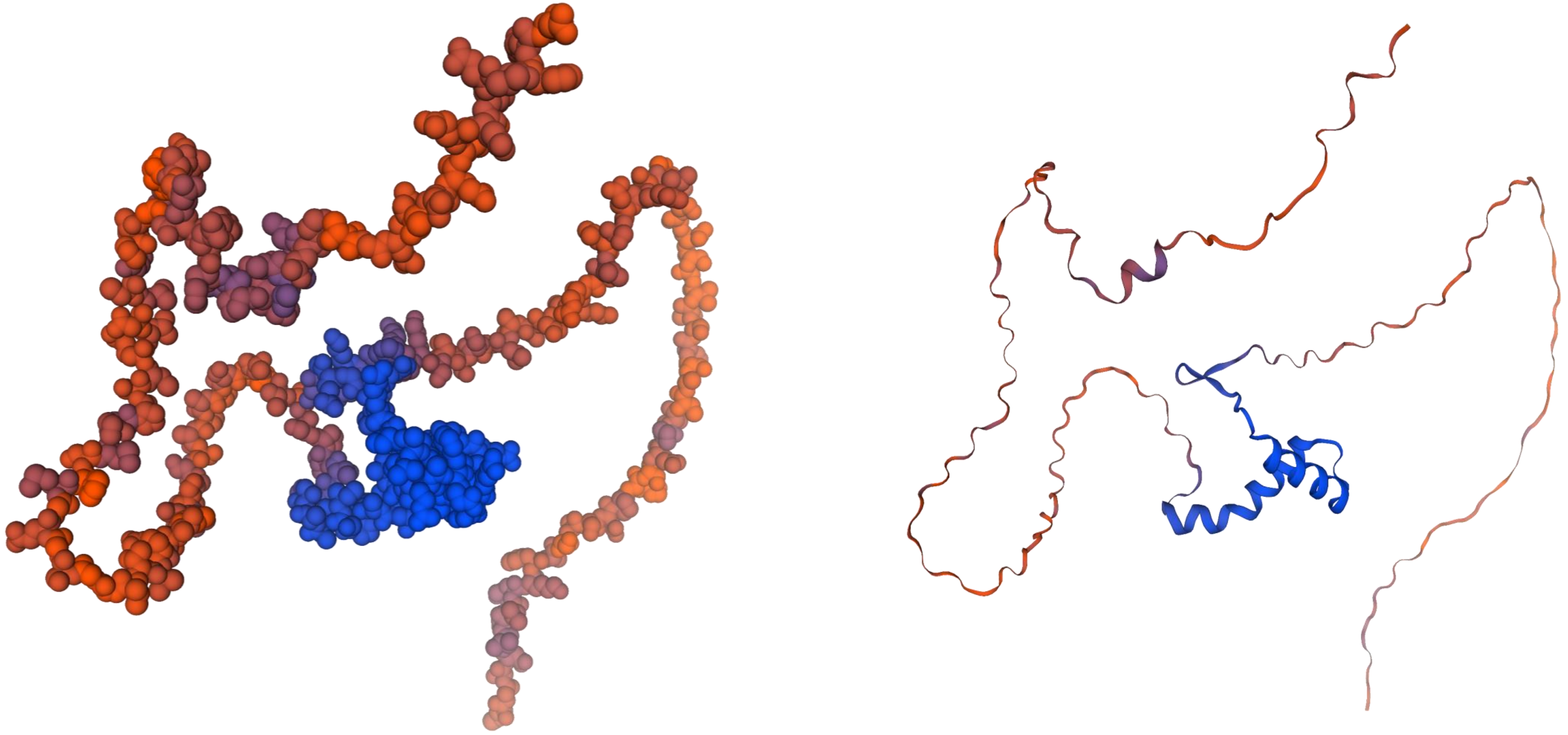

*CmTCP-13*

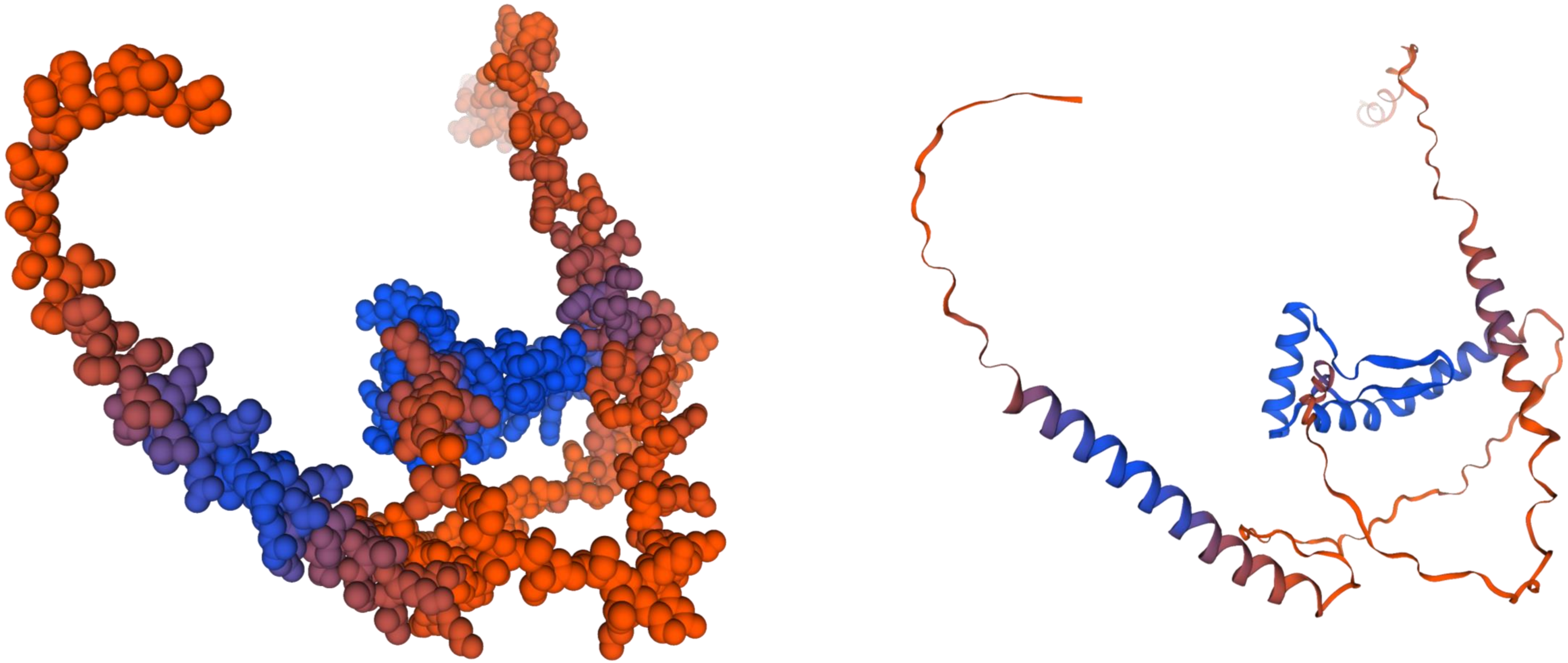

***CmTCP-14***

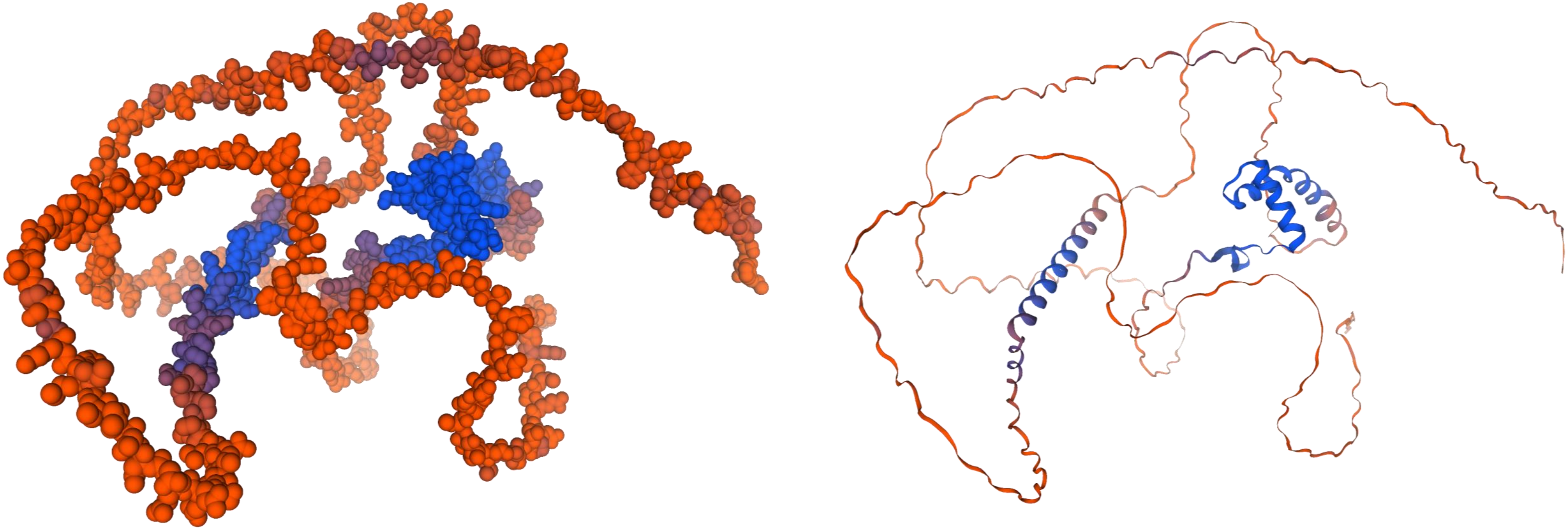

*CmTCP-15*

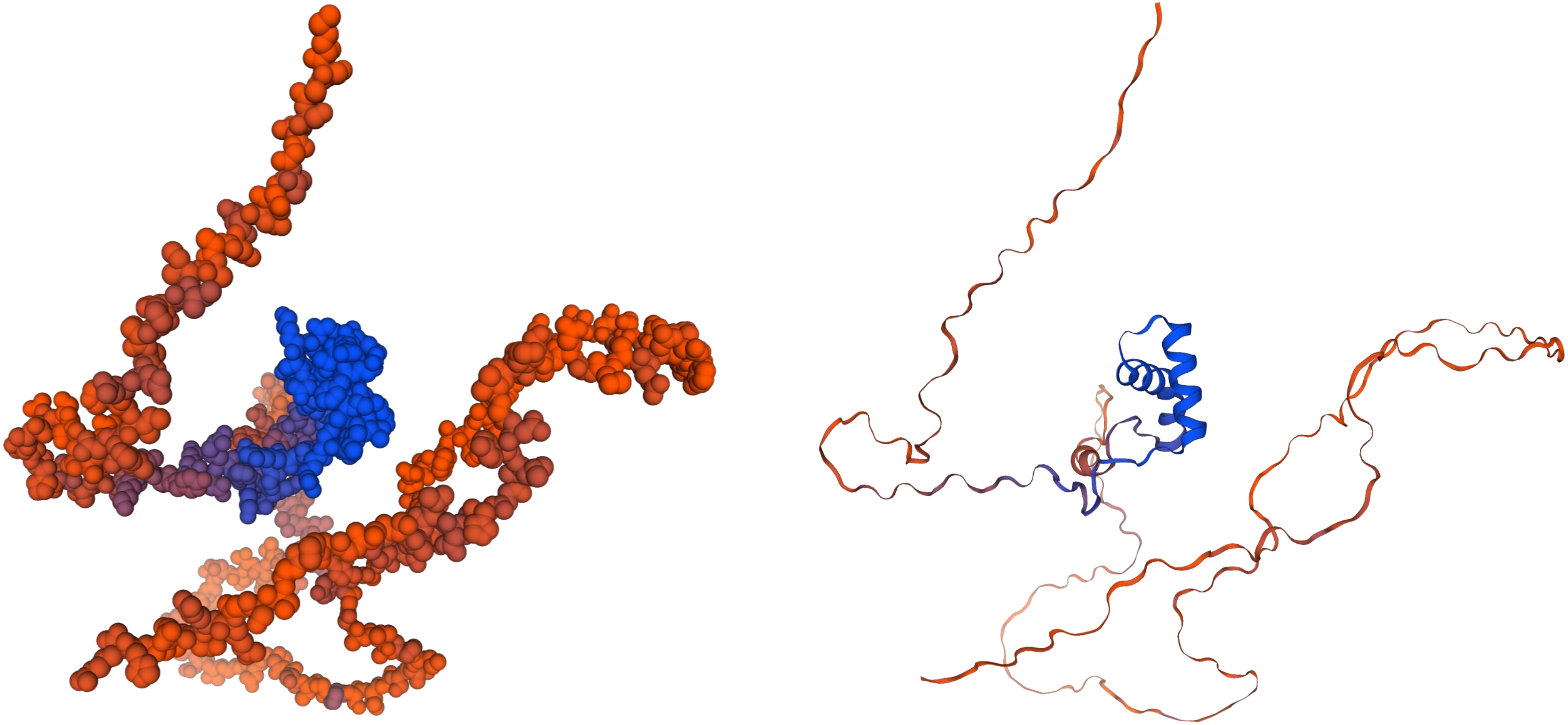

*CmTCP-16*

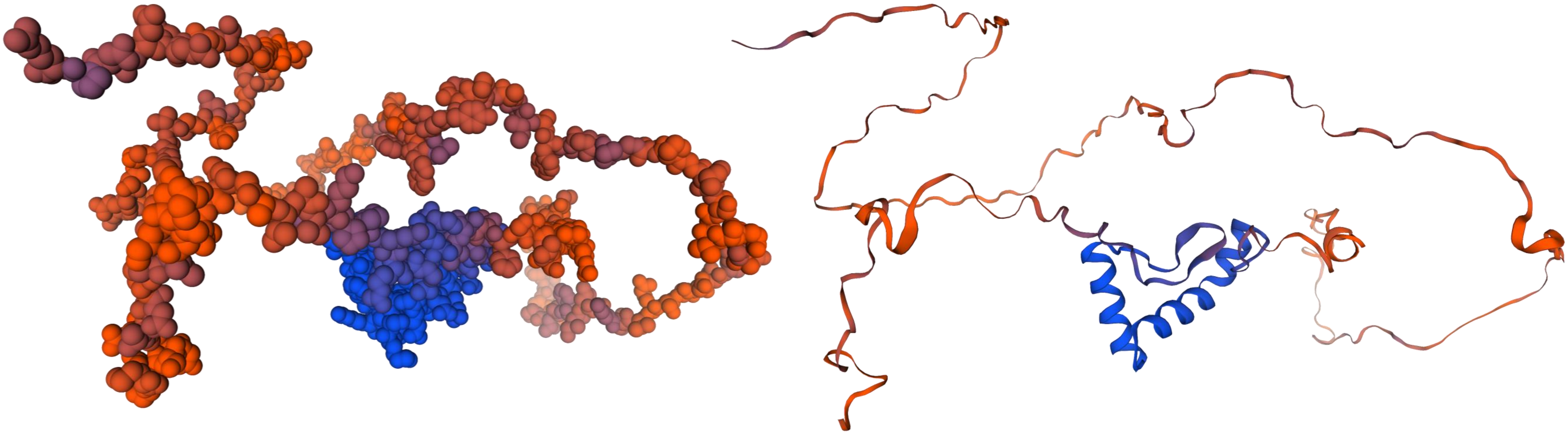

*CmTCP-17*

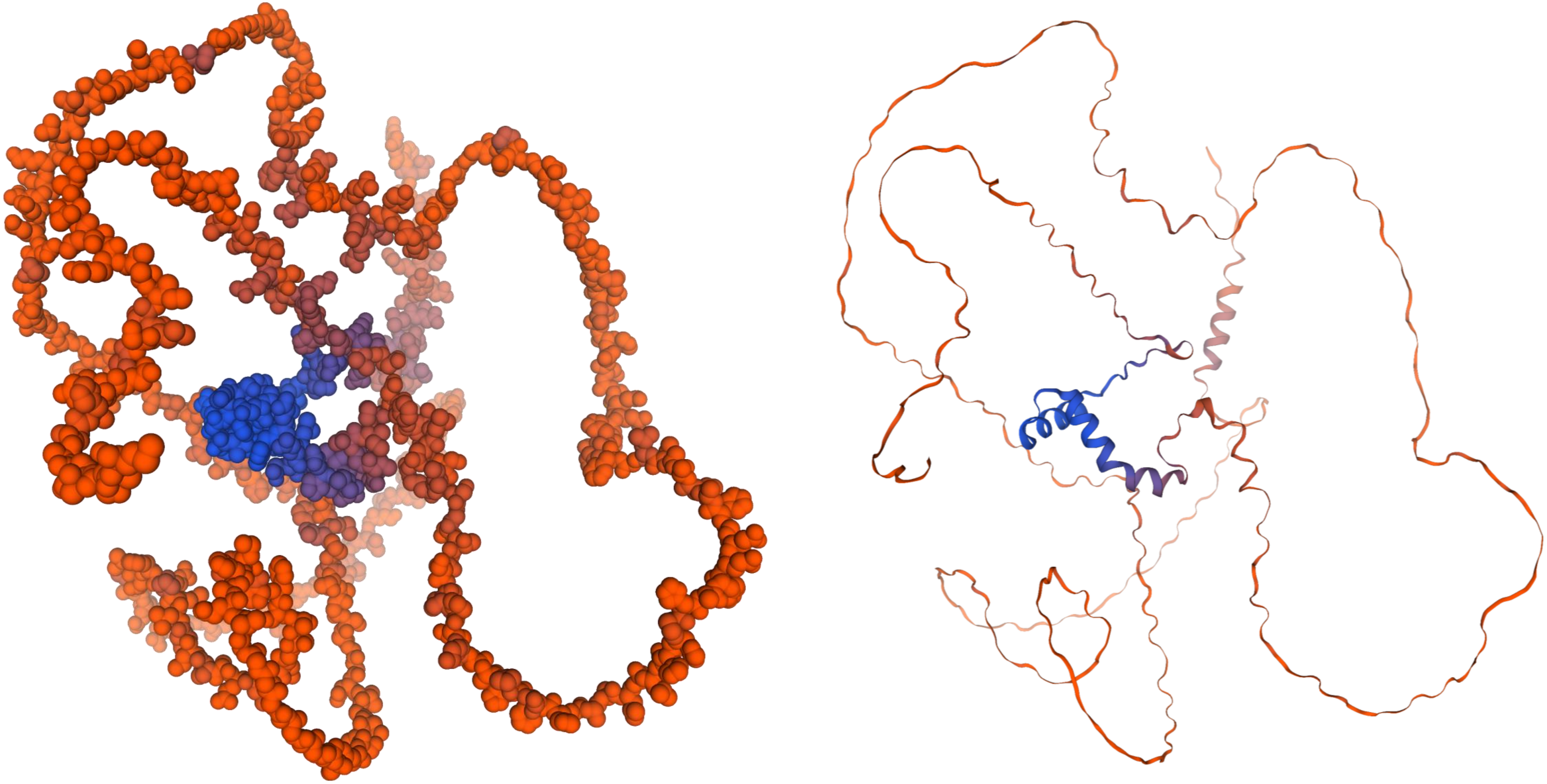

*CmTCP-18*

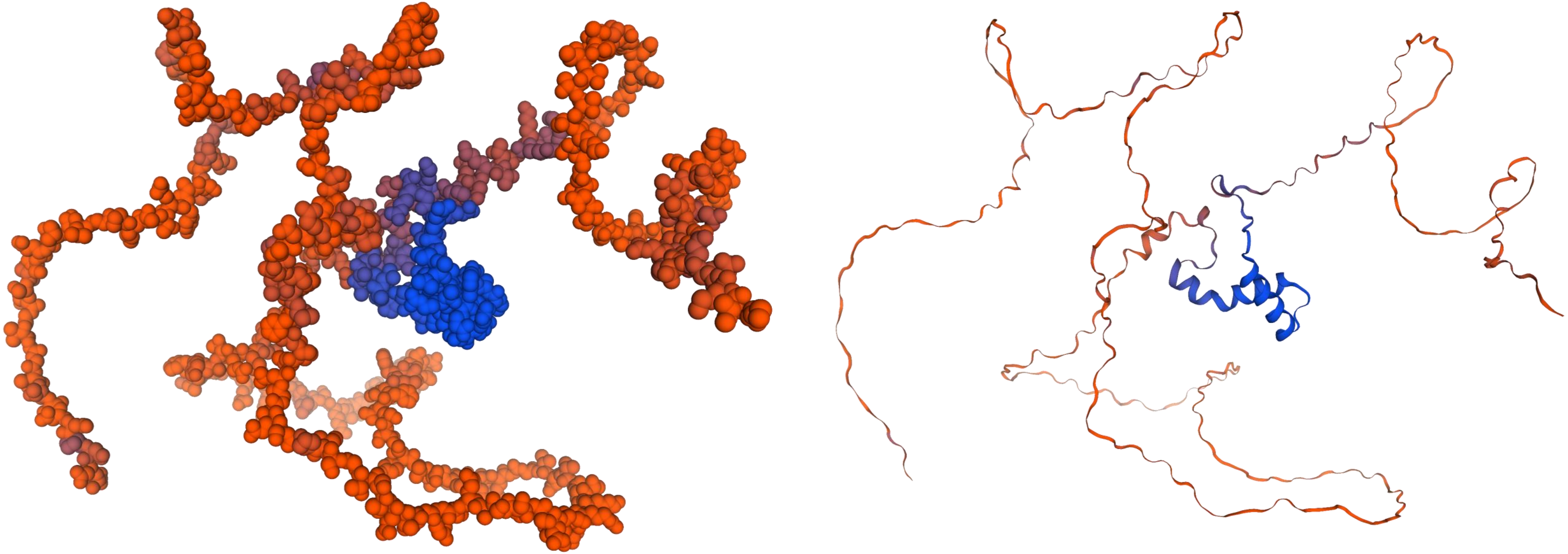

***CmTCP-19***

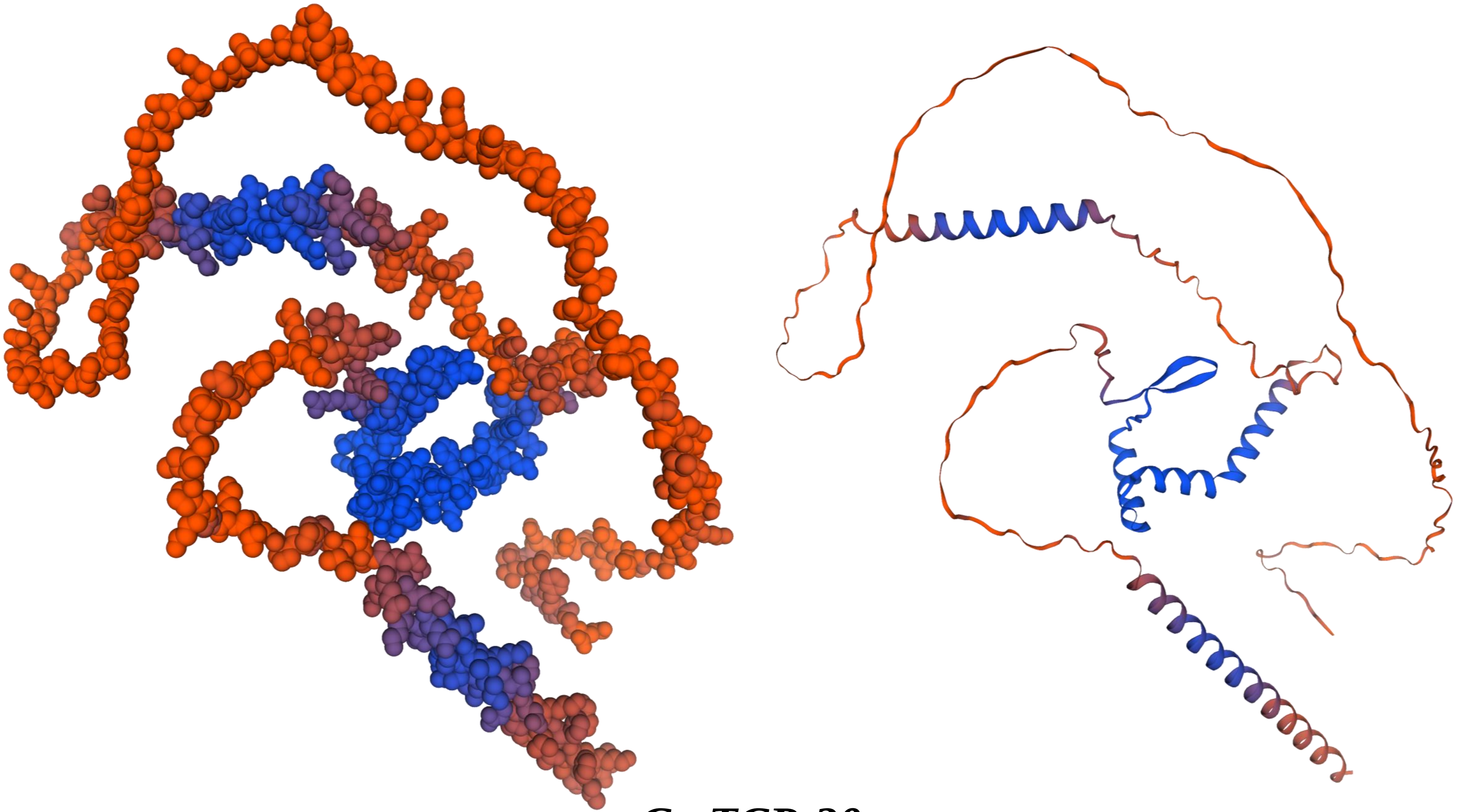

*CmTCP-20*

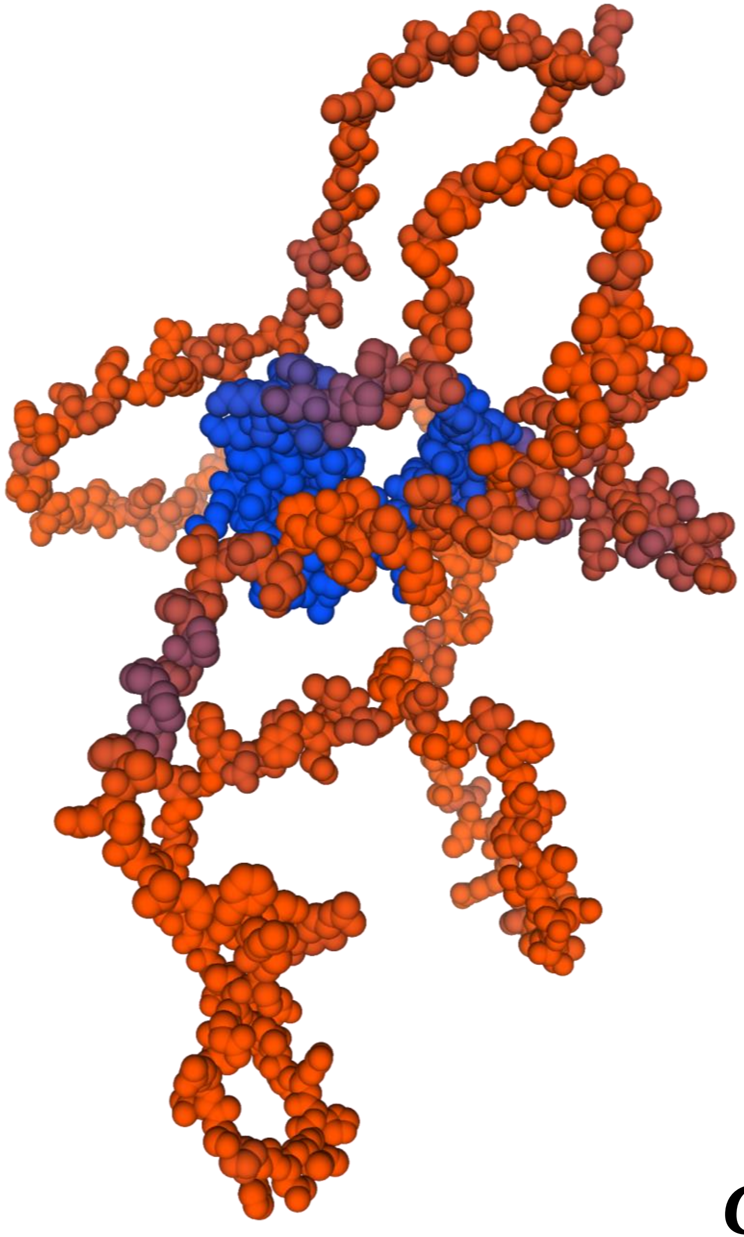

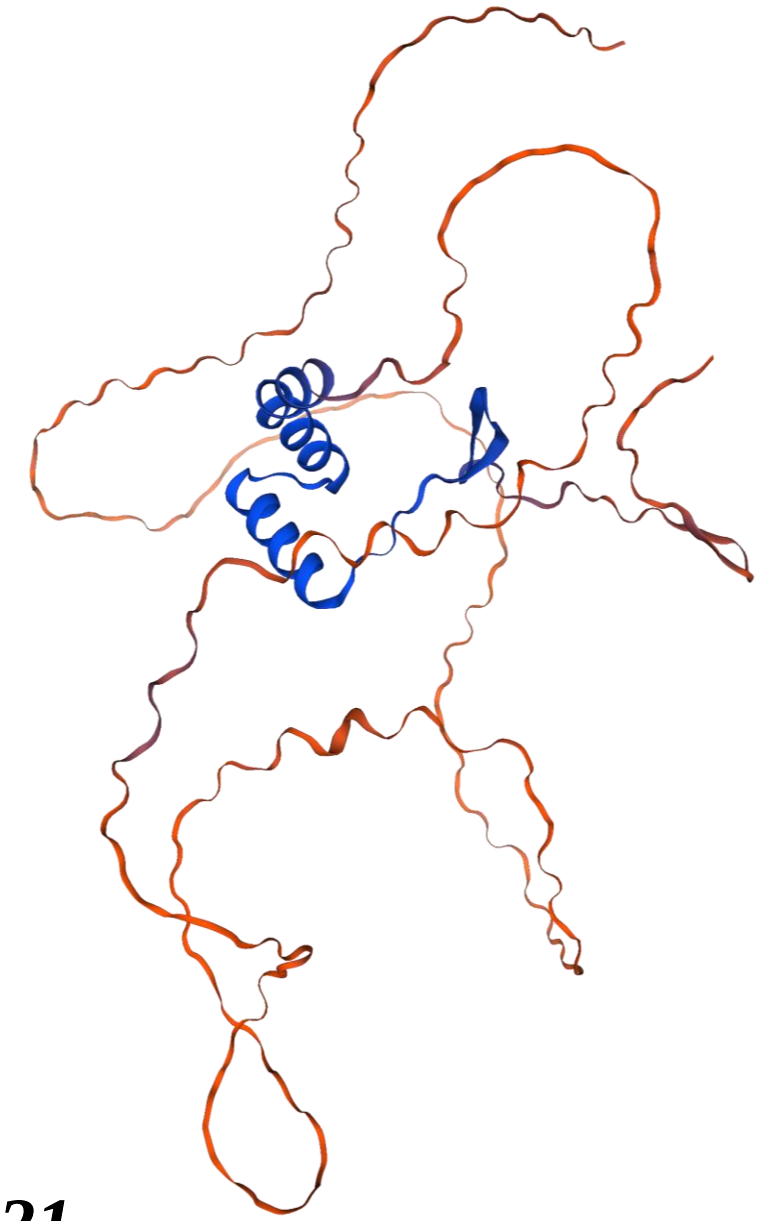

***CmTCP-21***

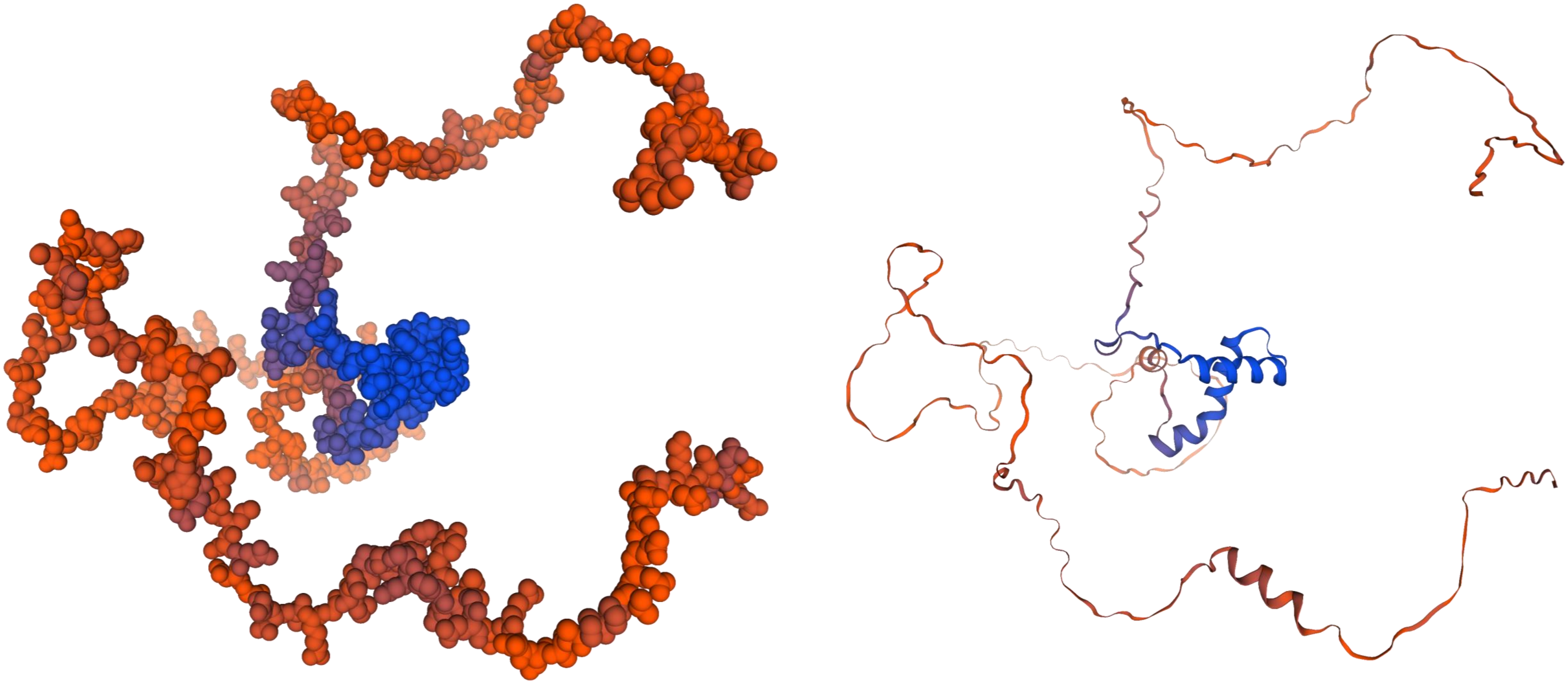

***CmTCP-22***

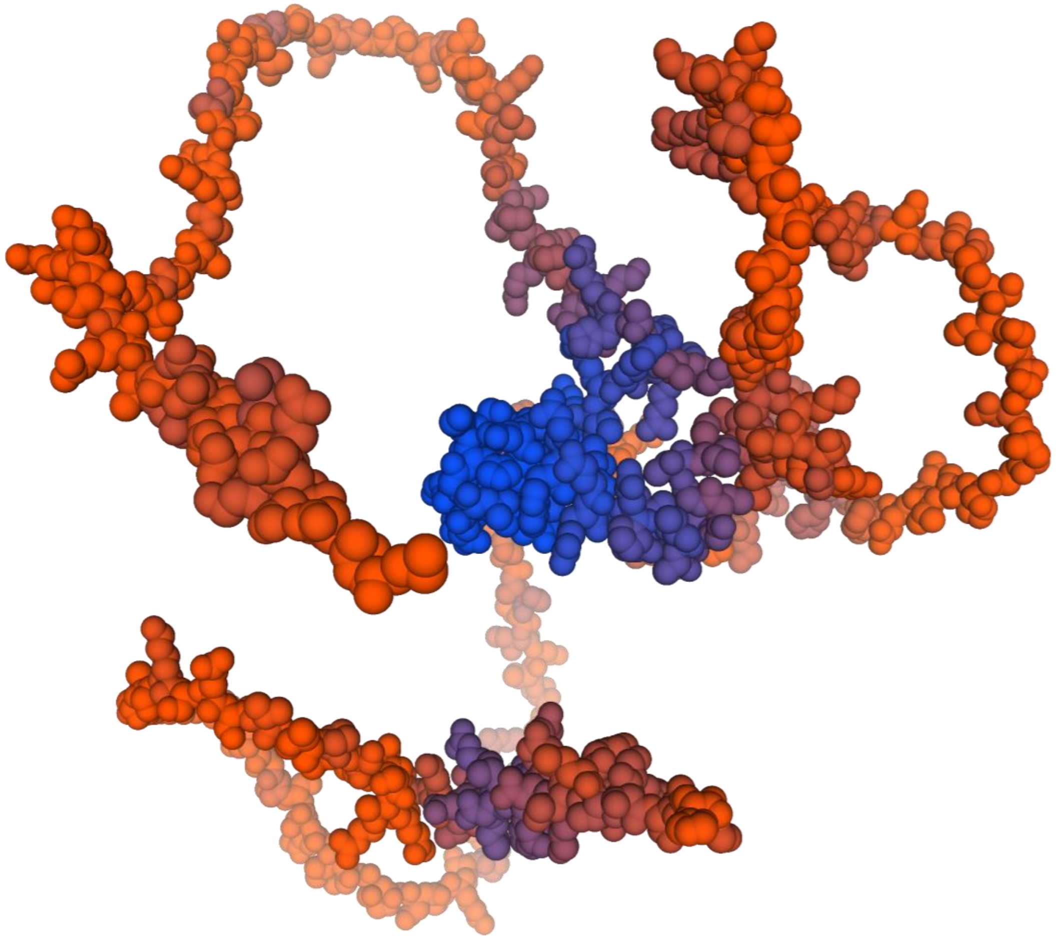

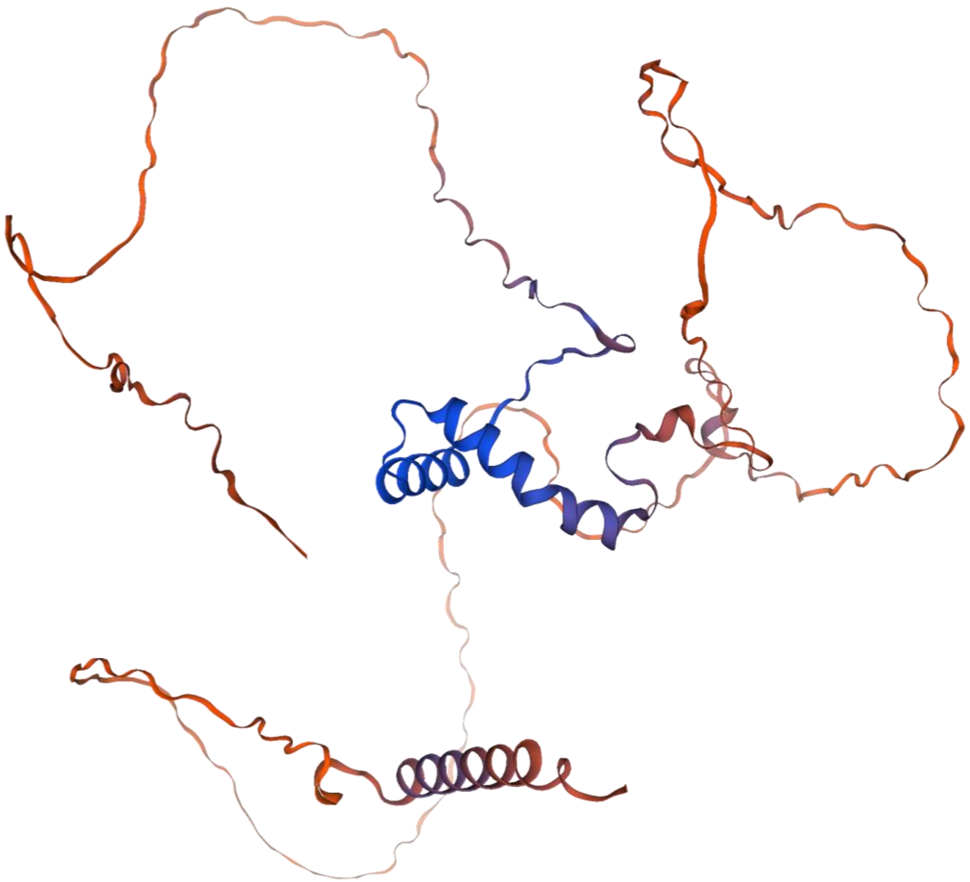

***CmTCP-23***

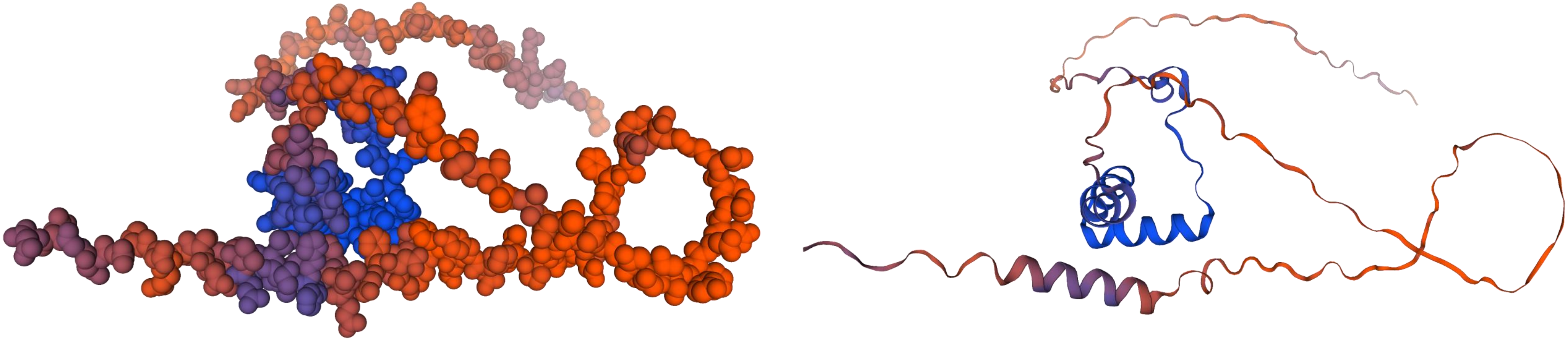

***CmTCP-24***

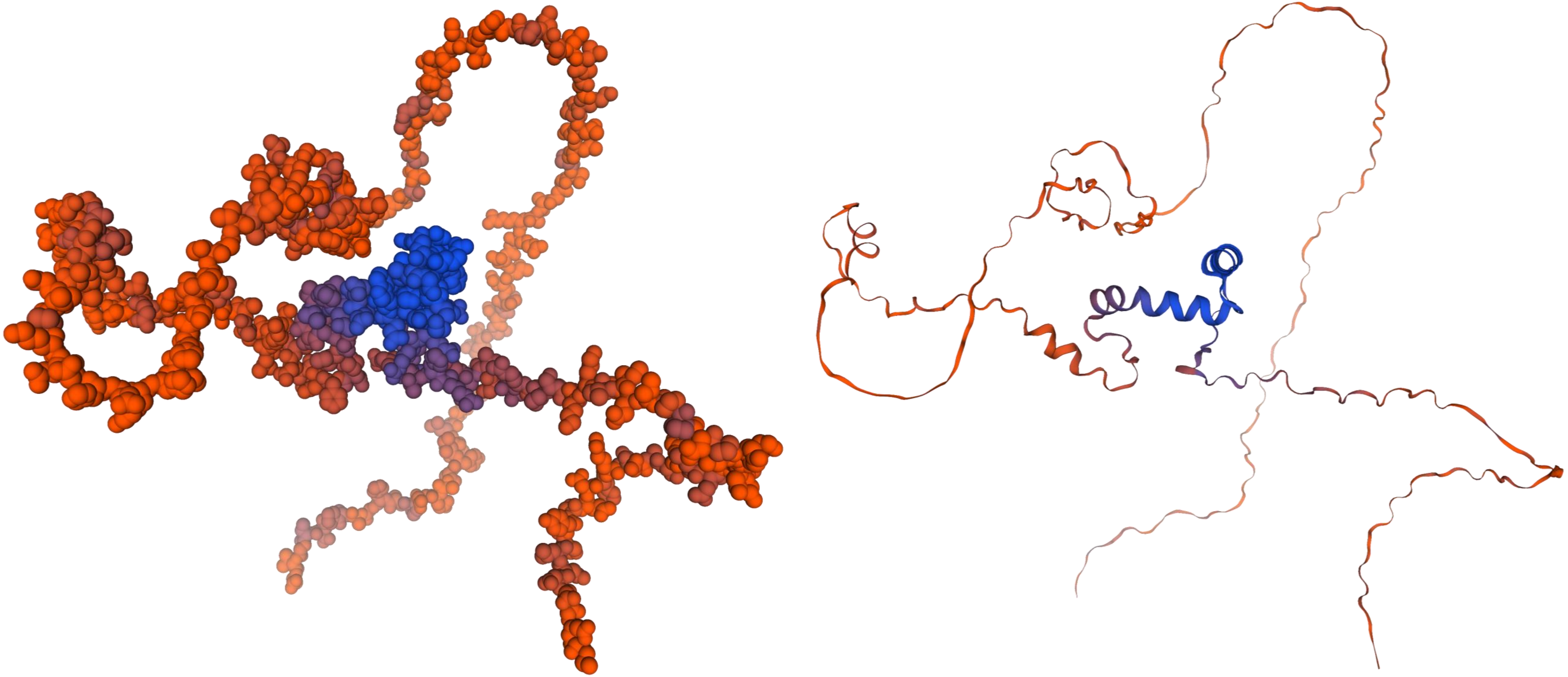

***CmTCP-25***

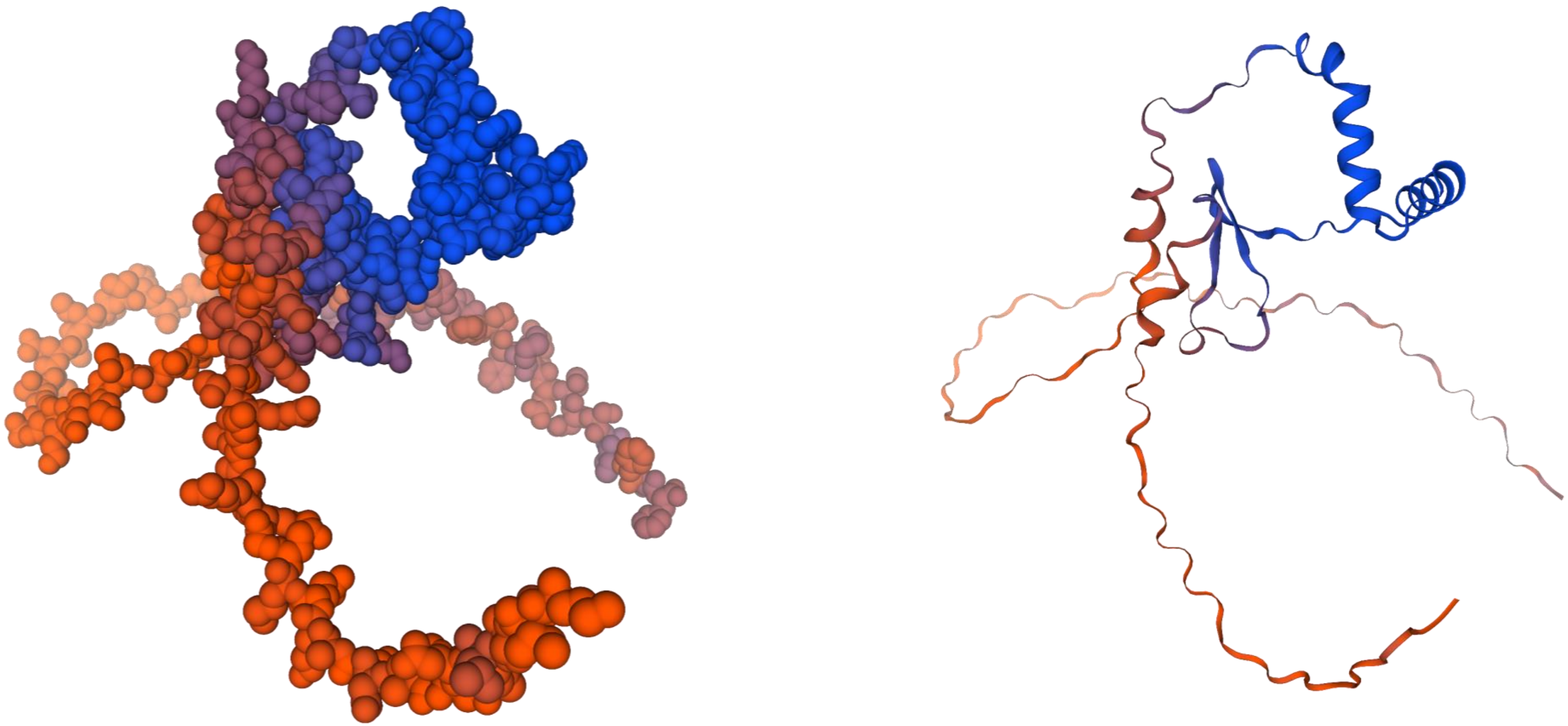

***CmTCP-26***

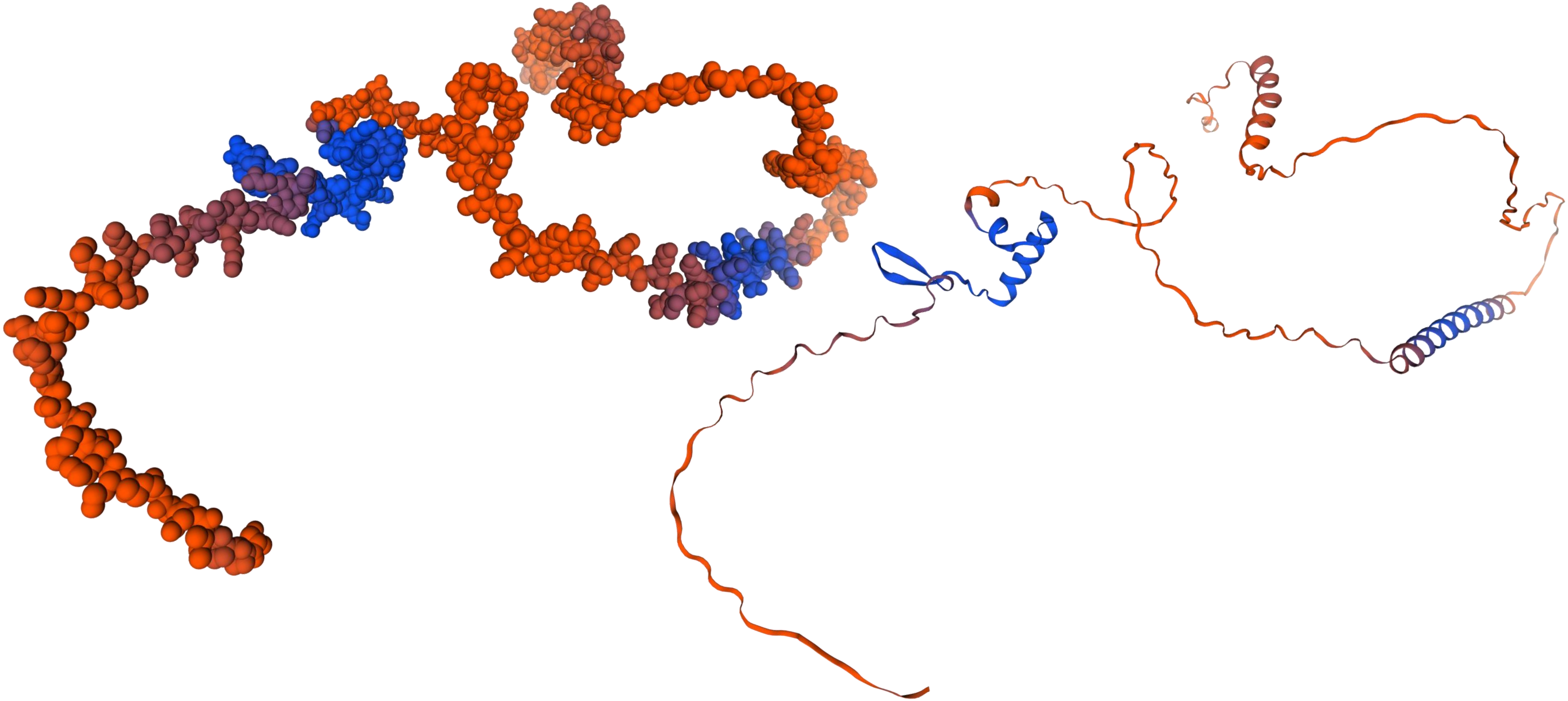

*CmTCP-27*

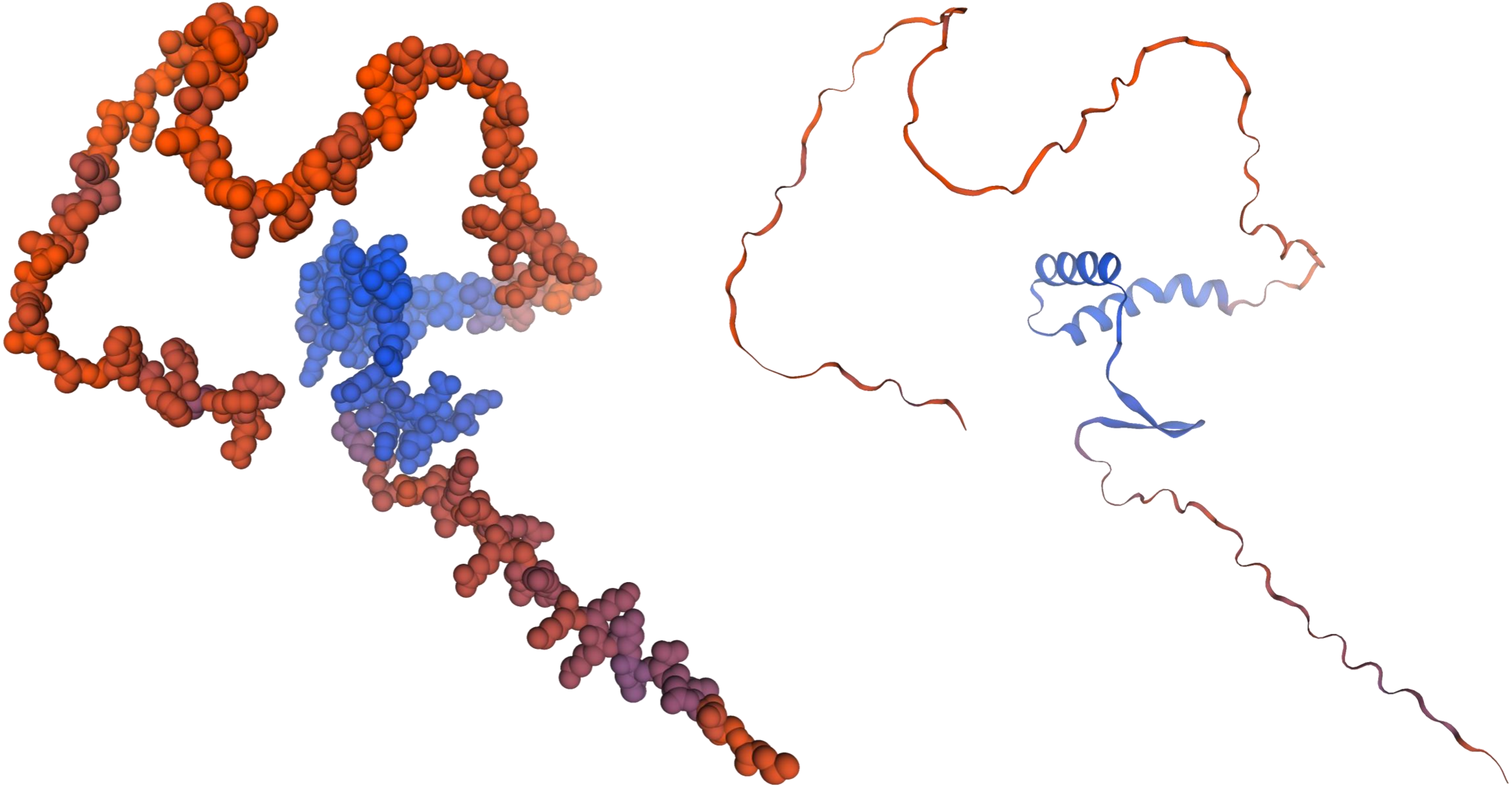

*CmTCP-28*

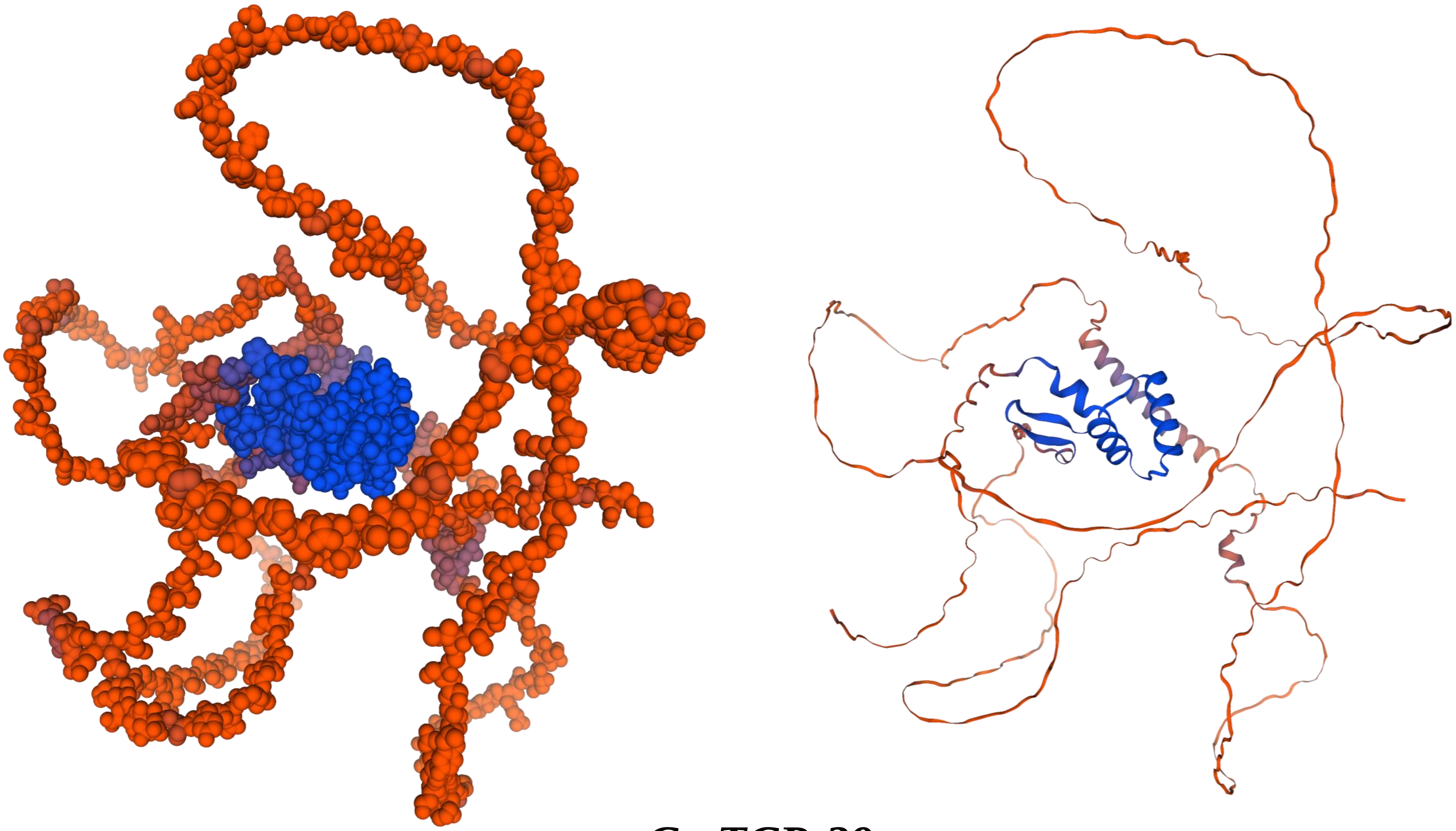

***CmTCP-29***

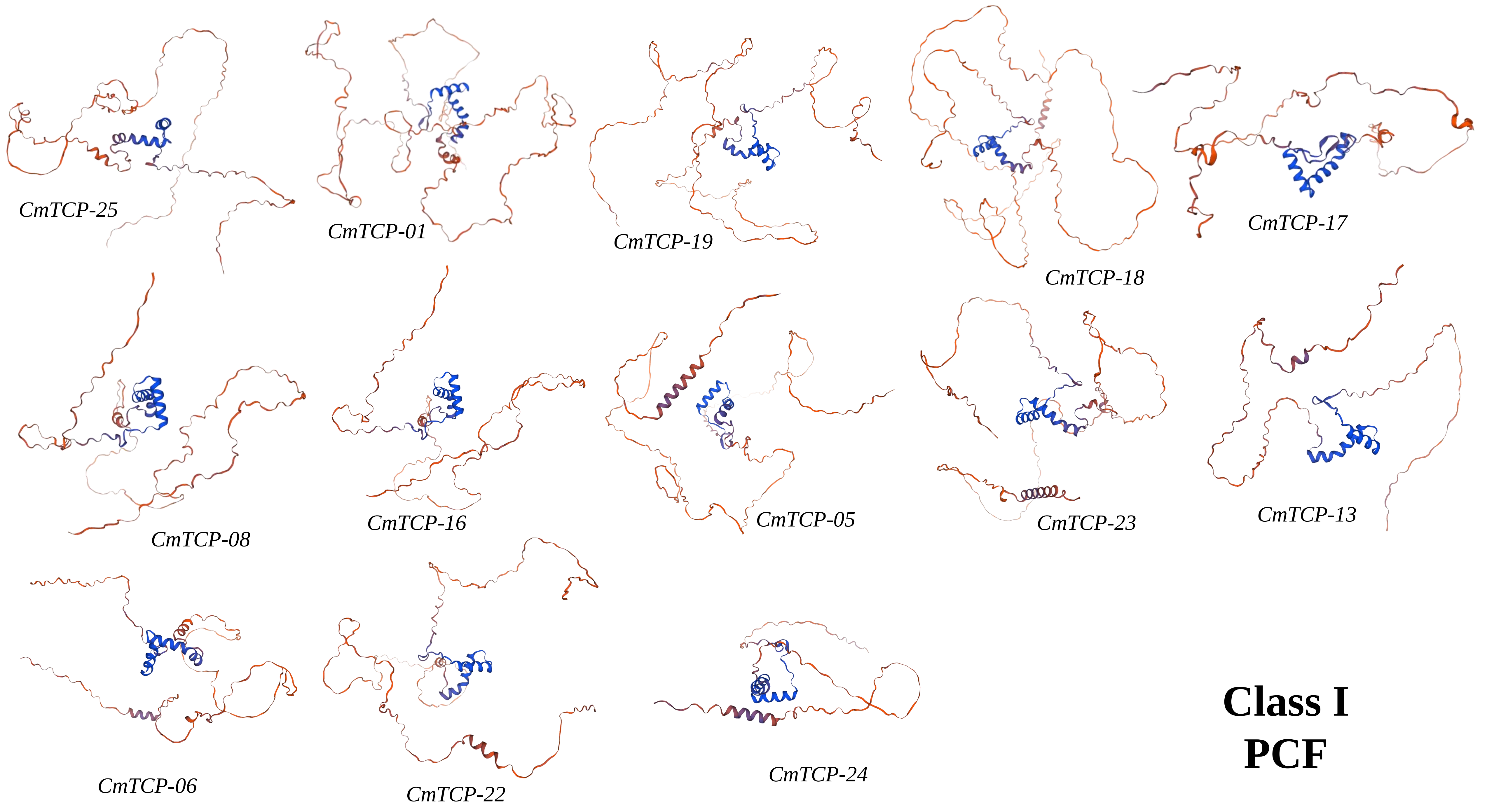
CmTCP-25
CmTCP-01
CmTCP-19
CmTCP-18
CmTCP-17
CmTCP-08
CmTCP-16
CmTCP-05
CmTCP-23
CmTCP-13
CmTCP-06
CmTCP-22
CmTCP-24
Class I
PCF

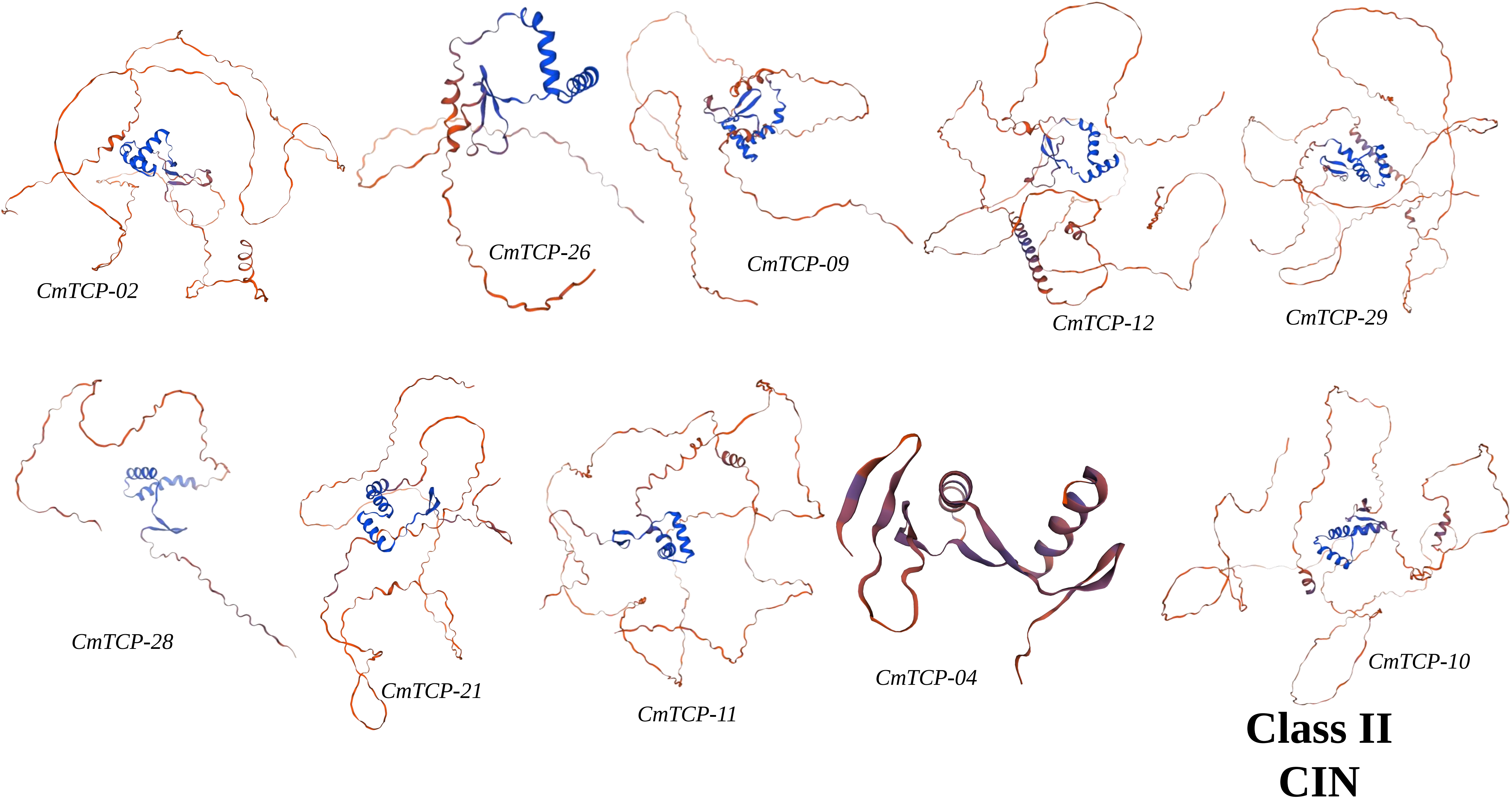
CmTCP-02
CmTCP-26
CmTCP-09
CmTCP-12
CmTCP-29
CmTCP-28
CmTCP-21
CmTCP-11
CmTCP-04
CmTCP-10
Class II
CIN

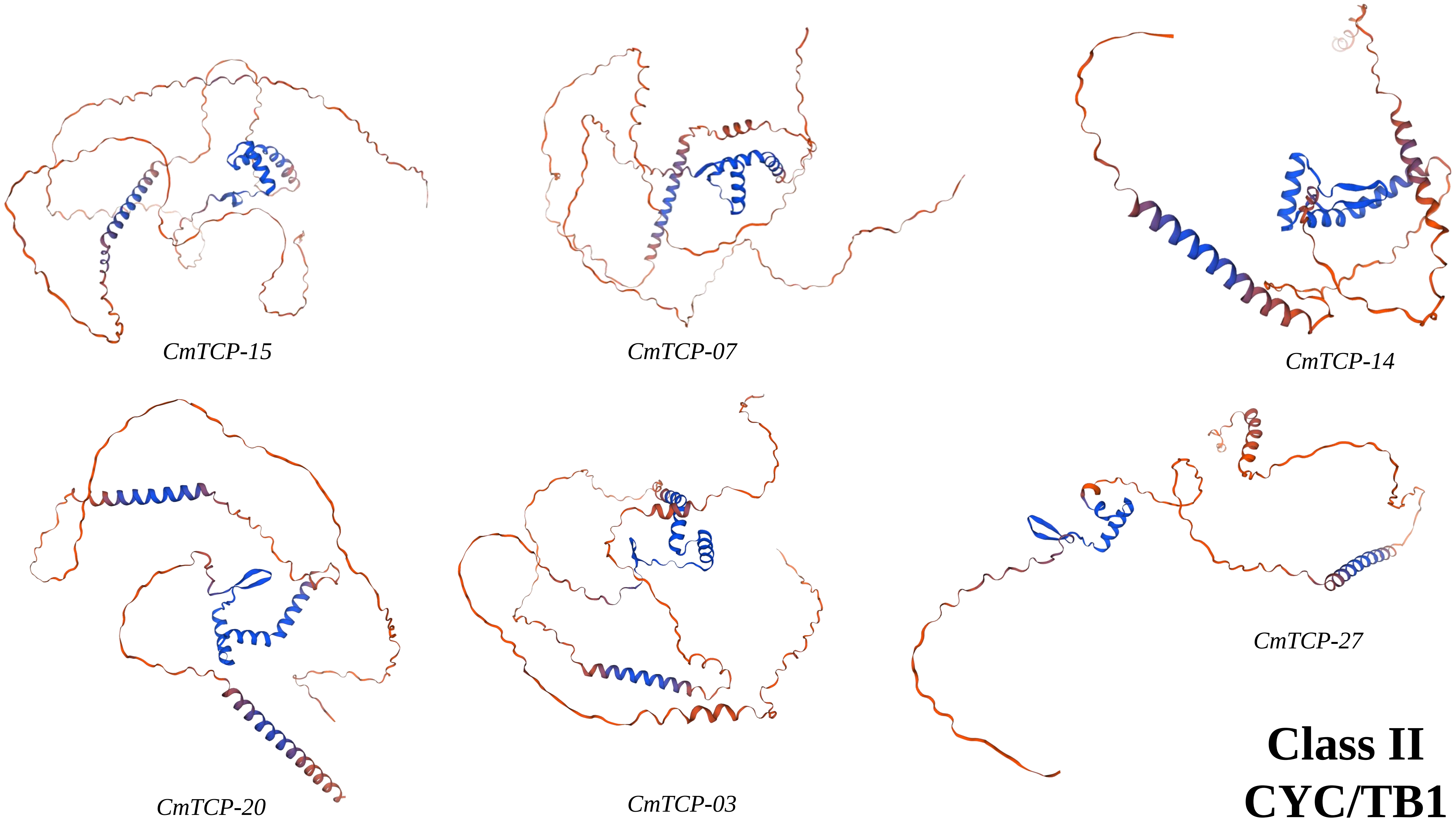
CmTCP-15
CmTCP-07
CmTCP-14
CmTCP-20
CmTCP-03
CmTCP-27
Class II
CYC/TB1

### Protein Homology Modeling using Swiss Model

| Protein | Template | GMQE | Seq Identity |
|---|---|---|---|
| *CmTCP-01* | K7KMJ2.1.A | 0.48 | 77.46% |
| *CmTCP-02* | A0A1S3BX07.1.A | 0.47 | 100 |
| *CmTCP-03* | A0A5D3DXB3.1.A | 0.55 | 100 |
| *CmTCP-04* | 7vp4.2.A | 0.61 | 62.07 |
| *CmTCP-05* | A0A1S3AYR7.1.A | 0.52 | 100 |
| *CmTCP-06* | A0A0A0L891.1.A | 0.56 | 98.85% |
| *CmTCP-07* | A0A1S3CGD1.1.A | 0.52 | 100 |
| *CmTCP-08* | A0A0A0KMY3.1.A | 0.56 | 96.21% |
| *CmTCP-09* | A0A1S3B916.1.A | 0.54 | 100 |
| *CmTCP-10* | A0A5A7UVA7.1.A | 0.36 | 99.16% |
| *CmTCP-11* | A0A0A0KZJ7.1.A | 0.47 | 96.04% |
| *CmTCP-12* | A0A5D3CQK2.1.A | 0.48 | 100 |
| *CmTCP-13* | A0A5D3C791.1.A | 0.63 | 99.03% |
| *CmTCP-14* | A0A0A0KRR2.1.A | 0.61 | 96.39 |
| *CmTCP-15* | A0A0K2RVY1.1.A | 0.54 | 100 |

| Protein | Template | GMQE | Seq Identity |
|---|---|---|---|
| *CmTCP-16* | A0A0A0KMY3.1.A | 0.56 | 95.02% |
| *CmTCP-17* | I1LF87.1.A | 0.57 | 81.86% |
| *CmTCP-18* | A0A5A7SSU2.1.A | 0.47 | 100 |
| *CmTCP-19* | A0A5D3D1W3.1.A | 0.49 | 100 |
| *CmTCP-20* | A0A1S3BFB2.1.A | 0.6 | 100 |
| *CmTCP-21* | A0A5D3CTP8.1.A | 0.53 | 100% |
| *CmTCP-22* | A0A1S3CAQ4.1.A | 0.54 | 100 |
| *CmTCP-23* | A0A0A0KCU1.1.A | 0.54 | 97.85% |
| *CmTCP-24* | A0A0A0KDC7.1.A | 0.63 | 96.24% |
| *CmTCP-25* | A0A0A0LSM3.1.A | 0.52 | 98.32% |
| *CmTCP-26* | A0A1S3BEN2.1.A | 0.65 | 100 |
| *CmTCP-27* | A0A0A0LVQ1.1.A | 0.51 | 95.14% |
| *CmTCP-28* | A0A1S3B7T4.1.A | 0.64 | 100 |
| *CmTCP-29* | A0A1S4DV39.1.A | 0.48 | 100 |
